# CENdetectHOR: a comprehensive tool for CENtromere profiling and HOR detection

**DOI:** 10.1101/2025.01.07.631657

**Authors:** Alessia Daponte, Paolo Pagliuca, Carlos Vila-Verde, Francesco Montinaro, Francesca Antonacci, Mario Ventura, Miguel Ceriani, Claudia R Catacchio

## Abstract

Centromeres are essential for accurate chromosome segregation and are characterized by long arrays of repetitive satellite DNA, showing extensive variation in sequence, length, and organization across species. Despite extensive research, fully characterizing centromeric DNA has been challenging due to its repetitive nature and rapid evolution. Here, we present CENdetectHOR, a computational tool to identify and analyze higher-order repeat (HOR) arrays in centromeric regions across diverse organisms, requiring no *a priori* information. We validate its efficacy using human and arabidopsis genomes, demonstrating its ability to reveal the complexity and diversity of centromeric architectures. CENdetectHOR also recognizes HOR variants, elucidating interindividual and interspecific variations in centromeric regions. These findings establish CENdetectHOR as a powerful, versatile, and rapid tool for advancing research on centromere structure, evolution, and function.

## Background

The faithful segregation of chromosomes into daughter cells during cell division is ensured by the attachment of the spindle apparatus to a multi-protein structure called the kinetochore, which assembles on the primary constriction of each chromosome. The genomic region promoting kinetochore assembly is the centromere, whose essential role in cell division has been largely demonstrated [1]. Chromosome missegregation can lead to a range of biological and pathological consequences, such as aneuploidy, cancer development, embryonic lethality, infertility, and developmental disorders.

As a consequence, since the last century, significant efforts have been made in genomic research to characterize the organization of centromeric DNA in various organisms [2–7]. In the vast majority of multicellular eukaryotes, the centromere is classified as “regional”, meaning it spans a large area of the chromosome, typically ranging from hundreds of kilobases to several megabases of DNA [8]. In these organisms, the centromeric DNA consists of long arrays of satellite sequences that are often composed of tandem repeats of short DNA motifs and can be highly variable in sequence, length, and organization across different species. The total amount of centromeric satellites also varies substantially: in humans and non-human primates, it makes up around 3-5%, whereas it constitutes up to 15-20% of the total genome in *Zea mays* (maize) [9–12].

In addition, the size of the repetitive centromeric unit also varies deeply across species, but it is generally in the range of 150-200 bp [13,14].

While the repetitive nature of centromeric DNA became immediately evident through molecular methods developed in the 1980s—such as restriction enzyme digestion, Southern blotting, and renaturation kinetics assays—the detailed description of species-specific centromeric satellite sequence and organization using sequencing approaches has proven to be extremely challenging due to the inherent repetitive nature of centromeric DNA itself and its rapid evolution. Indeed, the centromeric satellite sequences within a species tend to evolve in concert, where new mutations or changes in the repeat units rapidly spread across different chromosomes, resulting in the homogenization of these repeats within the species [15].

In humans, in particular, four decades of focused studies have demonstrated that the satellite DNA at centromeres, called alpha-satellite, is composed of the repetition of a ca. 171-bp long monomer organized into higher-order repeats (HORs).

A HOR can be defined as an organization of monomers into larger repeat blocks, where corresponding monomers in different blocks share high sequence identity, while adjacent monomers within a block show lower similarity. For example, in a dimeric sequence (in which the repeated unit is made of two monomers) every monomer shares a higher degree of similarity with the ones two steps ahead or behind than with the flanking ones [16,17]. The monomers composing a HOR belong to different families, defined as groups of evolutionarily related monomers. In humans, different units of the same HOR show less than 5% divergence and the degree of divergence between different monomers of the same HOR is 20-40% [18,19]. Very little is known, instead, about the possible presence of HOR arrays and the similarity ranges between monomer families in other species [20].

Untill a few years ago, the centromeric sequences of humans, non-human primates, and other eukaryotes have remained largely absent from reference genomes. The available data primarily come from sequencing of isolated clones or small genomic fragments, typically no larger than a few kilobases, leaving these highly repetitive regions very poorly represented in genome assemblies. This limitation has significantly hindered the comprehensive characterization of entire centromeres and pericentromeric regions, the understanding of the composition of HORs within the same centromere, the identification of active (i.e. able to host the kinetochore) versus inactive centromeric arrays, and, most importantly, the assessment of interindividual and interspecific variability in both qualitative and quantitative aspects. Attempts to characterize centromeric variants from whole-genome sequencing (WGS) data primarily relied on predictions derived from short-read assemblies or k-mer frequency analyses [21,22]. While these methods are effective at identifying the centromeric repeat unit in a genome, they are not able to accurately detect local rearrangements of repeat patterns, changes in direction within arrays, and family composition of HORs.

Recently, advancements in long-read sequencing technologies, combined with increasingly efficient assembly methods, have finally made it possible to fully explore the centromere composition of some genomes. For the first time, a single sequencing read can span an entire centromere, leading to the publication of the first telomere-to-telomere (T2T) reference genomes, which are mostly free of gaps in the centromeric regions [23–25]. The volume of centromeric data now available is remarkable and far too vast to manage through manual inspection. The detailed insights provided for the first time by these new sequencing approaches and the long-standing unanswered desire to explore the genetic variation of centromeric regions have thus sparked the development of numerous tools designed to describe and characterize centromeric structures.

Tools like NTRprism [26] and GRM [27] can successfully locate repetitive regions and provide monomer size and tandem repeat patterns, yet they lack the ability to extract monomeric sequences and phylogenetically analyze them. Alpha-CENTAURI [28] is designed to identify HORs within WGS datasets but needs a reference set for satellite families. StringDecomposer [29] improves sequence extraction but, like NTRprism, GRM, and Alpha-CENTAURI, does not analyze HOR family composition. CentromereArchitect [30] and HORmon [31] point to cluster monomers to infer HORs, but need a centromeric repeat consensus sequence, with HORmon’s predictions which are not based on the consecutive appearance of HOR units, essential to the HOR definition itself. Tools like HumAS-HMMER [32] allow precise alpha-satellite HOR mapping through organism-specific Hidden Markov Models (HMMs) but are currently limited to human genomes.

Here we describe CENdetectHOR, the first fast and efficient tool to comprehensively i) handle centromeric sequences from all possible organisms, ii) requiring no *a priori* information, iii) able to finely characterize HORs of any length, iv) precisely map each HOR in the input sequence, v) describe the monomer composition of each family comprised in the HORs, and vi) infer the phylogenetic relationships between monomers.

## Results

### Overview of CENdetectHOR

CENdetectHOR has been developed to perform an in-depth analysis of regional centromeres and needs as little prior information as the genomic sequence itself. Users may optionally provide a monomeric consensus sequence specific to the organism, enabling CENdetectHOR to define monomers accordingly in terms of start site and thus maintain consistency with regard to previous available data. Since the eventual presence of a consensus sequence does not computationally affect the subsequent steps of the pipeline, in both cases, the workflow (**Figure 1**) will result in the same HOR annotation.

**Figure 1.**
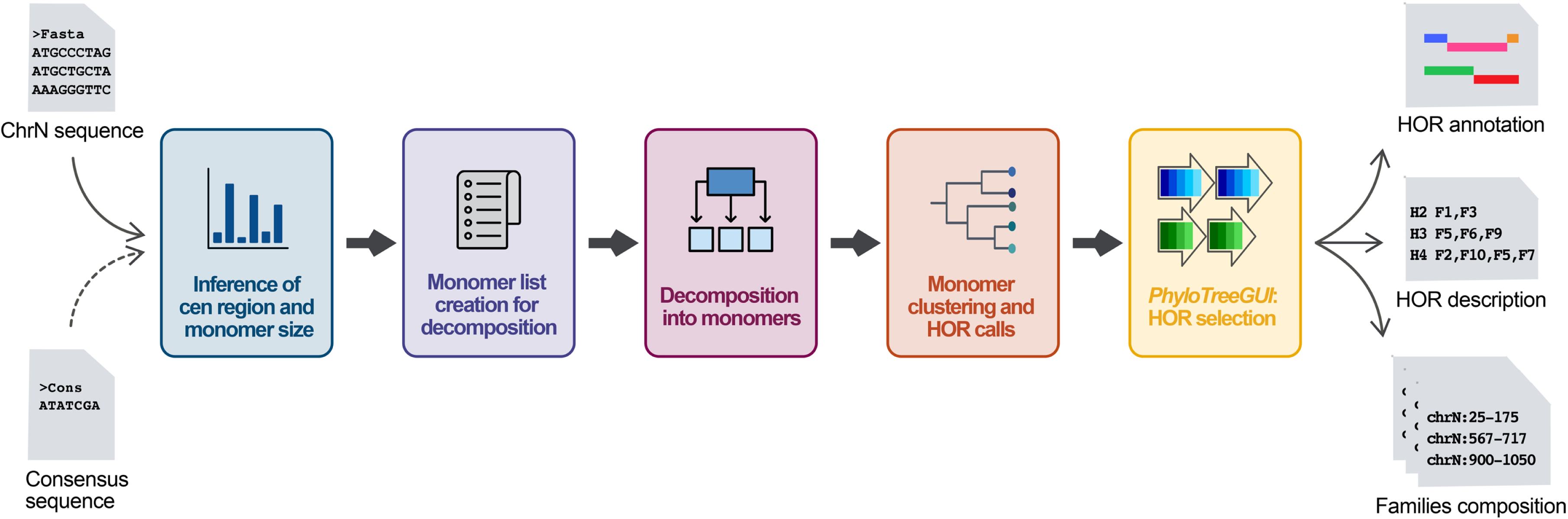
CENdetectHOR workflow. The CENdetectHOR workflow for identifying and annotating Higher-Order Repeats (HORs) in centromeric regions requires an input FASTA file and an optional consensus sequence. If no consensus sequence is provided, CENdetectHOR proceeds with only the FASTA file. Firstly, CENdetectHOR locates the repetitive region within the sequence and generates a histogram to infer the most abundant monomer size (blue frame). Using this inferred monomer size, the tool extracts a set of monomers (purple frame), to decompose the entire sequence into non-overlapping monomers (pink frame). A phylogenetic tree is subsequently constructed for all monomers based on sequence similarities (orange frame). CENdetectHOR uses both the positional data of these monomers and their phylogenetic distances to construct a comprehensive HOR tree that includes all possible HOR configurations. The user can view and filter the resulting HOR tree in *PhyloTreeGUI*, selecting the most appropriate HOR configuration for each centromere (yellow frame). After selection, output files containing HOR annotations, descriptions, and monomer family compositions can be downloaded via *PhyloTreeGUI*.

The process is designed to be executed chromosome by chromosome and begins by scanning each input fasta file to identify highly repetitive regions (see *Methods*). The tool then infers the size of repeated units (i.e., monomers) across these regions and plots this information in a histogram. The monomer length is then used as a filter on the detected repetitive windows to retain the concordant ones: when the consensus sequence is provided, only the windows with the monomer length matching the consensus size or its multiples are kept; otherwise, the putative monomer length is determined as the most abundant genome-wide size. With this information, the whole set of repetitive windows is filtered by keeping only the windows that match the putative monomer length (or its multiples).

Once the centromeric core (i.e. the region of the input sequence composed of tandem repetitions of the centromeric satellite) is identified, the next step relies on StringDecomposer [29] to extract the whole set of non-overlapping monomers from the selected windows that are then clustered based on their similarity. CENdetectHOR uses the clustering information to build a phylogenetic tree of the monomers, which is examined level by level from the root to the leaves to detect HORs. This means that for each region, every inspected level of the monomer tree will give a set of HOR calls, and moving from the root towards the leaves, the distance used to cluster monomers into families decreases. Through this process, HOR detection becomes progressively more refined, allowing the individual families comprising the HOR to be more distinctly identified (**Figure 2**). The result is a phyloXML file [32] that contains both the monomer and the HOR trees.

**Figure 2.**
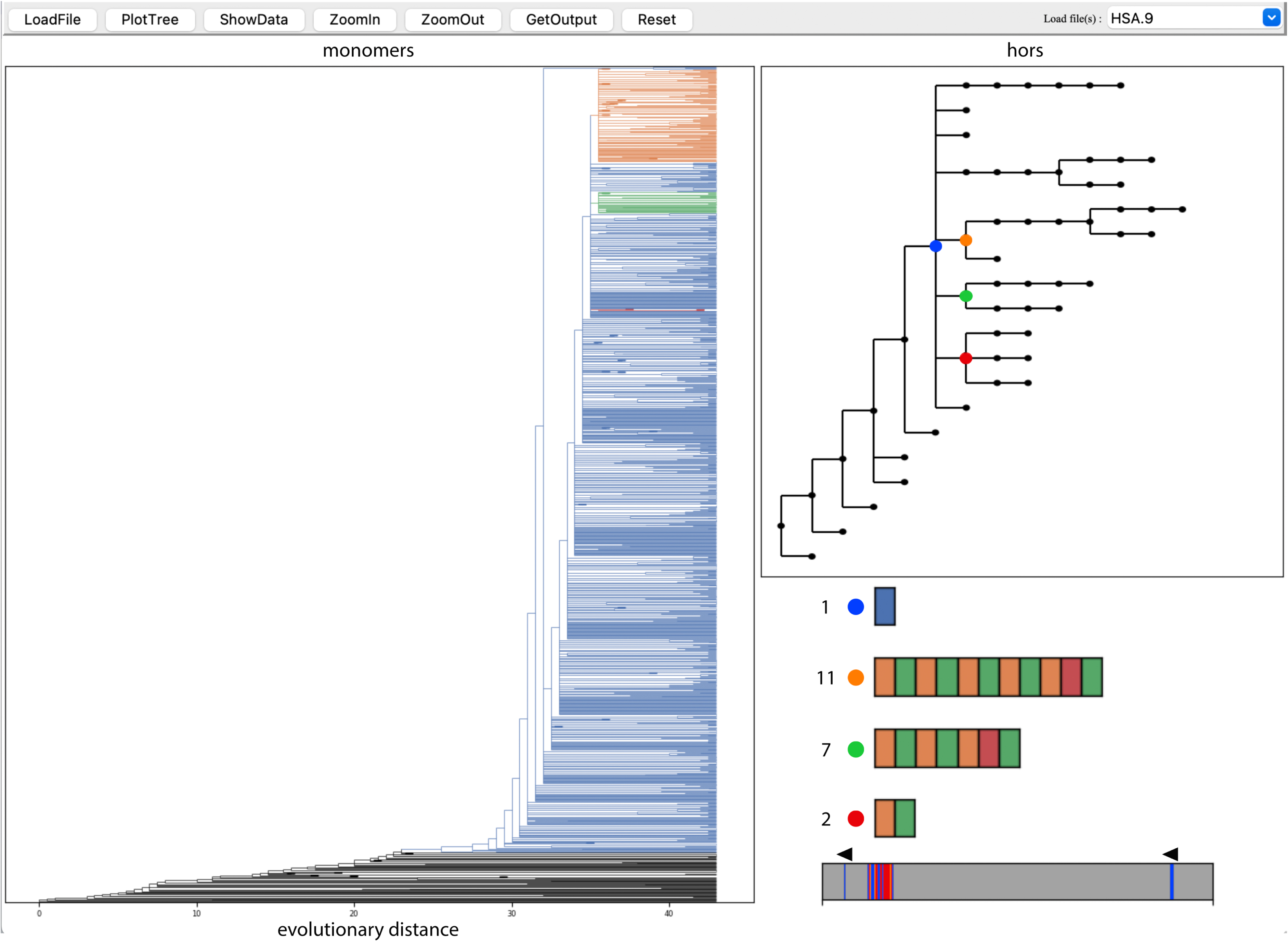
*PhyloTreeGUI* Interface in CENdetectHOR. The *PhyloTreeGUI*, developed for the CENdetectHOR pipeline, provides an interactive visualization and interpretation of HORs generated through monomer clustering within their genomic context. The interface includes three main panels: on the left, the phylogenetic tree of monomers; on the top right, the HOR tree displaying all HOR calls in the investigated centromere; and on the bottom right, a detailed description of the user-selected HOR. Control buttons above the panels allow users to operate the interface. Users can select one or more HORs by clicking on nodes in the HOR tree. Once the HOR becomes colored, in the bottom right panel, details appear, including the number of monomeric units, monomer family composition, and abundance. The corresponding clades in the monomer tree are color-highlighted for easy reference. This interactive tool enables users to customize the analysis, explore specific tree sections, and generate output files.

We benchmarked the clustering process by executing it with an increasing number of monomers, ranging from 5,000 to 33,000 (that is the number of monomers extracted from human chromosome 19, i.e. the maximum number of monomers extracted from a human chromosome). For each monomer count, we ran the process using different numbers of CPUs (from 2 to 16, in increments of 2) and monitored the time and RAM usage, as shown in **Supplementary Table 1**.

A graphical user interface called *PhyloTreeGUI* has been implemented for the CENdetectHOR pipeline to allow for the visualization and analysis of the monomer and the HOR trees. While exploring the output data on the *PhyloTreeGUI*, the user can choose to visualize one or more HORs from the HOR tree: the *PhyloTreeGUI* will show the composition of each selected HOR in terms of monomeric families and their abundance in the analyzed sequence. All the clades involved in the selected HORs will accordingly appear colored in the monomeric tree, which will be displayed next to the HOR one (**Figure 2**). Additionally, the *PhyloTreeGUI* can export the HOR annotation in a bed file, suitable for uploading to the UCSC genome browser [33], along with the genomic coordinates for all the monomers within each selected HOR clade.

CENdetectHOR features a unique naming system, visible in the BED file: each monomer family identifier contains the chromosome specification and a progressive number. For instance, “C2F3” labels the third family found on chromosome 2, numbered sequentially from the root to the leaves of the monomer tree. HORs are instead labeled with the chromosome specification followed by the letter H and the number of families involved in it (e.g. “C2H11” is the identifier of an 11-mer from chromosome 2).

### CENdetectHOR analysis on Human Centromeres

To evaluate the reliability of CENdetectHOR, we tested it on the annotated centromeric regions of the latest human reference genome release (T2T CHM13v2.0/hs1) (**Table 1**). Out of the 415 Mb analyzed (**Supplementary Table 2**), 87 Mb were identified with monomer sizes of 171±3 bp or multiples and retained for further analysis.

**Table 1.** Genomic coordinates analyzed by CENdetectHOR for each human chromosome, detected repetitive regions and extracted monomers. The “Detected repetitive regions with alpha-satellite DNA” column represents the total length of repetitive regions identified for each chromosome, summed from all occurrences listed in Supplementary Table 2.

| Chr | Input sequence coordinates (T2T CHM13v2.0/hs1) | Detected repetitive regions with alpha-satellite DNA (bp) | Monomers n. |
| --- | --- | --- | --- |
| 1 | 116,776,933-147,222,478 | 5.175.000 | 30.464 |
| 2 | 85,999,253-99,662,009 | 2.382.000 | 14.434 |
| 3 | 85,808,316-101,417,699 | 2.577.462 | 15.149 |
| 4 | 44,699,037-59,858,846 | 5.764.000 | 35.092 |
| 5 | 42,095,623-54,537,302 | 4.697.000 | 27.860 |
| 6 | 53,343,452-66,045,458 | 3.295.000 | 19.394 |
| 7 | 55,373,883-68,771,332 | 4.639.000 | 27.243 |
| 8 | 39,155,801-51,298,084 | 2.738.000 | 16.113 |
| 9 | 40,006,720-81,750,116 | 3.028.000 | 17.829 |
| 10 | 34,645,426-46,487,822 | 2.856.000 | 16.801 |
| 11 | 46,090,752-59,405,227 | 4.414.000 | 25.925 |
| 12 | 29,655,629-42,177,514 | 3.382.000 | 19.900 |
| 13 | 71,427-22,451,443 | 4.210.000 | 25.424 |
| 14 | 21,209-17,708,563 | 3.872.000 | 22.777 |
| 15 | 20,914-22,732,017 | 2.591.000 | 15.190 |
| 16 | 30,878,228-57,172,178 | 2.524.000 | 14.846 |
| 17 | 18,957,886-32,526,995 | 4.309.000 | 25.373 |
| 18 | 10,924,744-25,952,596 | 5.566.000 | 32.799 |
| 19 | 19,650,627-34,760,521 | 5.608.000 | 32.969 |
| 20 | 21,417,700-37,935,561 | 4.111.000 | 24.155 |
| 21 | 1-16,344,191 | 1.100.000 | 6.549 |
| 22 | 10,761-20,734,409 | 4.090.000 | 23.965 |
| X | 52,810,456-65,940,305 | 3.597.000 | 21.014 |
| Y | 5,382,483-15,934,471 | 541.000 | 3.264 |
|  | Tot (bp)= 415,029,876 | Tot (bp)= 87,066,462 | Tot (monomers)= 514,529 |

The decomposition of the selected windows resulted in 514,529 monomers, distributed across the 24 chromosomes as detailed in **Table 1**. The clustering process across all possible tree levels (see *Overview of CENdetectHOR*) resulted in 1,748 raw HOR calls. After analysis with *PhyloTreeGUI*, we manually filtered the most representative HOR calls and identified 160 HORs that comprehensively describe all the centromeres (**Table 2**, **Supplementary Table 3, Supplementary Table 4**). Of these, 117 HORs mapped to the active alpha satellite region, based on Altemose et al. [26](**Supplementary Table 3 and 4**).

**Table 2.**
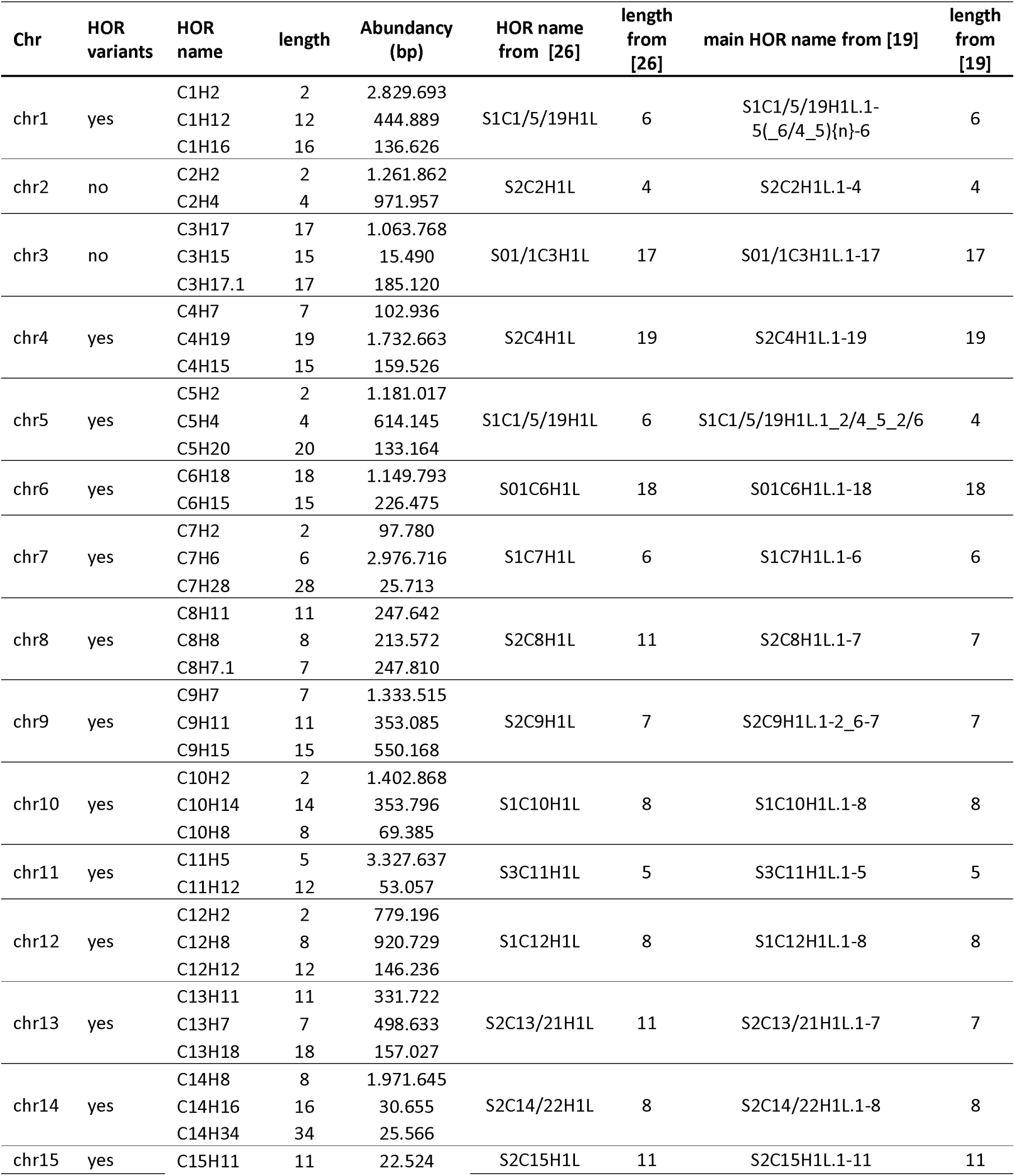
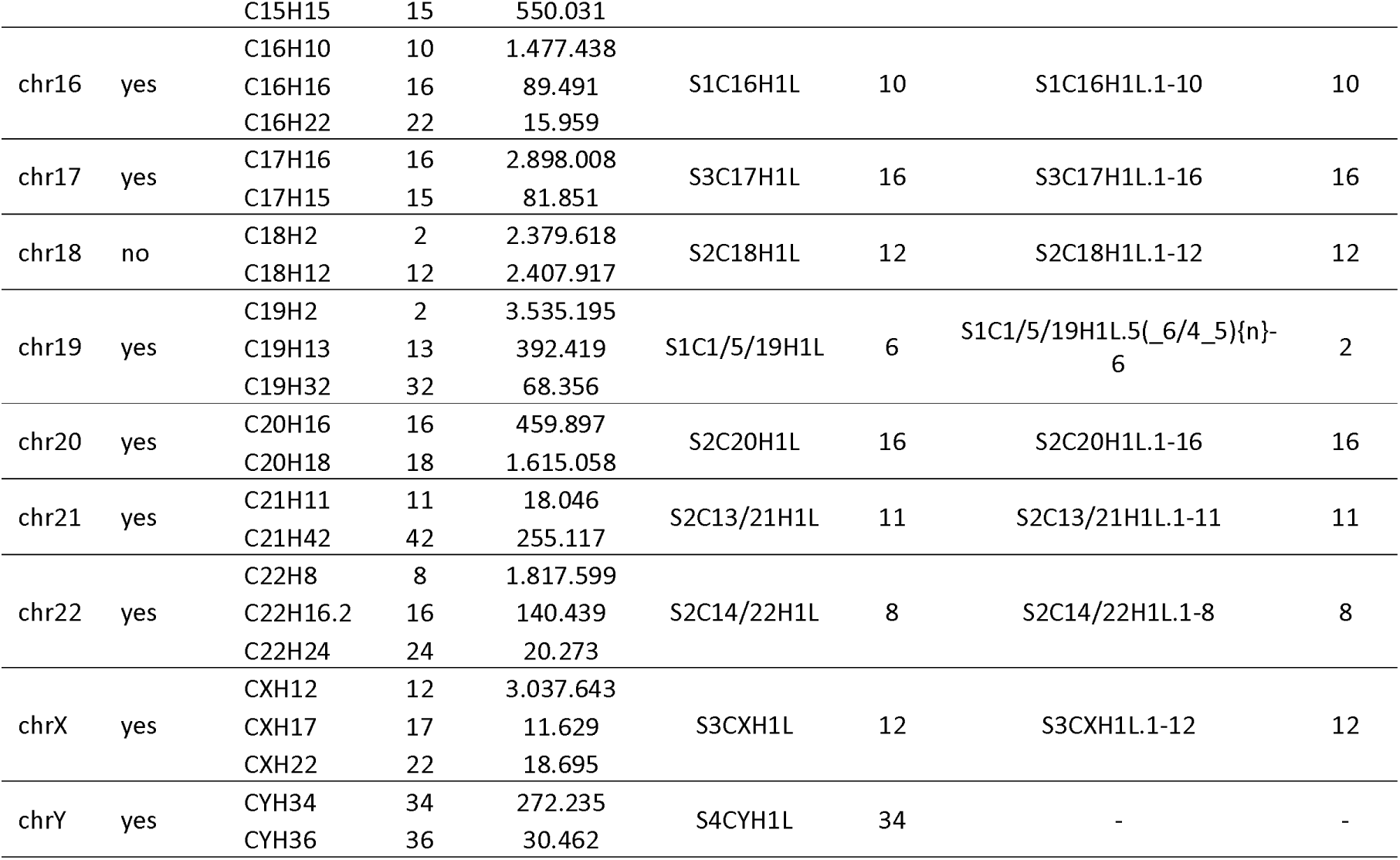
Comparison of the three most abundant active HORs detected by CENdetectHOR for each chromosome with the ones annotated by [26] and [19]. Lengths indicate the number of monomer families involved on the HORs.

The composition of the identified HORs ranges from a minimum of two families (chr1, chr2, chr5, chr7, chr9, chr10, chr12, chr18 and chr19) to a maximum of 42 families on chromosome 21 (**Supplementary Table 3 and 4**). Additionally, CENdetectHOR identified the presence of HOR variants in all the centromeres except for chromosomes 2, 3, and 18 (**Table 2 and Supplementary Figure 1**). For instance, in the active region of chromosome 10, 11 HORs were detected: C10H2(dimer), C10H6 (6mer), C10H8 (8mer), C10H10 (10mer), C10H12 (12mer), C10H12.1 (12mer), C10H14 (14mer), C10H14.1 (14mer), C10H16 (16mer), C10H20 (20mer), and C10H22 (22mer) (**Figure 3A and 3B**). The *PhyloTreeGUI* analysis confirmed that these are indeed variants of C10H8, i.e. HORs sharing a similar family composition and structure. The only difference is the presence or absence of one or a few monomers, or the substitution of monomers with those belonging to a different family. Another example of HOR variants in human centromeres is chromosome 6, where CENdetectHOR identifies only two HORs: C6H18 (18mer) and C6H15 (15mer). The latter differs from the 18mer by the deletion of three monomers (**Figure 3C**).

**Figure 3.**
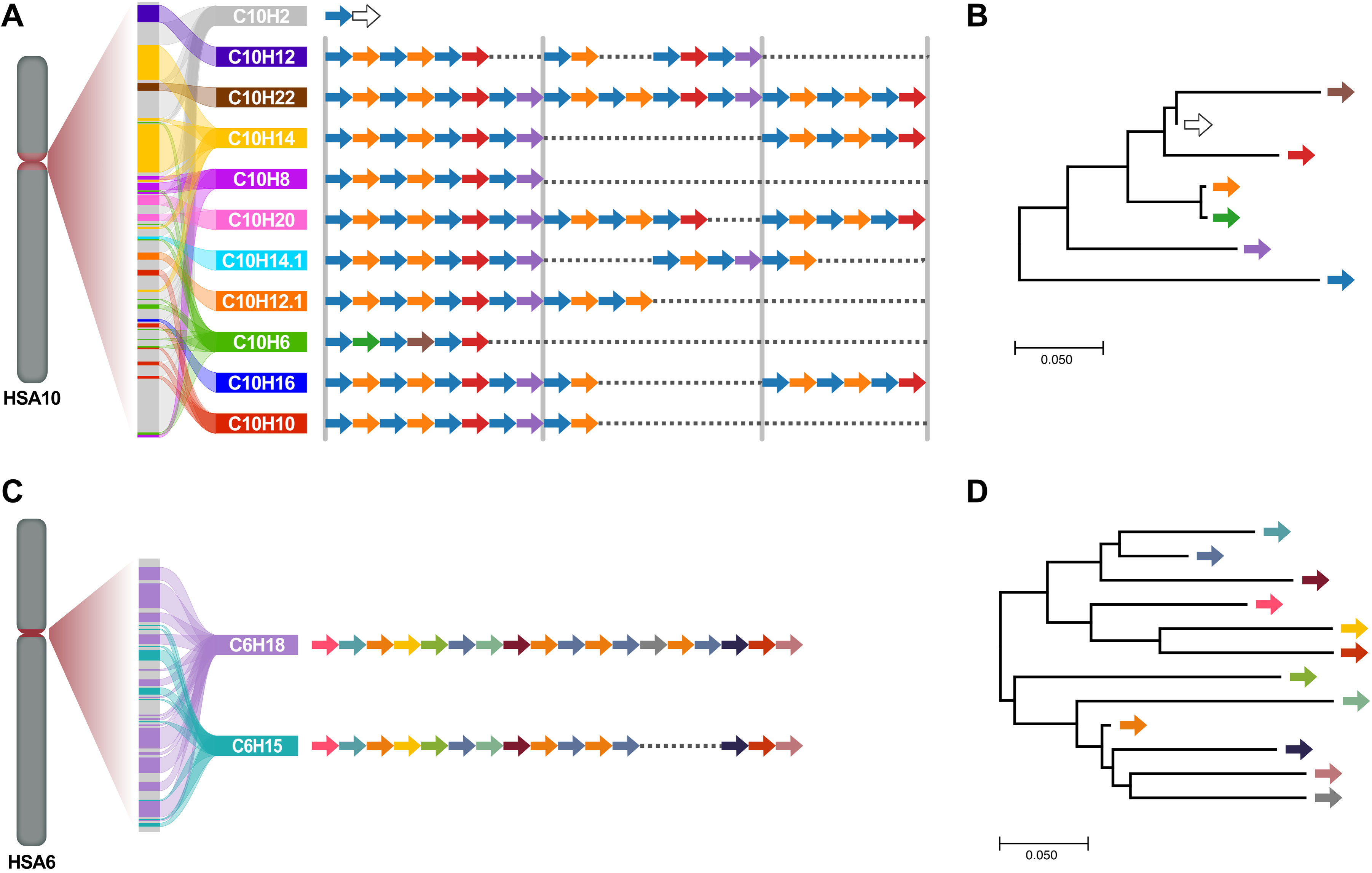
HOR variants annotation by CENdetectHOR. Examples of detailed characterization of HOR variants on chromosomes 10 (panels A and B) and 6 (panels C and D). Panel **A**: For chromosome 10, we identified and localized a dimeric HOR, shown in gray, along with nine active HORs, variants of the 8mer HOR, C10H8; for instance, C10H22 is a 22mer created from an incomplete triplication of the C10H8 8mer (missing the last two monomers). The relative positions of each variant in the centromere are displayed as colored segments on the bar to the left. On the right side, the detailed composition of each variant in terms of families is shown, with arrows of the same color indicating monomers that belong to the same family. Panel **B**: An evolutionary analysis (see *Supplementary Information*) of monomers forming the active HORs in chromosome 10 reveals that the monomer families shown in purple, red, green, and orange are more closely related to each other than to the monomer family in blue. Likely, these families initially formed dimeric arrays, which evolved into the 8mer structure. Panel **C** shows a distinct type of HOR variant detected by CENdetectHOR on chromosome 6, where only two variants are identified. These variants differ by the absence of three monomers in the shorter HOR. Panel **D**: An evolutionary analysis (see *Supplementary Information*) of monomers forming the active HORs in chromosome 6 reveals two distinct monomer groups that alternate to create the two observed HOR variants.

We aimed to determine whether the same monomer families are located on different chromosomes and whether the same HORs are distributed across various chromosomes. To address this question, we generated consensus sequences for each monomer family and constructed a distance matrix to analyze relationships among these consensus sequences (**Supplementary Table 5**). This approach allowed us to quantify the genetic diversity across monomer families and identify potential patterns in their chromosomal distribution. We ultimately obtained a total of 562 consensus sequences, corresponding to 562 distinct monomer families across different chromosomes. We found that nine of them had a perfect match with a family on another chromosome (**Supplementary Table 6**). This reduced the number of different families to 553 (**Additional File 1**).

Based on the accepted criterion that human alpha-satellite families consist of monomers with less than 5% sequence divergence [27,35,36], we used the generated consensus matrix to group monomers within this threshold (**Supplementary Table 7**).

In examining these groups of families, we observed that certain monomer families participate in HORs across different chromosomes. For instance, chromosomes 3 and 12 share two monomer families, while chromosomes 2 and 20 contain three common families, two of which are also present on chromosome 18, and one on chromosome 8. Additionally, one monomer family is shared between chromosomes 4 and 9 and one between chromosomes 9 and 20. Chromosomes 1, 5, and 19 also share two families that form dimeric HORs with chromosome-specific HOR variants.

Having established that monomers from the same family appear across different chromosomes, we investigated the possibility of identical HORs existing on multiple centromeres. Indeed, we identified that two pairs of non-homologous chromosomes (13/21 and 14/22) share near-identical HORs in their active regions, with each pair exhibiting the same monomer families forming the same HOR structure.

### HOR prediction in Arabidopsis thaliana genome

To assess CENdetectHOR performances in a non-human organism, we tested it on the *Arabidopsis thaliana* genome using the first T2T assembly (thus with the centromeric region completely resolved) (Col-CEN_v1.2). From the full genome sequence, CENdetectHOR successfully located the centromere regions on each chromosome: chromosomes 1, 2, 3, and 5 each contain a unique satellite block, while chromosome 4 has two distinct satellite blocks (**Supplementary Table 8**).

Our tool then used the 178bp consensus sequence from Naish et al. [6] to extract all monomers (n=66,037, **Supplementary Table 8**), cluster them into a monomer tree and call HORs. The analysis of the phylogenetic trees using PhyloTreeGUI identified 90 HORs (**Supplementary Table 9**), composed of a total of 100 distinct monomer families. For each family, we generated a consensus sequence (**Additional File 2**).

The only centromere showing a predominant non-monomeric HOR is the centromere of chromosome 4, where we identified a trimeric HOR spanning slightly over 1 Mb, and confirmed it with a dotplot (**Supplementary Figure 2**). Chromosomes 1 and 3 exhibited a similar centromere organization, each featuring two primary monomeric arrays spanning 474 kb and 59 kb on chromosome 1, and 473 kb and 62 kb on chromosome 3. The same monomer families assemble a dimeric HOR of 134 and 70 kb on chromosome 1 and chromosome 3, respectively. Additionally, these two monomer families generate a broader range of HOR variants, ranging from 3-mers to 12-mers, which cover a total of 125 kb on chromosome 1 and 165 kb on chromosome 3 (**Table 3, Supplementary Table 9**).

**Table 3.**
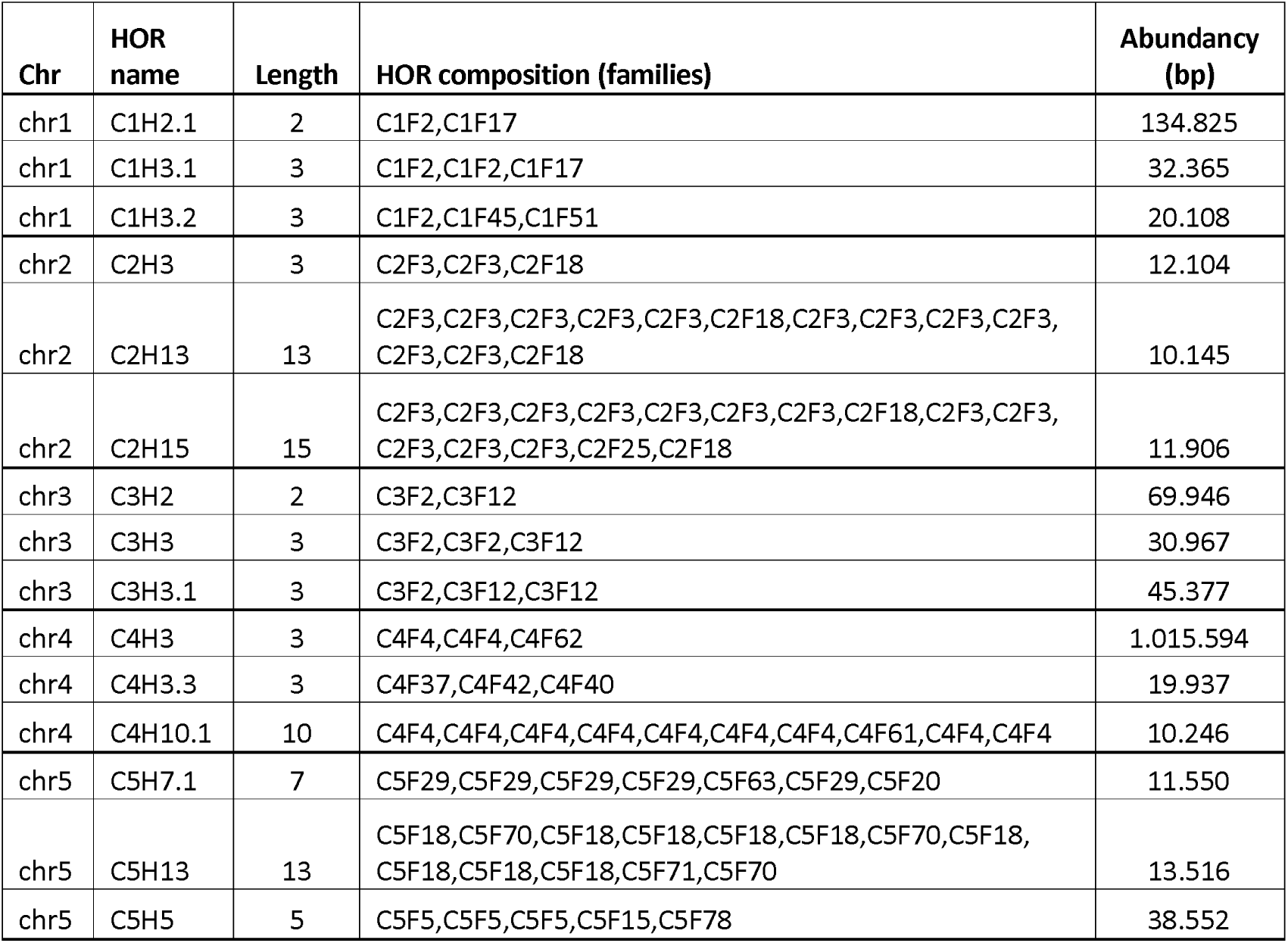
Description of the most abundant HORs detected with CENdetectHOR in the Arabidopsis thaliana genome (Col-CEN_v1.2). Lengths indicate the number of monomer families involved on the HORs.

In contrast, chromosome 2 is dominated by a single monomeric family, forming a monomeric array that spans 1,162 kb of the centromeric sequence. This same family also makes up a 12 kb trimeric HOR, in which two monomers belonging to this family are followed by a monomer belonging to a lesser-represented one. Monomers from these families are also organized in other HORs, reaching up to 15-mers. A third family of monomers forms a separate array, covering 41 kb of the centromeric region. Lastly, chromosome 5 mainly shows a monomeric organization (**Table 3, Supplementary Table 9**). To determine whether HORs are shared between different monomers, we constructed a similarity matrix (**Supplementary Table 10**), and applied a 97.5% threshold for defining monomer family membership [6]. We found that, in only a few cases, monomers from the same family appeared in HORs located on different chromosomes: one family is shared between chromosomes 1 and 5 (C1F2/C5F20 and C1F2/C5F21), another between chromosomes 2 and 4 (C2F3/C4F31), and a third between chromosomes 3 and 4 (C3F2/C4F44). However, we found no evidence of complete HORs being shared across different centromeres.

## Discussion

CENdetectHOR represents a significant advancement in centromere research, offering a unique, fast, and flexible approach for identifying and characterizing HORs across any genome. Unlike recently developed algorithms, which rely on a reference sequence [30,31], prior knowledge of the organism’s centromeric satellite DNA [32], or do not provide information about the family composition of the detected HORs [26,27], CENdetectHOR operates effectively without any prior information beyond a FASTA file with the genomic sequence itself. This independence makes it adaptable to various species, providing researchers with a powerful tool for comparative and evolutionary studies across diverse organisms.

The streamlined workflow of CENdetectHOR, which includes automated identification of repetitive regions, monomer clustering, and HOR annotation, allows it to achieve rapid processing speed without sacrificing accuracy. The method’s flexibility is further enhanced by its modular structure, enabling researchers to adjust parameters and intervene at specific points if necessary. One key parameter is the k-mer size used for identifying highly repeated regions within the input sequence. This is particularly important, as the degree of repetition—both between and within repetitive units—varies across the centromeric sequences of different organisms. In species with sub-repeats in their centromeric satellite monomers, longer k-mers are necessary for accurately identifying the entire centromeric core.

A key innovation of CENdetectHOR lies in its ability to construct a phylogenetic tree of monomers and detect HORs progressively, without relying on predefined distance thresholds for monomer clustering. Our clustering approach ensures that HORs are identified in a consistent and systematic manner, accommodating the varying evolutionary distances among monomers that can occur in different species. By examining the monomer tree from root to leaves, CENdetectHOR allows for increasingly refined HOR detection, providing users with a detailed view of centromere architecture and evolutionary relationships. In this context, a HOR can be defined more precisely as a sequence of different clades of the monomer phylogeny such that it is repeated head-to-tail. The user can then analyze the full picture to define the actual HORs, selecting among increasing levels of characterization. The tree of non-redundant HORs, which constitutes the output of the monomer clustering, can be used to analyze the structure of the chromosome under investigation. To this end, we implemented an interactive tool called *PhyloTreeGUI*: a graphic interface allowing the user to load a phyloXML file containing phylogenetic trees (i.e., monomer tree and tree of non-redundant HORs -see *Supplementary Information*) and perform different actions like selecting one or more HORs, identifying the monomers belonging to the chosen HORs and the corresponding branches in the monomer tree, or examining the portions of the chromosome covered by the selected HORs.

Moreover, the tool’s efficiency in handling large volumes of data—thanks to parallelized processing and memory optimization—makes it suitable for high-throughput analysis of complex centromeric regions, even in large genomes. This is achieved through two key strategies: i) parallelizing the clustering process across multiple threads, which significantly shortens run times, and ii) dividing the matrix into smaller segments for simultaneous computation, which reduces memory usage.

To develop and validate CENdetectHOR, we applied it to the human genome and compared our findings with prior annotations [19,26]. Using our approach, we identified a total of 160 HORs, doubling the 81 identified by Altemose et al. [26] and comparing with the 255 distinct, haplotype-specific α-satellite HORs identified by Logsdon et al. [19]. Moreover, the comparison of the active HORs identified across the 24 human centromeres revealed that, while Altemose and colleagues [26] had annotated a single active HOR per centromere, CENdetectHOR differentiated all HOR variants within these active regions—an achievement previously attained only by Logsdon and colleagues [19], who used HumAS-HMMER [32] which depends on precomputed HMMs. For 23 out of 24 chromosomes, we confirmed the presence of the active HORs annotated in prior studies. Specifically, in 13 of these 23 cases, the most abundant HOR identified by CENdetectHOR matched the annotations of Altemose et al. [26], while in all 23 cases, our results aligned with the annotations reported by Logsdon et al. [19] (**Table 2, Supplementary Table 3 and 4**). Chromosome 5 was the only centromere for which our prediction did not align with the reference annotation: while Altemose et al. and Logsdon et al. identified a 6mer and a 4mer organization, respectively, CENdetectHOR detected a predominant dimeric structure. To validate this finding, we created a dot plot for the region and found no evidence of a 6mer structure in the chromosome 5 centromere (**Supplementary Figure 3A**).

Furthermore, Altemose et al.[26], building on a hypothesis proposed by McNulty and Sullivan [37], identified the same 6mer HOR (S1C1/5/19H1L) on chromosomes 1, 5, and 19. However, subsequent analysis by Kunyavskaya and colleagues [31] questioned this, suggesting that the high divergence across these chromosomes precludes their classification as a single HOR. Contrary, our results show that while a 6mer pattern appears within 4.6 kb and 9.5 kb regions on chromosomes 1 and 19, respectively, the primary structure of these active centromeric regions consists of dimeric HORs (**Supplementary Figures 3B and 3C**). Additionally, we observed that these dimers and their HOR variants share the same monomer families, indicating conserved monomer composition across these variants. Additionally, as previously suggested by Altemose et al.[26] and Alexandrov [38], and in contrast to findings by Kunyavskaya and colleagues [31], our evolutionary analysis of the monomer families forming the HORs identified by CENdetectHOR revealed paired domains on the 13/21 and 14/22 centromeres.

The adaptability and performance of CENdetectHOR were further demonstrated in a non-human genome. When applied to *Arabidopsis thaliana*, CENdetectHOR identified HORs and variants across all centromeres, revealing structural insights that extend beyond previous annotations (**Table 3, Supplementary Table 9)**. Neish et al. [6] identified HORs in *A. thaliana*, reporting an average HOR length of 2.41 monomers. However, their study did not offer precise mapping details or specify the exact composition of these HORs. Our analysis, in contrast, mapped these HORs at high resolution and characterized their composition, thereby enhancing our understanding of centromere structure in *A. thaliana* and revealing previously undetected complexity within these regions and highlighting new structural insights into arabidopsis centromeres. Furthermore, examination of the similarity matrix showed a high degree of intrachromosomal homogenization, which aligns with earlier findings [6]. This strong homogenization within individual chromosomes supports the idea of concerted evolution driving monomer sequence uniformity across centromeres in arabidopsis.

Beyond its adaptability and speed, CENdetectHOR provides output in widely compatible formats, including BED files, which can be integrated with tools like the UCSC Genome Browser [34], enhancing accessibility and allowing researchers to visualize centromeric structures in their genomic context. The *PhyloTreeGUI* interface further enhances usability by providing an interactive platform for exploring HOR trees, selecting specific HOR configurations, and refining results, thus making the tool accessible to users with varying levels of computational expertise.

With CENdetectHOR, we have achieved an unprecedented level of resolution in centromere analysis, enabling precise identification of HOR variants down to the single-nucleotide level. This breakthrough opens new possibilities for comprehensive and large-scale studies of interindividual variations within centromeres, as well as variations between haplotypes. The ability to pinpoint specific HOR variants requiring no a priori information not only enhances our understanding of the structural diversity within centromeres across individuals but also facilitates comparisons of centromeric regions across populations and evolutionary lineages.

## Conclusions

CENdetectHOR represents a substantial advance in the study of centromeric regions, particularly in understanding HOR structures. By combining flexible, species-agnostic HOR detection with the ability to infer monomer family composition and phylogenetic relationships, CENdetectHOR overcomes key limitations in existing tools. Our tests on both human and arabidopsis centromeres demonstrate CENdetectHOR’s capacity to capture the complexity of centromeric organization, offering precise HOR characterization even in genomes with large arrays of repetitive sequences. The tool’s modular framework not only allows for tailored analysis in diverse organisms but also provides transparency in each analytical step, enhancing interpretability and adaptability for varying research needs. CENdetectHOR ability to rapidly and accurately characterize centromeric HORs across diverse organisms enhances our understanding of centromere diversity and supports comparative studies within and between species. CENdetectHOR’s robust performance, coupled with its versatility, positions it as an essential tool for advancing centromere research in a broad range of genetic and evolutionary studies.

## Methods

### CENdetectHOR validation approach

To facilitate code maintenance and reuse and to allow for the simultaneous execution of the process on all the sequences (chromosomes) provided for the analyzed organism, CENdetectHOR has been developed as a Snakemake [39] pipeline, composed of a sequence of modules that are designed to be both conceptually and practically independent of each other. Moreover, each module provides intermediate outputs to allow users to verify the accuracy of each step. All the custom scripts have been coded in Python 3.9 [40].

The above-mentioned modules are here summarized and further described below: 1) identification of highly repetitive regions and repeated unit size; 2) windows filtering and monomer selection; 3) decomposition into monomers; 4) hierarchical monomer clustering; 5) HOR identification; 6) visualization.

To validate CENdetectHOR applicability in all the genomes, we tested the pipeline and the *PhyloTreeGUI* with both human and Arabidopsis genomes. We used the T2T CHM13v2.0/hs1 version of the human genome, providing as input sequences the whole regions annotated as centromeric on the UCSC genome browser [25,26,34,41] (**Table 1**) and the human alpha satellite consensus sequence from NCBI [42](accession number: X07685). For the Arabidopsis thaliana genome, we provided as input the whole genome, using the Col-CEN genome assembly v1.2 and the consensus sequence for the centromeric satellite they provided [6].

To validate the HOR annotation detected with CENdetectHOR, we created dot plots using EMBOSS Dottup [43].

### Identification of highly repetitive regions and repeated unit size

To identify the highly repeated regions of the input sequences, we used an approach similar to the one followed by Naish et al. [6]. For each of the provided sequences, we first split it into non-overlapping windows of 1kb in size to identify, in each window, all the possible k-mers (i.e. all the k-mers extracted using a 1 bp sliding window) and count the non-unique ones. Contiguous windows with at least 10% of repetitive k-mers were merged. The k-mer size can be chosen based on the composition of the specific satellite (see *Discussion*). For the analysis of the human and Arabidopsis thaliana genomes, we used the default value of 8 bp.

Windows annotation are stored, and merged windows longer than 5kbp were further analyzed, estimating and visualising the distance between identical k-mers. The coordinates and the detected repetitive unit size for all the filtered windows are saved to a file.

### Windows filtering and monomer selection

To narrow the subsequent analysis to the centromeric core—defined as the windows (regions) containing the tandem repetition of the centromeric satellite—all the detected windows from the previous step across all chromosomes were merged and filtered. The filtering process selected windows that matched the size of the repetitive unit (or its multiples), allowing for a 4 bp tolerance in the unit’s length.

If the consensus sequence of the centromeric satellite is provided, as in the case of the human genome, the repeat length used to determine the actual monomer size is the length of the consensus sequence itself (i.e., 171 bp for humans). If no consensus is provided, the chosen monomer size is the most frequent one in the selected windows.

Then, a sample of 5 monomeric sequences for each chromosome was randomly selected to be used in the next step.

To prevent sampling the same monomer with different starting positions, the 5 sequences are selected from all monomers that begin with the exact same 6 bp. When analyzing genomes for which the centromeric satellite consensus sequence is annotated, this was accomplished by extracting the first 6bp from the consensus and searching for this pattern across the entire sequence of all selected windows using a 1 bp sliding window. All the sequences beginning with the searched pattern were stored and subsequently filtered based on their length. Only those sequences that matched the specified monomer size were retained.

Since some chromosomes (or part of chromosomes) may have the satellite sequences repeated in the opposite direction, the same process was performed by selecting the first 6 bp of the reverse complement of the consensus sequences as the search pattern. The extracted sequences were then reverse-complemented again to align with the consensus orientation. The pool of sequences with the highest representation from the two resulting sets was chosen as the source for selecting 5 random monomers.

When analyzing genomes without a provided consensus sequence, we extracted all unique 6-mers from the region and identified all monomers starting with each of these 6-mers. The 6-mer that yielded the highest number of corresponding monomers was selected as the primary starting point, and we then extracted 5 random sequences from this set.

### Decomposition into monomers

To obtain the coordinates of all the monomers within the selected windows, we used StringDecomposer v 1.1.2 [29] with the default parameters. StringDecomposer can partition one or more fasta files into contiguous monomers, by aligning the input sequence to a set of monomers given as an additional input file.

We used the complete set (i.e. 5 monomers for each chromosome) of extracted monomers, along with the human alpha-satellite consensus sequence, to perform a parallel decomposition of all human centromeric sequences.

StringDecomposer output was then parsed with a custom Rscript to obtain a bed file with the absolute coordinates of all the monomers falling in the selected windows.

### Hierarchical Monomer Clustering

To build the phylogenetic tree of all the identified monomers, we first computed a triangular matrix of distances between all the monomers by using the Levenshtein distance [44,45] and then clustered the monomers in a phylogenetic tree.

To allow CENdetectHOR to call HORs in inverted regions of the centromeres without classifying them as different ones, for all the monomers extracted with StringDecomposer [29] with a reverted annotation, we first compute the reverse-complement of the sequence, and then computed the distance matrix. The information on the correct orientation of the monomer is kept until the end of the process, visualized in the *PhyloTreeGUI* and saved in the final outputs (in the *strand* field of the bed file).

Given the high resource demands, both in terms of time and memory, for the above-mentioned steps, the process was designed to run in parallel across multiple threads, if available. To achieve this, the triangular distance matrix is computed in separate blocks (with a user-defined size) and merged at the end. Moreover, each matrix block is stored on the disk, allowing for recovery and reloading in case the process is interrupted.

The clustering step has been implemented using the scikit-learn library [46] and is based on Single Linkage agglomerative Clustering [47], in which the distance between two clusters is defined as the minimum distance between each pair of cluster items.

### HOR identification

Given the whole set of identified monomers and the putative phylogenetic tree computed in the previous step, the HOR discovery algorithm searches for HORs by screening the tree multiple times.

As a first step, the monomer tree is visited by levels, from the leaves to the root, to label each identified clade (family) with a different identifier. Once the labeling is completed for all the levels, the algorithm calls HORs visiting again the tree from the root to the leaves and searching for arrays of clades (families) tandemly repeated at least three times (by default). Moreover, at each level, HOR calls are compared with the ones called at the previous step and kept only if they have a longer repeat period, and show a higher intra- or inter-HOR diversity, according to the non-redundancy property (see *Supplementary Information*). Going from root to leaves, the algorithm considers first the most generic version of a HOR.

To simplify the analysis, any overlap of HOR calls is avoided by prioritizing HORs: i) with longer repeat periods, and ii) appearing first in the array (see *Supplementary Information*). Both the HOR tree and the monomer one are then stored in a phyloXML file.

### Interpreting Results Through Visualization

To interactively visualize and interpret the HORs generated through the monomer clustering in their specific genomic context, we implemented the *PhyloTreeGUI*.

It has been developed in Python (version 3.9.x) [40] with *tkinter* (version 8.6) [48] to create the GUI, and *matplotlib* (version 3.8.x), responsible for data plotting [49]. It also requires *Bio*, *Bio.Phylo* and *Bio.Phylo.PhyloXML* (version 1.6.2) libraries [50] to manage phyloXML trees (e.g., I/O operations) and *numpy* (version 1.26.2) [51], *pandas* (version 1.5.1) [52] and *seaborn* (version 0.13.2)[53]. The tool has been successfully tested on both Ubuntu (version 20.04) and macOS (version 14.5).

According to the user selections, the GUI integrates the HOR information with the monomer one. This enables: i) a detailed characterization of HORs regarding the number of monomer families involved, ii) an understanding of the relationships among the monomer families, iii) the identification of the monomer composition of each family that forms all the HORs, and iv) the localization of each HOR within the initial input sequence.

The tool is interactive, enabling users to customize the analysis, explore in detail different parts of the tree and generate output files. These files are structured to comprehensively annotate all the features described above, in a format compatible with the UCSC genome browser [34] (see *Results*).

Noteworthy, while biologically HORs refer to units of at least two monomers, the HOR detection of our algorithm includes cases where the repeated sequence is represented by just one clade (monomeric arrays). This allows the identification and annotation of monomeric regions with consistency and simplicity.

### Comparative analysis of monomer families

Given the genomic locations of all the monomers belonging to each family (which is one of the output files of the *PhyloTreeGUI*), we extracted the relative sequences using a custom script in Python and performed a multiple-sequence alignment using mafft [54] using the iterative refinement method with two cycles only. Consensus sequences for each family were then obtained by using EMBOSS cons (v:6.6.0.0) [55]. The distance matrix of all the consensus sequences was built using the same approach used for the clustering process.

### Availability of data and materials

The codebase of CENdetectHOR, Monomer families and HOR decompositions of human and Arabidopsis thaliana centromeric arrays are available at GitHub (https://github.com/orgs/CENdetectHOR/).

## Competing interests

The authors declare no competing interest.

## Acknowledgments

We would like to thank Dr. Glennis A. Logsdon for her critical reading of the manuscript and valuable suggestions.

## Funding

We acknowledge financial support under the National Recovery and Resilience Plan (NRRP), Mission 4, Component 2, Investment 1.1, Call for tender No. 104 published on 2.2.2022 by the Italian Ministry of University and Research (MUR), funded by the European Union – NextGenerationEU– Project Title *Insights into the fast genome evolution of Gibbons through single-cell strand sequencing and simulation-based approaches* – CUP H53D23001720006-Grant Assignment Decree No. 970 adopted on 30/06/2023 by the Italian Ministry of Ministry of University and Research (MUR) (C.R.C.); and Project Title *Telomere-to-telomere sequencing: the new era of Centromere and neocentromere eVolution (CenVolution)* – CUP H53D23003260006 - Grant Assignment Decree No. 1015 adopted on 07/07/2023 by the Italian Ministry of Ministry of University and Research (MUR) (M.V.).

FM was supported by Fondazione con il Sud (2018-PDR-01136) and by the Italian Ministry of University and Research (MUR) (2022P2ZESR).

This work was also supported by the Italian Ministry of University and Research (MUR) grant PRIN 2020 (project code 2020J84FAM, CUP H93C20000040001) to F.A. and the PNRR project “Fostering Open Science in Social Science Research (FOSSR)” (CUP B83C22003950001) to M.C..

## Authors’ contributions

CENdetectHOR algorithm development, A.D., M.C., P.P., F.M. and C.R.C; CENdetectHOR code development, A.D., M.C. and P.P.; data analyses: A.D., C.V.V., F.A., M.V. and C.R.C.; manuscript draft, A.D., P.P., M.C., F.A. and C.R.C.; editing, all authors; conceptualization, C.R.C. All authors have read and agreed to the published version of the manuscript.

## Figure Legends

**Supplementary Figure 1. HOR variants annotated by CENdetectHOR on chromosomes 8 and 15.** Panel **A** shows the nine HOR variants annotated for chromosome 8, while panel **B** shows the two variants detailed for chromosome 15. The position of each variant is identified in the sequence and represented by a different color. All the variants are then detailed in their composition on the right, with arrows in different colors indicating monomeric families. Chromosome 15 primarily exhibits a 15-mer organization, formed by two repetitions of an array of four monomers followed by an array of seven monomers. In a few regions, an 11-mer variant is observed, where the array of four monomers is repeated only once.

**Supplementary Figure 2. *Arabidopsis thaliana* trimeric HOR.** Dot plot generated using EMBOSS dottup (https://europepmc.org/article/MED/38597606) for the centromeric region of chromosome 4 in *Arabidopsis thaliana*. CENdetectHOR identified a trimeric HOR in this region, which appears in the dot plot as a continuous line after every two dashed lines. The exact coordinates of the plotted region are indicated on the plot axes.

**Supplementary Figure 3. Dot plot graphics for the centromeres of human chromosomes 5, 1 and 19.** Dot plot generated with EMBOSS dottup (https://europepmc.org/article/MED/38597606) for the centromeric regions of chromosomes 5 (A), 1 (B), and 19 (C) in the latest human genome release (T2T CHM13v2.0/hs1). In these regions, CENdetectHOR identified a dimeric HOR, visible in the dot plot as a continuous line alternating with dashed lines. The precise coordinates of each plotted region are shown on the plot axes.

**Additional File 1.** Consensus sequences derived for all the monomer families identified by CENdetectHOR in human centromeres.

**Additional File 2.** Consensus sequences derived for all the monomer families identified by CENdetectHOR in the *Arabidopsis thaliana* centromeres.

## Notes

https://github.com/CENdetectHOR

### Competing Interest Statement

The authors have declared no competing interest.

