## Additional File 1 for "CENdetectHOR: a comprehensive tool for CENtromere profiling and HOR detection"

>C10F106

GAGTTTGCAAGTGGAGATTTCAAGCGATTTGAGGCTAATCTTTGAAATGGAAATATCTTCGTGTAAAAACTACACAGAATCATTCTCAGAAACTGCTTTGTCATCTGTGCGTTCAGTTCACAGAGTTTCACCTTTCTCTTCATAGAGCAGTTTGGAAAGACTCTGTCTGTA

>C10F9

AAGTCTGCAAGTGTTATTTGGACCTCTTTGAGGCCTTCGTTGGAAACGGGATTTCTTCATATAATGCTAGACAGAAGAATTCTCAGTAACTTCTTTGTGTTGTGTGTATTCAACTCACAGAGTTGAACCTTCCTTTAGACAGAGCAGTTTTGAAACACTCTTTTTGTG

>C10F120

GAATTTGCTAGTGTAGATTTCAAACGCTTCGAAGACAGTGATAGAAAAGGATATATCTTCGTATTAAAAGTAGACAAAATCATTCTCAGAAAACTCTTTGTGATGTGTGTGTTCAACTCACAGAGTTTAACCTTTCTTTAATCGAGCAGTTTGGAAATACACTCTTTGTA

>C10F145

GAATTTGCAAGTGGAGATTTCAAGCGATTTGACGCCAATCTTAGACATGGAAATATCTTCATATTAAAAGTACACAGAGTCATTCGTAGAAACTAGTTTGTGATGTGTGCCTTCAACTCACAGAGTTTAACCTTTCTTTTCATAGAGCAGTTGGAAACACTCTATTTGTA

>C10F93

GAATTTGCAAGTGCAGATTTCAAGCGATTCTAGGCCTATGGCAGAAAAGGAAATATCTTCGTATAAAAACTACACAGAATCATTCTCAACAACTACTTTGTGATGTGTGCGTTCAACTCACAGAGTTTAACCTTTCTTTTCATAGAGCAGTTTGGAAACACTCTGTTTGTA

>C10F95

GAATTTGCAAGTGCAGATTTCAAGCGCTTCTAGGCCTATGGCAGAAAAGGAAATATCTTCGTATAAAAACTACACAGAATCATTCTCAACAACTACTTTGTGATGTGTGCGTTCAACTCACAGAGTTTAACCTTTCTTTTCATAGAGCAGTTTGGAAACACTCTGTTTGTA

>C10F85

GAATTTGCAAGTGGAGATTTCAAGCGATTTGAGGCCAATCTTAGAAATGGAAATATCTTCGTATAAAAACTACACAGAATCATTCTCAGAAACTACTTTGTGATGTGTGCGTTCAACTCACAGAGTTTAACCTTTCTTTTCATAGAGCAGTTTGGAAACACTCTGTTTGTA

>C10F56

AAGTCTGCAGTAGGATATTTGGACCTCTTTGAGGCCTTCGTTGGAAACGGGATTTCTTCATATAATGCTAGATAGAAGAATTCTCAGTAACTTGTTTGTGTTGTGTGTATTCAACTAACAGAGTTGAACCTTCCTTTAGAAAGAGCAGTTTTCAAACACTCTGTTTGTG

>C11F29

AATCTGCAAGTGGATATTTGGACCTCTCTGAGGATTTCGTTGGAAACGGGATAAACTTCCCAGAACTACACGGAAGCATTCTGAGAAACTTCTTTGTGATGTTTGCATTCAACTCACAGAGTTGAACCTTGCTTTCATAGTTCAGCTTTCAAACACTCTTTTTGTAG

>C11F50

GATCGGCCAGTGGATATTTGGACCTCTTCGAAGATTTCGTTGGAAATGGGATAAACTTCACATAAAAGCTAAACCGAAGCATTCTCAGAAACTTCTTTGTGATGTTTGCATTCACCTCACAGAGTCGAACTTTCCCTCTGATACAGCACCTTTGAAACGCTCGTTTTCTAG

>C11F53

ATTCTGCAAGTGGATATGTGGACCTCTGTGAAGATTTCGTTGGAAACGGGTTCATCTTCACAGAAAAACTAAACAGAAGCATTCTCAGAAACTGCTTTGTGATGTTTGTGTTCCACTTCAAGAATTGAACTTTCCTCTTGACAGAGCAGCTCTGAAACCCTCTTTTTCTAG

>C11F54

ATTCTGCAAGTGGATATGTGGACCTCTGTGAAGATTTCGTTGGAAACGGGTTCATCTTCACAGAAAAACTAAACAGAAGCATTCTCAGAAACTGCTTTGTGATGTTTGTGTTCCACTTCAGAATTGAACTTTCCTCTTGACAGAGCAGCTCTGAAACCCTCTTTTTCTAG

>C11F68

AATCTGCAAGTGGATATTTGGACCACTTTGTGGCCTTCCTTCGAAACGGGTATATCTTCACATCAAACCTAGACAGAAGCATTCTCAGAATGTTTCCTGTGATGACTGCATTCAACTCACAGAGGTGAACAATCCTGCTGATGGAGCAGTTTTGAAACTCTCTTTCTTTGG

>C11F80

AATCTGCAAGTGGACATGTGGAGCGCTTCCAGGCCTGTGGTGGAAAAGGAAACATCTTCACATAAGAACTAGAGAGAAGCATTGTCAGAAACTTCTTTGTGATGATTGCATTCAACTCACAGAGTTGAAGATTCCGTTTGAAACAGCAGTTTCGAAACACTCTTTCTGTGG

>C11F82

AATCTGTAAGAGGATATGCGGACTTCTTTGAAGATTTCTTTGGAAACGGGAATATCTTCACAGAAAAACTAAACTGAAGCATTCTCACAAACTTCTTTGTGATGTTTGTGTTCGAGTCACACAGTTTAACCTCGCTTTTCACAGAGCGGTTTTGAGACACTCCTTTCGTAG

>C11F86

AATCTGCGATTGGAGATTTGGACTGCTTTGAGGCCTACTGTAGTAAAGGAAATAACTTCATCTAAAAACCAAACGGAAGCATTCACAGACAATTCTTAGTGATCATTGGATTGAACTAACAGAGCTGAACATTCCTTTAGATGGAGCAGTTTCCAAACACACTTTCTGTAG

>C11F9

AATCTGCAAGTGGACATTTGGAGGGCTTTGAGGCCTGTGGTGGAAAAGGAAAATCTTCACATAAAAACTAGATGGAAGCATTCTCAGAAACTACTTTGTGATGATTGCATTCGACTCACAGAGTTGAACATTCCTATAGATAGAGCAGGTTGTAAACAATCTTTTTGTAG

>C12F10

TACAAACAGAGTGTTTCCAAACTGCTCCATCAAAAGAAAGGTTAAACTCTGTGAGTTGAACGCACACATCACAAAGTAGTTTCTGAGAATGATTCTGTCTAGTTTTTATACGAAGATGTTTCCTTTTCTACTTTGGCCTCAAAGCGCTTGAAATCTCCACCTGCAAATT

>C12F12

TACAAACAGAGTGTTTCCAAACTCTCCATCAAAAGAAAGGTTAAACTCTGTGAGTTGAACGCACACATCACAAAGTAGTTTCTGAGAATGATTCTGTCTAGTTTTTATACGAAGATGTTTCCTTTTCTACTTTGGCCTCAAAGCGCTTGAAATCTCCACCTGCAAATT

>C12F51

CACAAAAAGAGTGTTTCACATCTGCTCTGTCTAAAGGAAGTTCAACTCTGTGAGTTGAATACACACACACAAAGAAGTTACTGAGAATTCTTCTGTCTAACATTATATGAAGAAATCCCGTTTCCAACGAAGGCCTCAAAGAGGTCCAAATATCCACTTGCAGACTT

>C12F53

CACAAAAAGAGTGTTTCAAATCTGCTCTTCTAAAGGAAGGTTCAACTCTGTGAGTTGAATACACACACCACAAATAAGTTACTGAGAATTCTTCTGTGTAACATTATATGAGGAAATCCCGTTTCCAACGAAGGCCTCAAAGAGGTCCAAATATCCACTTGCAGACT

>C12F66

CACAAAAAGGGTGTTTAACATCTGCTCTTCTAAAGGAAAGTTCAACTCTATGAGTTGAATACACACAGCACAAAGAAGTTACTGAGACTTCTCCTATCAAACATTATATGAAGAAATCCCGTTTCCAACGAAGGCCTCAAAGAGGTCCAAATATCTGCTTGCAGACTT

>C12F76

CAAAAAGAGGGTTTCACATCTGCTCTGTCTAAAGGACAGTTCACCTCTGTGAGTTGAATAGAGGCAACACAAAGAACTTACTCAGTATTCTTCTTTCTAGCGTTCTATGAAGAAATCCCGTTTCCAACGAAGGCCTCAAAGAGGTCCAAATATCTGCTTGCAGACTT

>C13F30

TACAAAAAGAGTGTTTCAAACCTGCTCTATGAAAGGCCATGTTCATCTCTATGAGTCGAATGGAAATATCCGAAAGAAATTTCTGGGAATGCTGCTGTCTAGTTTTTATACGAATTCCCGCTTCCAACGAAATCCTCAAAGCAATCCAAATATCCACTTGCAGAATC

>C13F34C21F43

TCCAGAAAGAGTGTTTCAAACCTGCTCTATGAAAGGGAATCTTCAACTCTATGAGTTGAATGCAGACATCAGAAAGAAATTTCTGAGAATGCTGCTGTCTACCTTTTATTTGAATTCCCGCTTCCAACGAAATCCTCCAAGCTATCCAAATATCCACTTGCAGATTC

>C13F40

CACAAAAAGAGTGTTTCAAAACTGCTCTATCAATAGAAAGGTTCAACTCTTTTAGTTGAGTACACACATCACAAACAAGTTTCTGAGAATGCTTCTGTCTGGCTTTTATTGGAAGACGTTTCCTTTTCACCAAAGGCATCAAAGCGCTCCAAATGTCCACTTCCAGATTC

>C13F43

CACAAAAAGAGCGTTTCAAAACTGCTCTATCAATAGAAATGTTCAACTCCTTTGGCTGGGTACACACATCACAAACAAGTTTCTGAGAATGCTTCTGTCTAGTTTTTATGGGAAGACATTCCCTTTTTCACCAAAGGCATCAAAGCGCTCCAAATGTCCACTTCCAGACAC

>C13F46C21F62

CACAAAAAGAGTGTTTCAAAACTGCTCTCTATCAATGGCAAAGTTCAACTCTGTTAGTTGAGGACACATATCACCAACAAGTTTCTGAGAATGCTTCTGTCTATTTTTTATGGGAAGATATTTCCTTTTTCACCGTAGGCGTCAAGGCGATCGAAATGTCCACTTCCACAAAC

>C13F49

CCACAAAAATAGAGTTTCAAAGCTGCTCTGTAAAAAGAAAGGTTCCACTCTGTTAGCTGAGTACACACATCACAAACTTGTTTCTCAGAATCCTTCTGTCTCGTTTTTATGGGAAGATATTTACTTTTTCACCGTAGGCATCAAAGCGCTCCAAATGTCCACATCCAGATAC

>C13F61C21F79

TACAAAACGAGTGTATCCACACTGCTCAATCAAAAGAAAATTTCAACTGTGTGAGATGAATGCACACATCAAAATAAATTTCTCCAAAACTTCTGCCTACTTTTTATGGGAAGATATTTCGTTTTTCAACGTAGGCCAAAAGCACTCCAAATATCAATTTGCAGATT

>C13F62

TTCCAAAAGAGTGTTTGAAACGTGCTCAAAGTAAGGGAATGTTCAACTCTGTGACTTGAATGCAGATATCACCAAGTAGTTTCTAATAGTGCTTCTGTCTAGATTTTAGATGATGATATTCCCGTTTCCAACGAAATCGTTAGAGCTATCCAAATATCCACTTACAGTTT

>C13F69

CTACAAAAACAGTGTTTCCAAACTGCTGCATCAAAAGAAAGGTTCAACTCTGTTAGTTGAGGACACACGTCACAAAGAAGTTTGTGAGAATGCTTCTGTCTAGATTTTGTATGACGATATTCCCTTTTCCAACGATATCATTAAAGCAATCTAAATATCCATTTGCAGAAT

>C13F72C21F80

TAAAAAACTACTGTTTCCAAACCGCTCAATCACAGGAAAGGTTTAACTCTGTGAAATGAATGCATCCATCACAGAGAAGTTTCTCAGAATGCTTCCGTCTCGTTTTCATGTGAAGAAGATTCCTTTTCCACCATATTCCTCATGCGCTCCAAATAAACACTTGCAGATTC

>C14F14

CTACAAAAAGAGGTTTCAAACTGCTCATCAAAAGAAAGTTTAACTCTGTGAGTGAAGCACACATCACAAAGAAGTTTCTCAGAATGCTTCTGTTAGTTTTATGTGAAGATATTTCCTTTTCACATAGGCCTCAAAGCCTCAAATTCCACTTGCAGAT

>C14F16

GTATCTGCAAATGGATATTACCAGTGCTTTGAGGCCTATGGTGAAAAAGGAAATATCTTCACATAAAAACAAGGCAGAAGCATTCTGAGAAACTTCTTTTTGATGTCTGCATTCATCTCACAGAGTTGAACCTTTCTTTTGATTGAGCAGTTTTGAAACGCTCTATTTGTA

>C14F18

AATCTGAAAGGGATATTTGGAGCGCTTTGCAGCCTATGGTGAAAAAGGAAATATCTTCACATAAAAGCTAGACAGAAGCATTCTAAGAAAGTGCTTTGTGACGTGTGCATTCATCTCACAGTGTTGAACCTTTCTTTTGATTGAGCAGTTTTGAAACACTCTTATTGTAG

>C14F19

AATCTTCAAGTGGATATTTGGAGGTTTGTGGCCTGTGGTGGAAAAGGAAATATATTCACATAAAAACTAGATAGAAGCATTCTGAGAAACTTCTTTGTGATGTGCTCATTCAACTCACAGAGTTGAACTTTTCTTTTGTTGAGCAGTTTGCAAACAGTCTTTCTGTAG

>C14F30

TACAAAAAGAGTGTTTCAAACCTACTCTGTGAAAGGGAATATTCAACTCTGTGACTTGAATGCACATATCACAAGGAAGTTTCTGAGAATGCTTCTGTCGAGATTTTATATGAAGATATTCCCGTTTCCAACGAAATCCTGAAATCTATCCAAATATCCCCTCGCAGATT

>C14F37

TACAAAAAGAGTGCTTCAAAGCTGCTCTCTGAAACGGAATGTTCAACTCTATGAGTTGAATGCAAACATCACAAAGACGTTTCTGAGAATGCTTCTGTCTAGATTTGATATGAAGATATTCCCGTTTCCAACGAAATCTTCAAATCTATCCAAATGTCCACTTGCAGATTC

>C14F48

TACTACAGAGTGTTTCAAAACTGCTGTACGAAAGGGAATGTTCAACTCTGTGACTTGAATGCACACATCACAAAGAAGTTTCTGAGGATGCTGCTGTCTACTTTTTATACTTAATCCCGTTTCCAACGAAATCCTCCAAGCTATCCAAATATCCACTTGCAGATTC

>C14F49

TACAAAAAGAGTGTTTCAAACCTACTCTGTGAAAGGGAATATTCAACTCTGTGACTTGAATGCACACATCACAAAGAAGTTTCTGAGGATGCTGCTGTCTACTTTTTATACGTAATCCCGTTTCCAACGAAATCCTCCAAGCTATCCAAATATCCACTTGCAGATTC

>C14F50

TATCTGCAAATGGCTATTTGGAGAGCTTTGAGGCCTATGGTGGAAAAGGAAATCTCTTCCCATAAAAACTAGACAGCAGCATTCTGAGAAACTTATTTGTGATCTGTGCATTCATCTCACAGAGTTGAACCTTTCTTTTGATTCAGCAGTTTTGAAACTGTCGTTTTGTAG

>C14F51

CAATCTGCAGAAGGATACTTGTGAGCCGATTGAGGTCTATGGGGTGATAAGAAATATGTTCACATAAAAACTAGATAGAAAGTTTCTGAGAAACTTCTTTGTGATATTTGCTTTTATCTCATAGAGTTGAAACTTTCTTTTTATTGAGCAGTTTGGGAACAGTCTTTTTGTA

>C14F53

AATCTTCAAGTGGATATTTTCAGCGCTTTGAGGCCTATGGTGGAAAAGAAAATATCTTCACATAAAAACTAGTCAGAAGCATTCTGAGAAACTTCTTTGTGACGTGTGCATTCAACTCATGGAGTTCAACCTTTCTTTTGATTCAGCAGTTTGGAAACAGTCTTTTTACAG

>C14F54

TATCTGCAAACGGATATTTGGAGCACTTTCAGGCCTATAGTAGGAAAGGAAATATCTTCACATAAAAACTAGACAGCAAATTACTGAGAAACTTCTTAATGATGTGTGCATTCATCTCACAGAGTTGAAACTTCTTTTGATTGAGCAGTTTGGAAACACTCTTTTAGTAG

>C14F61

GAATATGCAAAGGAATATTTGTGAGCCCATTGATGCCTCTGGGGAAACAGGAAATATCTTCACATAAAAACGAGACAGAATCTTTCTCAGAAACGTCTTGGTGATGTGTGCATTCATCTCACTGAGTTGAACTTTACTTTGATTGAGCAGTTTGGAAACAGTCTTTTCTA

>C14F62

AATCTGCAAAGGAATATTTGTGAGCCCATTGAGGCTTCTGGGGTGATAGGAAATATCTTCACATAAAAACTAGACAGATACTTTCTGAGAAACTATTTTGTCATGTGTGACTTCTACTCACCGGGTTGAAACTTTCTCTTGATTGAGCAGTTTGGAAACAGTCTTTTTGTAG

>C14F63

ATTCTGAAAGTAGATATTTGGAGAGACTTGAGGACTACGGTGGAAAAGGAAATATCTTCACAAAAAAACTAGACAGAAACATTCTGAGAAGCTTCTTTGTGATGTGTGCATCCATCTCAAAGAGTTGAACCTTTCTTTTGATTGAGCATTTTTGAAGCACTCTTTTTGTAG

>C14F75

AATCTGCAAGTGGATATTTGGAGAGTTTGAGGCCACTGGTGGAAAAGCAAATATCTTCACATCAAAACTAGACAGAATCATTATAAGTAATCTCTTTGAGATGCGTGCATTCAACTCACAGAGTTGGACATTTCCTTTGATTGAGCAGTTTGGAAACAGTCTTTTTGCAG

>C14F76

TATCTGtAAATGGATATTTAAGTCTGAGGCCTAGGTGAAAAAGGAAATATCTCATAATCAGATGGAAGCATTCTAGAAACTTCTTTGTGATGTGTGCATTCATCTCACAGACTAAACTTTCTTTTGATTGAGCAGTTTTGAAACACTCTTTTGG

>C14F79

GAGTCTGCAAGTGGATATTTGGAGCACTTTGTGGCCTATAGTGAAAAAGGAAATATCTTCACATAAAAACTAGATAGAAGAATTCTGAGAAACTTCCTTTGAATGGGCGCATTCATCTCACACTGTTGAACTTTTTTTTTGATTGAGCACCTTCTAAACAGTCATTTTGTA

>C14F80

AATCTGCAAATTGATATTTGGAGTGCTTTTGGCCTACGTTGAAAAACGAAATATCTTCCCATAAAAAGTAGGCAGAAGTTTTGGAGAAATTTATTTTGATGTGTGCATTCATCTCACACAGTTGAAATTTTCTTTTGATTGAGCAGTGTGGATACACTCGTTTTGTA

>C14F82

AATCTGCAAGTGGATATTAGGAGTGCATTACGGCCTATAGTGGAAAATGAAATATCTTCACATAAAAACTAGACAGAAACATTATGAGAAACTGCTTTGTGATGCGTGCATTCATCACCAGAGTTGAGTTTCTCTTTTGATTGAACAGTTTTGAAACACTCTTTCTGTAG

>C14F83

GAATATGAAAGGGAATATTTGAGAGCCCATTGAGGCCTCTGGGGAAATAAGAAATATCTTCACCTAAAAACTAGACAAAAACTTTCTGAGAAACACCCTTGTGATGTGTGCATTCATCATACACAGTTGAACTTTCTTTTGATTGAGCAGTTTGGATACAGTCATTTGTA

>C14F84

GTATCTGCAAGTGGATATTTGGAACGCTTTGAGGCCTATAGTGGAAAAGGAAATATCTTCACATAAAAAACTAGAAAGAAGAATTCTGAGAAACTTCCTAGGAAGGTGTaTTTTCGTCTCACACTGTTAAACCCGTCTTTTGATTGAGCAGCTTCGATACAGTCATTTAGtA

>C14F86

GAATCTGCAAGTGTTTATTTGGAGCGCATGAGGAATATGGTGGAAAAGGAATCTTCTTCACATAAAAACGAGACGGAAGCATTCTGAGAAACTTCTCTGTGATGGATGCATTCATTTCACAGAGTTAAACCTTTCCTGTGATTGAGCGGTTTGGAAACAGTAGTTTTTTA

>C14F88

AAACTGCAAGGGGATATTTGGAGCGTTTTGTGGTCTATGGTAGAAAAGGCTATATCTTCACATAAAAATAGAAGCATTCTGAGGAACTTCCTGATGTGTGCATTCATCTCAAAGAGTTGAACTTTTCTTTTGATTGAGCAGCTTTGAAAAACTCTTTCTGCAG

>C14F91

AACAAAAAGTGTTTTTCAGAACTGCTCTATCAAAAGAAAGATCCACCTCTGTTAGCTGAGTTCACACATCACAAACAAGTTTATGAGAATGCTTCTGTCTAGTTTTTATTTGAAGATATTTCCTTTCTCACCATAGAGCTGAAAGCTGTCCTAATGTTCACTTCCAGATAC

>C14F95

AACAAAAAGTGTTTTTCAGAACTGCTCTATCAAAAGAAAGATCCACCTCTGTTAGCTGAGTTCAGACATCACAAACAAGTTTATGAGAATGCTTCTGTCTAGTTTTTATTTGAAGATATTTCCTTTCTCACCATAGACCTGAAAGCTGTCCTAATGTTCACTTCCAGT

>C14F96

AATCTGCCAGTGGATATTTGGAGCGCTCTGTGGCCCATAGTGGAAAAGGAAATATCTTCATAAAAAAAATAAACAGAAGCACTTTGAGAAACTTCTCTGTGTTGTATGCAGTCATATCTCAGACATGAAACTTTCTTTGGTACAGCAGTTTTAAAACACTCTTTTTGGAG

>C15F101

CTACAAAAAGAGTGTTTCCAAACTGCTGTATCAAAACAAAGGTTCAACTCTGTTAGTTGAGGACACACATCACAAATAAGTTTCTGAGAATGCTTCTGTCTAGTTCTTATTTGAAGACATTTCCTTTCTCACCTTAGGCCTGAAAGCGCTCGAAATATCCACTTCCAGATA

>C15F111

CCACAAAAAGAGTGTTTCCAAACTGCTCTGTGAAAAGGAAGGTTCAACTCTGTTAGTTGAGTACACACATCACAAAGAGGTTTCTGAGAATGCTGCTGACTAGTTTTTATTTGAAGATATTTCCCTTTTCACCTTAGGCCTAAGAGTGCTCGAAATGTCCATTTCCACATAC

>C15F118

TACAAAAAGAGTGTTTCAAACCTGCTGTATGAAAGGGAATGTTCAACTCTATGAGTTGAATGCAAACATTACAAAGAAGTTTCTGAGAATGCTTCTGTCTAGATTTTATATGAAGGTTTTCCCGTTTCCAACGAAATTTTCAATGCTCTCAAAATATCCACTTGTAGATT

>C15F129

TCCACAAAGTGTGTTTCAAACGTGCTGTATGAAAGGGAATGTTCAACTCTATGAGTTGAATGCAAACATCACAAAGAAGATTCTGAGAATGCTTTTGTCTAGATTTTATATGAAGATATTCCCGTGTCCAACGAAATTTTCAAAGGTCTCCAAATATCCATTTGTAGATT

>C15F131

TACAGAAAGAGTGTTTCAAACCTGCTCTATGAACGGGAATGTTCAGCTCTGTGAGTTGAATGCAAACATCACAAAGCAGGTTCTGAGAATGCTTCCGTCTAGATTTTAAATGAGGATATTCCCGTTTCCAACGAAATCCTCGAAGCTATCCAAATATCCACTTGCAGATTC

>C15F135

TACGACAGAAACAGTGATTCAAACCTGCTCTATGAAAGGGAATGTTCAACTAGGTGACTTGAATGCAAACATCACAAAGCAGTTTCTGAGAATGCTGCTGTCTACTTTCTATTTGTAATCCCGTTTCCAACGAAATCCTCAGAACTATCGAAATTTCCAATTGCAGATTC

>C15F141

CTATGGAAAGATTGTCTCAAAACTGCTCAATCAAACCAAAGGTTCAACTCTGTGAGATGAATGCCCACATCACAAAGAAGTTTCTCAGAGTACTTCTGTGTAGTTTCTATTTGAGGATAGTTCCTTTTCCACCACAGACCAGAAAGGGCTCCAAATATCCATTGCAGATG

>C15F147

CTACGGAAAGATTGTCTCAAaACTGCTAAATCAAAACAAAGGTTCAACTCTGTGTGATGAATGCATTCATCACAaAGAAGTTTCTCTGAATGCTTCTGTGcAGTTTTTATTTGAAGATAaTTGCTTTTCCAGTATAGGGCGAAATAGGGCTCCAAATATTCACTTGCAGATT

>C15F14

CTACAAAAAGAGGTTTCAAAACTGCTCAATCAAAAGAAAGTTCAACACTGTGAGTGAATGCACACATCACAAAGAAGTTTCTCAGAATGCTTCTGTTAGTTTTTATTGAAGATATTTCCTTTTCCACAATAGGCCTCAAAGCCTCCAAATATCCACTTGCAGATT

>C15F157

GAATCTGCAAGTGGATATTTGGAAAGCTTGAGGCCTATTGTGAAAAAGGAAATATCTTCACATAAAAACTACAGAGAAGCATTCTGAGAAACTTCTTTGTGAGGCATGGATTCAACCCACAGAGTTGGACTTATCATTGAGCAGTTTTGAATCTCTCTTTTTGTC

>C15F158

CTACAAAAGGAATGTTTCCAAAATGCTGTATCCAAACAAAGGTTCAACTCTGTGAATTGAGGGCATACATCACAAAGAAGATTCTGAGAATGCTTCTGTCTAGATTTTATATGAAAATATTCCCGTTTCCAACGAAATCCTCAAAGCTATCCAAATATCCACTTGCAAATG

>C15F163

CCACAAAAAGAGAGATTCAAAACTGCTCAATCACAAGATAGGTTCAACTTGGTAATTGGAAAGCACACATGACAAACAATTTCTGAGAATGTTTCTGTGTAGTTTTTAAGGGAAGATATTTGATTTTCAAATGTAGGCCTCAAATCGCTCCAAATATCCACTTGCATATT

>C15F167

CTCCAAAAGAGTGTTTCAGAATTGCTCAATCAAAGGGAAGGTTCAATTCTGTGTGACCAATGCACTCATCACAAAGAAGTTTGTCTGAATGCTTCTGTGTAGAATTGATTTGAAGATAATTCCTTTTCCACCACAGTCCGCAAAGGGCTAAAAATATCCACTTGCCGATT

>C15F16

CTACAAAAAGAGAGATTCAAACTGCTCAATCAAAAGATAGGTTCAACCTGTGAGTTGAATGCACACATCACAAAGAAGTTTCTCAGAGTGCTTCTGTTAGTTTTTATGTGAAGATATTTCCTTTTCCACAATAGGCCTCAAAGCTTTCCAAATATCCACTTGCAGATT

>C15F172

CTACAAAGAGTGTTTCCAAACTCCTCAATCATAAGATAGGTTCAACTCCGATAGTTGAATGCACACATCACAAAGAAGTTTCTCGGAAAGCTTCTGTGTAGTTTTTGATGAAGATATCTTCTTCTCTAAAACAGAACTCCAAGCCCTCCAAATATTCACTTCAAGATT

>C15F175

CTGCAAAAAGAGAGATACAAAACTGCTCTATCAAAAGATAGATTCGACTCTGTGAGTTGAATGCCAACATCGCAAAGAAGTTTCTCAGAATGCTTCTCTGCAGCTTTTTTGTGAGTATGTTTCGTTTTCCACCATAGGGCGAAATGGGGCTCCAAATATCCACTTGCATTT

>C15F182

TACAAAAAGAGTGTTCCAAAACTGCTCAATCATGAAATAGGATCAACCCTGTGAGATGAATGTACGTATGACAGAGAAGTTTCTCAGAATGCTTCTGTGTAGTTTTTATGCGAAGATATTCGACTTTCCACAGTACGCCTCAAAGTTCTCCAATTATCCACTCGTAGATCC

>C15F186

AATCTGCAAGTTGATACTTGGAGCCCTGTTTCACCCTATAGTGGAAAAGCAAATATCTTCACATAAACAAACCCTACAGAGAAGCATTCAGAGAAAGTCCTTTGTGATGTGTGCATTGAACATGCAGAGTTGACACTATCTTTTGATTGTACAGTTTTGAATACGTCTTTTTGTAG

>C15F191

CAATCTGCAAGTGGATATTTGGAGCCCTTTGCGGCCTATGGTGGAAAAGGAAATATCTTCAAATAAAAACTACACAGAAGCATTCTGAGTCCCTATTTTGGAAAAGAAAATATCTTCACTTAAACAACTACGCAGAAATACTGTGAGAAACTTCTTTGTTATGTGAGCATTCAACTCACAGAGTTGAACCTATCTTTTGATTGAGCAGTTTTGAATCTCTCATTTTGCAG

>C15F37

AAACTGCAAGTGGATATGTAGAGCGATTTGAGGCCTACTGTGGAAAAGCAAATATCTTCACATAACAACTACACAGAAGCACTCCTAGAAACTTCTTTGTGATGTGTGAATTCAACTCACAGAGCTGAACCTATCTTTTGATGGAGTAGCTTAGAATCTCTCTTTTTTTA

>C15F38

GATCTGCAAGTGGATATTTGGCGTGCTTTGAGGCCTATCGTGGAAAAGCAAATAACTTCAGATAAAAACTATACAGAAGCATTCTGAGAAACTTCTTTGTGATGTGTGCATTGATCTCACAGAGTTGAAAGTGTATTTTGATTGAGCAGTTTTGAAACACTCTTTTTGTAG

>C15F39

GAGCCTGCAAGTGGATATTTAGAACGATTTGAGGCCTATTGTGGAAAAGCAAATATCTTCACATAAAAACTACACAGAAGCATTCTGAGAAACTTCTTTGGCATGTGTGCATTCAACTAACAGTGTTGAACGTATCTTTTGATTGAGCAGCTTAGAATCTCTCTTTTTGTA

>C15F43

GAATCTGCAAGTGGATATTTGGAGCCCTTTGCAACCTAGGGTGGAAAAGGAAATACCTTCAAATAAAAACTATATAGAAGCATTCCGTAAAACTTCTTTGTGATGTGTGCATTCGTCTCACAGAGTTGAACCTATCTAATGATTGAGCGGTTTTGAAACACTCATTTTGTAG

>C15F45

AATCTGCAAGTGGAtaTTTGGAGCTCTTTGCACCCTGTGGTGgAAAGGAAATATCTTCATATAAAAACTACAAAGAAGCATTCAGAGAaACTTCTTTGTGATGAATGCATTCCTCACACAGAGTTGAACTTTCTTTTTATTGAGCAGTATTGAAACCTCTTTTTGCA

>C15F52

CTACAAAATGAGAGATTCAAAACTGCTCAATGAAAAGATAGGTTCAACTCTGTGAGTTGAATGCACACCTCCAAAGAAGTTTCTCAGAATGCTTCCGTGTAGTTTTTATGTGAAGATATTTACTTTTCCACAGTTGTCCCAAAGCTCTAAAATGTCCACTTGCAGACC

>C15F54

AATCTGCAAGTGGATATTTGGAGCGCTTTGAGGCCTAATGTGGAAAATCAAATATCTTCACATAAAAACTACACAGAGGCATTCTGAGAAACTTCTTTTTTGTGTGTGCATTCAACTCACATAGTTGAAGTAATCTTTGGATTTAGCTGTTTTGAATCTCCTTTTTGCAG

>C15F55

TGCAAAAAGAGAGATTCAAAACTGCTCAATCAAAAGATAGTTTCTACTCCATTAGCTGAAAGACCACATCACAAAAAAAGTTTCTCAGGATGCTTCTGTGTAGTTTTTATGTGAAGATATTTGGTTTTCCACAGTAGGCCTCAAAGCGCTCCAAATATCCACTCACAGATT

>C15F58

GAATCTGCACGTGGATATTTGGAGCGCTTTGAGACCTAAAGTGGAAAAGCAAATATCTTCACATAAAATCTACATAGAGGCACTCTAAGAAACTTCTTTTTGATGTGTGCATTCACCTCACAGAGCTGAACCGATCCTTCGAGTGACCAGTTTTGAATCTCTCTTTTTATA

>C15F59

GAATCACCAAGTGGATATTTGGAGAGCTTTGGGGCCTGTTTTGGAAAATGAAATATCTTCAAAGTAAAACTACACAGAACCATTCTGAGAAACTTCTTTATGATGTGTGCATTCAACTCTCAGAGTTGAACCTAcCTTATGATTGAGCAATTTGGAAACACTCTTTTTGTA

>C15F62

AATCTGCAAGGGGATATTTGGAGCCCTTTGCGGCCTATGGTGGAAAAGGAAATACCTTCAAATGAAAAGCACACAGAGGCATTCTGAGAAACTTCCTCGTGATTGTGCATTCAACTCACAGAGTTAAACCTATCTTATGATTGACCAGTTTTGGAACACTCTTTTCATAG

>C15F63

GTACAAAAAGTGAGATTCAAAACTGCTCAATCCAAAGGTAGTTTCAACCATGTGATATGAATGCACACAGCACAGAGAATTTTCTCAAAATGCGTCTGTCTAGTTTTTATTTGAAGATATTTCCTTTTCTACCATAGGCCACAAACGTCTCCAAATATCCACATGCAGCTT

>C15F69

AACCTGCAAGTGGATATTGGGAGTACTTTGTGGCCTTCTTTGGAAAAGGGAATATCTTCACATAAAAACTACAAAGAAGCATTCTGAGAAACTTCTTTGTGATGTGTGCATTCATCTCACAGTGTTGGACGTTTCTTTTGATAGGGCAGTTTTGAAACACTCTTTTTCTAG

>C15F70

AGTCTGGAAGTTGATATTTGGAGGGCTTTGAGGTCTATTTCGGAAAAGAAAATATCTTCACTTAAAAACTAGGCAGAAATACTGTGAGAAACTTCTTTGTTATGTGAGCATTCAACTCACAGAGCTGAACCTATCTTTTGATTGAGCAGTTTTGAATCTCTCATTTTGCAG

>C15F71

GTACAAAAAGAGAGATTCAAAACTGGTCACTCAAAAGTTAGGTCCAGCTCTGTGAGCTGAATGCACACATCACAAAGATGTTTCTCAGAAGGTTTCTGTATAGTTTTTATATGAAGATATTGCTTTTCCACAATATGCCTCAAATCTCCCCAATTATCCACTTGCAGATT

>C15F76

CTGCAAAAAGAGAGATTCAAATCTGCTGAATCAAAAGATAGTTTAACTCTGTGACTTCAATGCACACCTCACAAGGGTGTTTCTCAGAAGCTTCTGTGTAGTTTTTATATGAAGATATCTCCTTCTCCAAAAGGTCTCAAAGCTCTCCAAATATTCACTTCCAGATT

>C15F85

AATCTGCGAGTGGATATCTGGAGAACTTGGAGGCCTATTTGGAAAAGGAAATATCTTCACATATAAACTATGCAGAAGCATTTTGAGATTCTTCTTTGTGAGGTGTGCATTCAACTCACAGAGTTGAACTTATCTTTTCCTTGAGCACTTTCATATCTCATTTTCTGTAG

>C15F88

CACAAAAAGAGTGTTTCAAAGCTGCTCTGTAAAAAGAAAGGTTCAACTCTGTTAGTTGAATACACACGTCACAAACAAGTTTCTGAGAATGCTTCTGTCTAGTTTTTATGGGAAGATATTTCCTTTTTCACCGTAGGCCTCAAAGCGCTCCAAATGTCCACTTCCACATAC

>C16F10

GAATTTGTAAGTGGAGAATTCAGCCGCTTTGAGGTCAACGGTAGAAAAGGAAATATCTTCGTATAAAAACTAGACAGAATGATTCTCAGAAACTGTTTTGTGATGTGTGCGTTCAACTCACAGAGTTTAACCTTTCTTTTCAAGAGCAGTTAGGAAACACTCTGTTTGT

>C16F20

GAATTTGCAAGTGGAGATTTCAAGCGCTTCGATGCCAATGGTAGAAAAGGAAATATCTTCGTATAAAAACAAGACAAACTCGTTCCCAGACACTGCGTAGTGATGTGTGTGTTTAACTCACAGAGTTTAACCTTTCTTTTCATACAGCATTCTGGAAACCCTCTGTTTGTA

>C16F27

GAATTTGCAAGTGGAGATTTCAAGCGCTTTGAGGCCAAAAGCAGAAAAGGAAATATTTTCTATAAAAACTCGACAGAATCATTCTCAGAAACTGCTCTGTGATGTGTGCGTTCAACTCACAGAGTTTAACTTTTCTTTTCATTCAGCAGTTTGGAAACACTCTGTTTGTA

>C16F41

AAGTCTGCCAGTGGATATTCAGACCTCTTTGAGGCCTTCGTTGGAAACGGGATTTCTTCATATTATGCTAGACAGAAGATTTCTCAGTAACTTCTTTGTGTTGTGTGTATCAACTCACAGAGTTCAACCTTCCTTTAGACAGAGCAGATTTGAAACACTCTTTTTGTG

>C16F51

AAGTCTGCACGTGGATATTTTGACCTCTTTGAGGCCTTCGTTGGAAACGGGTTTTTTTCATGTAAGGCTAGACAGAAGAAATCTCAGTAACTTCCTTGTGTTGTGTGTATTCAACTGACAGAGTTGAACCTTCCTTTAGACAGAGCAGATTCGAAACACTCTTTTTCTG

>C16F56

AAGTCTGCAAGTGGATATTTGGACCTCTTAGATGCCTTCGTTGGAAACGGGATTTCTTCATATAATGCTAGAGGGAAGAATTCTTAGTAACTTCTTTGTGTTGTGTGTATTCAACTGACAGAGTTGAACCTTCCTTTAGACAGAGCAGATTTGAAAGTCTCTTTTTGTG

>C16F62

AAGTCTGCAAGTGGATATCTTGGCCTCTTAGAGGCCTTCGTTGGAAACGGGTTTTTTCATGTAAGGTTAGACAGAGGAATTCCCAGTAACTTCCTTGTGTTGTGTGCATTCAACTCACAGAGTTGAATGATTCTTTACACAGAGCAGATTTGAGACACTCTTTTGGTG

>C17F106

CATCTGCAAGGGGACATGTAGACCTCTTTGAAGATTTCGTTGGAAACGGAATCATCTTCACATAAAAACTATACAGATGCATTCTCAGGAACTTTTTGGTGATGTTTGTATTCAACTCCCAGAGTTGAACTTTCCTTTGGAAAGAGCAGCTATGAAACACTCTTTTTCTAG

>C17F10

GAATGTGCAAGTGGAGATTTGGAGCGCTTTGAGGCCTATGGTAGTAAAGGGAATAGCTTCATAGAAAAACTAGACAGAAGCATTCTCAGAAAATACTTTGTGATGATTGAGTTTAACTCACAGAGCTGAACATTCCTTTGGATGGAGCAGGTTTGAGACACACTTTTTGTAG

>C17F121

AAACTGCAAGGGGATAATTGCACTCTTTGAGGAGTACCGTAGTAAAGGAAATAACTTCCTATAAAAAGAAGACAGAAGCTTTCTCAGAAAATTCTTTGGGATGATTGAGTTGAACTCACAGAGCTGAGCATTCCTTGCGATGTAGCAGTTTAGAAACACACTTTCTGCAG

>C17F117

AAACTGCAAGTGGATAACTGCACTTCTTTGAGGCCTATCGTAGTAAAGGAAATAACTTCCTATAAAAACAAGACAGAAGCTTTCTCAGAAAATTCTCTGGGATGATTGAGTTGAACTCACAGAGCAGTACTTTCCTTGGGATGGAGTAGTTTCGAAACACACTTTCTGTAG

>C17F135

GATCTGCAAGTGGATATTTGGACCACTCTGTGGCCTTCGTTCGAAACGGGTATATCTTCGCATAAAATCTAGACAGAAGCCTTCTCAGAAACTTCTCTGTGATGATTGCATTCAACTCACAGAGTTGAACCCTCCTATGGATAGAGCAGTGTTGAAACTCTCTTTTTGTGG

>C17F144

GATCTACAAGTGGATATTTGGACCACTCTGTGTCCTTCGTTCGAAACGGGTATATCTTCACATGACATCTAGACAGAAGCATTCTCAGAAGCTTCTCTGTGATGACTGCATTCAACTCACGGAGTTGAACACTCCTTTTGAGAGCGCAGTTTTGAAACTCTCTTTCTGTGG

>C17F151

GATCTGCAAGTGGATATTTGGACCACTCTGTGGCCTTCGTTCGAAACGGGTATATCTTCGCATAAAATCTAGACAGAAGCCTTCTCAGAAACTTCTCTGTGATGATTGCATTCAACTCACAGAGTTGAAGGTTCCTTTTCAAAGAGCAGTTTCCAATCACTCTTTCTGTGG

>C17F17

GAATCTGCAAGTGGAGATATGGACCGCTTTGAGGCCTATGGTAGTAAAGGAAATAGCTTCATATAAAAGCTAGACAGTAGCATTCACAGAAAACTCTTGGTGACGACTGAGTTTAACTCACAGAGCTGAACATTCCTTTGGATGGAGCAGTTTCGAAACACACTATTTGTAG

>C17F31

AATCTACAAGTGGATATTTGGACCTCTCTGAGGATTTCGTTGGAAACGGGATAACTGCACCTAACTAAACGGAAGCATTCTCAGAAACTTCTTGGTGATGTTTGCATTCAAATCCCAGAGTTGAACCTTCCTTTGATAGTTCAGGTTTGAAACACTCTTTTTGTAG

>C17F38

AAACTGCAAGTGGATATTTGGCCTCTCTGAGGATTTCGTTGGAAACGGGATAAACCGCACAGAACTAAAACAGAAGCATTCTCAGAACCTTCTTCGTGATGTTTGCATTCAACTCACAGTGTTGAACCTTTCTTTGATAGTTCAGGTTTGAAACACTCTTTTTGTAG

>C17F53

AATCTGCAAGTGCATATTTGGACCTCTGTGAGGAATTCGTTGGAAACGGGATAATTTCAGCTGACTAAACAGAAGCAGTCTCAGAATCTTCTTTGTGATGTTTGCATTCAAATCCCCGAGTTGAACTTTCCTTTCAAAGTTCACGTTTGAAACACTCTTTTTGCAG

>C17F61

AATCTGCAAGTGGATATTTGGACCTATTTTGAAGATTTCGTTGGAAACGGGAGAATCTTCACAGGAAAGCTAAACAGAAGCATTCTCAGAAACTTCTTTGTGATGCTTGCATTCAACTCACAGAGTTGAACTTTCCTTTCGAGAGAGAAGCTTTGAAACACTCTTTTTCCAG

>C17F68

AATCTGCAAGTGGATATGTGGACCTCTCCGAAGATGTCTTTGGAAACGGGAATATCTTCACATAAAAACTAAACAGAAGCATTCTCAGAAACTTCTCTGTGATGTTTGTGTTCAACTCCCAGAGTTTCACATTGCTTTTCATAGAGTAGTTCTGAAACATGCTTTTCGTAG

>C17F79

TGTCTACAAGTGGACATTTGGAGCGCTTTCAGGCCTGTGGTGGAAAACGAATTATGGTCACATAAAAACTGGAGAGAAGCATTGTCAGAAACTTCTTTGTGATGATTGCATTCAACTCACAGAGTTGAAGGTTCCTTTTCAAAGAGCAGTTTCCAATCACTCTTTCTGTGG

>C17F86

ACTGTCTGCAAGTGGACATTTGGAGAGCTTTCAGGCCTGTGTTGGAAAATGAATTATCGTCACATAGACACTAGAGAGAAGCATTGTCAGGAACTTGTTTGTGATGGTTGCATTCAACTCACAGAGTTGAAGGTTCCTTTTCAACCAGCAGTTTCCAAGCACGCCTTCTGTGG

>C17F91

AATCTGCAAGTGGACATTTGGAGGGCTTTGAGGCCTGTGGTGGAAAAGGAATTATCTTCCCGTAAAAGCTAGATAGAAGCATTCTCAGAAACTACTTTGTGATGATTGCATTCAAGTCACAGAGTTGAACATTCCCTTTGACAGAGCAGTTTGGAAACTCTCTTTGTGTA

>C17F99

AATCTGCAAGTGGACGTTTGGAGGGCTTTGTGGTTTGTGGTGGAAAAGGAAATATCTTCACCTAAATACTAGATAGAAGCATTCTCAGAAACTGCTTTGTGATGATTGCATTCACCTCACAGAGTTGAACATTCCTATTGATAGAGCAGTTTGGAAACACTCTTGTTGTG

>C18F105

GAATCTGCAAGTGGATATTTGGATAGCTCTAACGATTTCGTTGGAAACGGGAATACCTTCATATAAAATCTAGACAGTGGCACTCTCAGAAACTGCTTTGTGATATCTGCATTCAAGCCACAGAGTTGAACATTTCCCTTCCTAAAGCAGGTTTGAAACACTCTTTTTGTC

>C18F110

GAATCTGCAAGTGGATATTTGGATAGCTCTAACGATTTCGTTGGAAACGGGAATACTTTAGTATAAAATCTAGACAGAGGCACTCTCAGAAACTGCTTTGTGATATGTGCATTCAAGTCACAGAGTTGAACATTCCCTTTATTAGAGCAGGTTTGAAACACTCTTTTTGTA

>C18F112

GAATCTGCAAGTGGATATTTGGATGGCTTTGTGGATTTCGTTGGAAACGGGAGTATCTTCATAGAaAACCTAGACAGTAACATTCTCAGAAACTGCTTTGTGATATCTGCATTCACGTCACAGAGTTGAACATTCCCTTTCATAGAGCAGGTTTGAAACACACTTTCTGTA

>C18F115

GAATTTGCAAGTTGATACATGGATAGCCCTAACTATTTCGTTGGAAACGGGAATATCTTCATATAAAACCTAGACAGAAGCACTCTCAGAAACTACTTTGTGATATCTGCATTGATATCAGAGAGTTGAATATTCCCTTTCTAAGGGCAGGCTTGAAAGCGTCTTTTCGTG

>C18F120

GAATCTGCAAGTGGATATTTGGATAGCTCAAGCTATTTCGTTGGAAACGGGAATAGCTTCATATAAACTCTAGACAGAAGCACTCTCAGAAACTACTTTGTGATATCTGTATTCAAGTCACAGAGTTGAATATTCCCTTTCTTAGAGCAGGTTTGAAACCGTCTTTTCGTG

>C18F122

GAATATGCAAGGGGATATTTGGATAGCTCGAAGTATTTCGTTGGAAACGGGAATATCTTCATATAAAATCTAGACAGAAGCACTCTCAGAAACTACTTTGTGATATCTGCATTCAAGTCACAGAGTTGAATATTCCCTTTCTTAGAGCAGGTTTGAAACCGTCTTTTCTTG

>C18F125

GAATCTGCAAGTGGATATTTGGATAACTTTGAAGATTTCGTTGGAAACGGGAATATCTTCATGTAAAATCGAGACAGAAGCATTCTCAGAAACTGCTTTGTGATGTCTGCATTCACGTCACAGAGTTGAACATTCGCTTTCATAGAGCAGGTTTGAAACACTCTTTCTGCA

>C18F129

GAATCTGCAAGAGGATATTTGCATAGCTTTGAGGATTTCGTGGGAAACGGGATTGTCTTCAGGTAAAATCTAGACAGAAGCATTCTCAGAAACTTCTTTGGGATGTTTGCATTCAAGTCACAGAGTAGAACATTCCCTTTGGTAGAGCAGGTTTGAAACACTCTTTTTGTA

>C18F13

GTATCTGGAAGTGGACATTTCGATCGATTTCAGGCCTATGTTGAAAAAGGAAATATCTTAACATAAAAACTAGACAGAAGCATTCTCAGAAACGTCTTTGTGATGTGTGTCCTCAACTAACAGAGTTCAACCTTTCTTATGATACAGCAGTTTGGAAACACTCTTTTTATA

>C18F144

GAATCTGCAGGAGGATATTTGGATAGCTTTGAGGGTTACGTTGGAAACGGGATTACATATACAAAGTAGACAGCAGCATTCTCAGAAGCTTCTTTGTGATGTTTGCGTTTAAGTCACAGAGTTGAACGTTCCCTTTCATAGAGCAGGTTTCAAACCCTCTTTCTGCA

>C18F148

GAATCTGCAGGAGGATATTTGGATAGCTTTGAGGATTTCGTTGGAAACGGGATTACATATACAAAGTAGACAGCAGCATTCTCAGAAGCTGCTTTGTGATGTTTGCTTTTAAGTCACAGAGTTGAACATTCCCTTTCATAGAGCAGGTTTCAAACACTCTTTCTGTA

>C18F149

GAATCTGCAGGTGGATATTTGGATAGCTTTCAGGATTTCGTTGGAAACGGGATTACATATACAAAGTAGACAGTAGCATTCTCAGAAGCTTCTCTGTGATGTTTGCTTTTAAGTCACAGAGTTGAGCATTCCCTTTCATAGAGCAGGTTTGAAACACTCTTTCTGTA

>C18F150

GAATCTGCAGATGGATATTCGGATAGCTCTGAGGATTTCGTTGGAGACGGGAATACATAAAGAAAGTAGACAGCAGCATTCTCAGGAGATTCTTTGTGATGTTTGCTTTTAAGTCACAGAGTTGAATATTCCCTTCAATAGAGCAGGTTTGAAACACTCTTTCTGTA

>C18F154

GAATCTGCAAGTGGCTATTTGGCTAGATTTGAGGATTTCGTTGGAAACGGGATTACATATAAAAAGCAGACAGCAGCATTCTCAGAAAGTTCTTTGTGATGATTGCATTCAAGTCACAGAATTGAACATTCCCTTTCACAGAGCAGGTTTGAAACACTCTTTTTGTA

>C18F162

GAATCTGCAAGTGGATATTTGGCTAGTTTTGAGGATTTCGTTGGAAGCGGGAATTCATACAAATTGCAGACTGCAGCGTTCTGAGAAACATCTTTGTGATGTTTGTATTCAGGACACAGAGTTGAACATTCCCTATCATAGAGCAGGTTGGAATCACTCCTTTTGTA

>C18F172

GAATCTGCAAGTGGATATTTGGCTAGCTTTGGGGATTTCGCTGGAAGCGGGAATACATATAAAAAGCACACAGCAGCGTTCTGAGAAACTGCTTTCTGATGTTTGCATTCAAGTCAAAAGTTGAACACTCCCTTTCATAGAGCAGTCTGAAACACCCCTTTTGTA

>C18F17

GTATCTGGAAGTGGACATTTCGAGGGCTTTCAGGCCTATGGTGAAAAAGGAAATATCTTCCCATAAAAACTAGACAGAAGCATTCTCAGAAACTTATTTGTGATGTGTGTCCTCAACTAACAGAGTTGAACCTTTCTTTTGATACAGCAGTTTGGAAACACTCTTTTTGTA

>C18F185

TATCTGCTAGTGGATATTTGGAGAGCTTTAAGGATTTCATTGGAAACCGGAATATCTTCAGGTAAAATCTAGACAGAGGCATTCTCAGAAACTTCTTTGTAATGTGTGTCCTCAACTAACAGTGTACAACCTATCTTTTGATACAGCACGTTGGAAACACTCTTTTTATA

>C18F187

AATCTGCAAGTGGATATTTGGAAAGCTTTAAGGATTTCATTGGAAACCGGAATATCTTCAGGTAAAATCTAGACAgaGGCATTCTCAGAAACTTCTTTGTGATGTGTGTCCTCAAGTAACAGAGTACAACCTGTCTTTTGATACAGCAGTTTGGAAACACTCTTTCTGTA

>C18F18

GTATCTGGAAGTGGACATTTGGAGCGCTTTGACGCCTTTGCTGAAAAAGGAAATATCTTCTCTTCAAAACTAGACAGAAGCATTCCCAGAAACTTCTTTGTGATGTGTGTCCTCAACTAACAGAGTTCAACCTCTCTTATGATACAGAAGTTTGGAAACACTCTTTTTGTA

>C18F190

GAATCTTCAAGTGGACATTTGGAGGGCTTCACTGCCTGTGGTAGAAAAGGAACTTTCTTCACATAAAATCTAGACAGCAGCTATCTGAGAAACTTCTTTGTGATGCCTGCATTCATTTCACAGAATTGAACATTTCTTTTGACTGAGCAGTTTGGAAACACCCTTTCTTTA

>C18F20

GTATCTGGAAGTGGACATTTCGAGCGCTTTCAGGCCTATGGTGAACAAGGAAATATCTTCCCATGCAAACTAGACAGAAGCATTCGCAGAAACTTGTTTGTGATGTGTGTCCTCAACTCACAGAGTTGAACATTTCGTTTGACAGAGCAGTTTGGAAACACGATTTTTGTA

>C18F25

GTATATGGAAGTGGACATTTGGAGCGCTTTCAGGCCTACGTTGGAAAAGGAAATATCTTCCCATAACAACTAGACAGAAGCATTCTCAGAAACTAGTTTCTGATGTGTGTCCTCAACTAACACAGTTGAACATTTCTTTAGACAGAACAGTTTTGAAACACTCTTTTTGTG

>C18F34

GTATCTGGAAGTGGACATTTGGAGCGCTTTCAGGCCTATGTTGAAAAAGGAAATATCTTCCCATAACAACTAGACACAAGCATTCTCAGAAACTTGTTTGTGATGTGTGCCCTCTACTGACAGAGTTGAACCTTTCTTTTCATAGAGCAGTTTTGAAACACTCTTTTTGTA

>C18F45

GTATCTGGAAGAGGACATTTCGAGCGCTTTCAGGCCTATGGTGAACAAGGAAATATCTTCCCATACAAACTTGACAGAAGCATTCTCACAAACTGGTTTGGGATGTATGTCCTCAGCTAACAGAGTACAACCTGTCTTTTGATACAGCAGTATTGAAACACTCTTTCTGTA

>C18F49

GTATCTGGATGTGGACACTTGGAGCGCTTTGACGCTTACGGTGAAAAAGGAAATATCTTCCCATAAAAACTAGACAGAAGCATTCTCACAAACTGGTTTGTGATGTATGTCCTCAACTAACAGAGTTGAACCTTTCTATTTACAGAGCAGTTTTGAAAGACTCTATTGGAG

>C18F4

GTATCTGGAAGTGGACATTTGGAGCGCTTTCAGGCCTATGTTGGAAAAGGAAATATCTTCCCATAACAACTAGACAGAAGCATTCTCAGAAACTTTTTGTGATGTGTGTCCTCAACTAACAGAGTTGAACCTTTCTTTTGATAGAGCAGTTTTGAAACACTCTTTTTGTG

>C18F50

GTATCTGGATGTGGGCACTTGGAGCGCTTGGACGCTTATGGTGAAAAAGGAAATATCGTCCCATAAAAACTAGACAGAAGCATTCTCACAAACTGCTTTGTGACGTATGTCTTCAACTAACAGAGTTGAACATTTCTATTCACAGAGCAGTTTTGAAAGACTCTTTTGGAG

>C18F56

GTGTGTGTAAGTGGACATTTGGAGCACTTTCCGGCCTAAGGTGAAAAAGGAAATATCTTCCCATAAAAACTAGACAGAAGCATTCTCAGAAACTTACTCGTGATGTGTGTCCTCAACTAAAGGAGTAGAACCTTTCTTTTCATAGAGAAGTTTTGAAACGCTCTTTTTGTG

>C18F69

GTATCTGGAAGTGGACATTTGGAGCACTTTGACGCCTTTGGTGAAAAAGGAAATGTCTTCCCATCAAAACTAGACAGAAGCATTCTAAGAAACATTTTTGGGATATATGTACTCAACTAACAGAGTTGAACCTTTCTCTTTATAGATCAGTTTTGGAAAGCTCTTTATGTG

>C18F73

GTATCTGGAACTGGACTTTTGGAGCGATTTCAGGGCTAAGGTGAAAAAGGAAATATCTTCCCATAAAAACTGGACAGAAGCATTCTCAGAAACTTGTTTATGCTGTATCTACTCAACTAACAAAGTTGAACCTTTCTTTTGATAGAGCAGTTTTGAAATGGTCTTTTTGTG

>C18F84

GTATCTGGAAGTGGACATTTGGAGCGCTTTCAGGCCTATGTTGGAAAGGGAAATATCTTCCCGTAACAACTAGGCAGAAGCATTCTCAGAAACTTATTTGAGATGTGTGTACTCAACTAAGAGAATTGAACCACCGTTTTGAAGGAGCAGTTTTGAAACACTCTTTTTCTG

>C18F98

GAATCTGCAAGTGGATATTTGGCTAGCTTTGAGGATTTCGTTGGAAACGGGATTCATATAAAATGCAGACAGCAGCATTCTCAGAAACTTCTTTGTGATGTTTGCATTCAAGTCACAGAGTTGAACATTCCCTTTCATAGAGCAGGTTTGAAACACTCTTTTTGTA

>C19F105

AATTGCAGGTGAATCTTTGGAGCGCTTTGAAGCCTTTGTTGGAAATGGGAATATCTTCACACACAAACTAGCCAGAAGCATTCTCAGAAACTTCTTTGTGATGTGTGCGTTGAACCCAGAGAGATGAACCTTTCCTTTGATAGAGCAGTTTTGAAACGTGTTTTTGTAAG

>C19F111

CTACAAAAAGCGTGTTTCCACACTGCTCTATCAAAAGAAAGTTTCAAGTCTGTGAGTTGAATTGCACACATCCCAGTGAACTTTCGGAAAATGCATGGGTCTACTTTTCATGTGAAGATACCCGTTTCCAACGAATTCTTCAAAGAGTTCCAAGTATCCACAAGCAGATT

>C19F112

CTACAAGAAGTGTGTTTCAACACTGCTCTATCAAAAGAAAGTTTCCAGTCTGTGAGTTGAATGCACACATCACAACGAACTTTCTGAGAATGCTTGGGTCTACTTTTTATGTGAAGATACCCGTTTCCAACGAATAACTCAAAGAGTTCCAAATATACACAGTCAGGTA

>C19F113

AATCCACAATTGGATAATTGGAACGCTTTGATGCCCATGGTAGAAAAGGAAATATCCTCATATAAAAACTAGACAGAAGGATTCACAGAAAATGCTTTGTGATGTGTGCATTCAAATCACGGAGTTGAATCTTTCTTTTGTCAGAGCAGTTTTGAAACACTGTTTCTGTG

>C19F122

AAGTCTGCAAGTGGATATTCAGACTCCTTGAGGCCTTCGTTGGAAACGGGATTTCTTCATATTCTGCTAGACAGAAGAATTCTCAGTAACTTCCTTGTGTTGTGTGTATTCAACTCACAGAGTTGAACGATCCTTTACACAGAGCAGACTTGAAACACTCTTTTTGTG

>C19F135

AAGTCTGCAAGTGGATATTCAGACATCTTTGAGGTTTTCGTAGGAAACGGGATTTCTTCATATTCTGCTAGACAGAAGAATTCTCAGAAACTTCCTTGTGTTGTGTTTATTCAACTCACAGAGTCGAACGATCCTTTACTCAGAGCAGACTTGAAACACTCCATTTGTG

>C19F140

AAGTCTGCACGTGGATATTTTGACCACTTAGAGGCCTTCGTTGGAAACGGGTTTTTTTCATGTAAGGCTAGACAGAAGAATTCCCAGTAACTTCCTTGTGTTGTGTGCATTCAACTCACAGAGATGAACGTTCCCTTAGACAGAGCAGATTTGAAACACTCTATTTGTG

>C19F143

AATCTGCATGTGGATATCTGGAGCGGTTTGAGGCCTACGGTCAAAAAGGAAATATCTTCCTGGGAAAAATAGACGAAAGCATTCTCAGAAACTGCTTTGTGATATGTGCATTCGACTCACCGAGTTGAAACTTTTTTTTGATAGAGCAGTTTTGAAACACTCTGTAG

>C19F149

TACAAAAGCAGTGTTTTGAAACTGCTCTGTCAATAGAAAGGTTCAACTCTGTaAATTGAGCACACACATCCAAAGGAGTTTCTGAGAATCCTTCTGTCTAGTTTTTATTTGAAGATATTTCTTTTTCCACCGTAGGCAACAAAGCGCTCCAAAGGAACTCTTGCGGATT

>C19F150

CTACAAAAATACCGTTTCAACACTGCTCTCTCCAATGGAAGGTTCAACTCTGTGAGTTGAATGCACACATCACAAAGCAGTTTCTGAGAATGCTTCTGTCTAGTCTAGTTAGAATGTGAAGATGTCCCGTTTACAACGAATtCCTCCGAGAGCTCTAAATATCTGGAAGCAGATT

>C19F151

CTACAAAAAGTGTGTTTCAACACCGCTCTATCCAAAGAAAGGTTCGAGTCCGTGAGTTGAACGCACACAACAGAAAGCAGTTTCTGAGGATGCTTCCGTCTGGTTTGTATGTGAAGTCATTTCCTTTTCCATCTAAATCGCCTCAAATCGCTAAAAGATCCACTCGCAGATAC

>C19F153

CTCATAAAAGACGGCCTCAAAACCGCTCTATCAAAAGAAGGGTTCCACTCCAAGAGGTGAATTCACATATCACGGAGAAGTTTCTGAGAATGCTTCTGTCCAATTTTTATGTGAAGATtTTCCCTTTTCCATCATAGGACTCAAGTCGCTCTAAATATCCAATTTCAGATA

>C19F155

TACAAGAAGACTGTTTCAAAACCGCTCTCTCAAAAGGAAGGTTCGACTCCGTGAGTTGAATGCACACGTTACGAAGCAGTTTCTGAGAATGCTTCTGTCTATTTTTCAGGTGAAGATATCACTTTTTCCAACATACGTAAAAAAGTACTCGAAATGAACACTTGCAGATTC

>C19F158

TACAAAAAGGATGTTTCAACACTGCTCTATCAAAAGAAAGGTTCAACGATGTGAATTGAACACACACATCACAAAGGAGTTTCAGAGAATGCTTCTGTCTAGTGTTTAGGTGACGATATTACCTTTTCCCACATAGGCAACAAAGCGCTCCAAATGAATACTTGTGGGTT

>C19F159

CCACAAAAGCAGTGCTTCAAACCTGCTCTaTCAAAGGAAAGGTTCAGCTCGGGGAATTGAACACAAACATCACAAAGGAGTTTCTGAGAATGCTTCTGTCTAGTGTTTCTATGAAGATAATTCTTTTTCTACCATAGGCAACAATACGCTGCAAATGAACCCTTGCAGATTT

>C19F163

CTGCAAAAGGAGTGTTTCACTCCTGCTCTGTCAAAAGACAGTTTCAACTCTGTTAGTTGAATGCACACATCTCAATGAAGTTCCTGAGAAGGCTTCTGCCTAGTTTTTTGTGAAGATATTCCCTTTTCCACCATAGGCTTCACAGCGCTCCAAATGAAAACTTGCAGGTC

>C19F164

CTACAAAAGGAGTGCTTAAATGCTGGTCTATCTAAAGACAGATTCAACTCTGTTAGTTAAATGCACACATCTCAGTGAAGTTCCTGAGAATGCTTCTTTCTAGTTCTTATGTGAATGTATCTCCTTTTCCACCATAGGCTTCACAGCGCTCCAAATGAGAACTGGCAGATC

>C19F165

AATGCACCAGTGGGCTTTTGGAGCACGTCAAGGGCTATGGTGAAAAAGGAAATATCTTCACATAAAAACTAGACAGAAGTATTCTGTAAAACTCCTTTGTGATGTTTGCATTCAACTCAGAAAGTTGAACTTCTCTTTATATAGTCCAGTTTTCAAACACTATTTTTGTAG

>C19F166

CACAGAAACTGTGTTTTAACACTGCTCTATCAAAACAAAGGTCCAAGTCTGTGAGTTGACTGCACACATCACAAAGCAGTTTCGGAGAATGATTCTGTCTAGCTTTTTATGTAACGATATTTTCTTTGCACCAGAGGCAACAAAGCACACGGAATGAAATCTTGCGGATT

>C19F16

AATCTGCAAGTGGATATTTGGAGCGCTTTGAGGGCTATGGTGGAAAAGGAAATATCTTCCCATAAAAACTAGACAGAAGCATTCTCAGAAACTTCTTTGTGATGTTTGCATTCAACTCACAGAGTTGAACATACCTTTTCATAGAGCAGTTTTGAAACACTCTTTTTGTAG

>C19F178

TACAAACTAGACAGAAGCATTCTCAGAAACTGCTTTGTGATGTGTGCATTCAACTCACAGAGTTGAACCTTCCTTTTGAGAGAGCGTTTTGAAACAGTCTTTTTGTAGTATCTGCAAGTGGATATTTGGAGCGATTTGAGGCCTAGATGGAAAAGGAAATATCTTCAC

>C19F17

AATCTGCAAGTGGATATTTGGAGCGCTTTGAGGCCTATGGTAGAAAAAGAAATATCTGCCTCTAAAAACCAGACAGAAGCATTCCGAGAAACTTCTTTGTGATGTTTGCATTCAACTAGCAGAGTTGAACCTTCCTTTTGATAGGGCAGTTTGGAAACACTCTTTTTGTAG

>C19F182

TACAAATTACACAGAAGCATTCTCAGAAACTGCTTTGTGATGTGTGCATTCAACTCACAGAGTTGAAACTTTCTTTTGAGAAAGCAGTTCTGAAACAGTCTTTTTGTAGTATCTGCAAGTGGATATTTGGAGCGATTTGAGGCCTATGATGGAAAAGGAAATATGTTCACA

>C19F190

TGTGAAGATATTTCCTTTTCCATCATAGGCCTCAAATCGCTCCAAATATCCACTTGCAGATACTACAAAAATACCGTTTCAACACTGCTCTCTCCAATGGAAGGTTCAACTCTGTGAGTTGAATGCACACATCACAAAGCAGTTTCTGAGAATGCTTCTGTCTAGTcTaGttAgaa

>C19F191

TGTGAAGATATTCCCTTTTCCATCATAGGACTCAAGTCGCTCTAAATATCCAATTTCAGATACTACAAAAAGACTGTTTCAAAACTGCTCTCTCAAAGGGAAGGTTCAAGTCCGTGAGTTGAATGCACACATCACAAAGCAGTTCCTGAGAATGCTTCTGTCTAGTTTGTa

>C19F192

TACAAAGTAGACAGAAGCATTCTCAGAAACTGCTCTGTGATGTGTGCATTCAACTCACAGAGTTGAACCTTCCTTTTGCGAGAGCTGTTTTGAAGCAGTCTTTTTGTGGTATCTGCAATTGGATATTTGGATCGATTTGAGGCCTAAGATGGAAAAGGAAATATCTTCACA

>C19F194

TACAAACTAGACAGAAGCATTCTCAGAAACTGCTTTGTGATGTGTGCATTCAACACACGGAGTTGAACCTTCCTTCTGAGAGAACAGTTTTCAAACAGTCTTTTTGTAGTATCTGCAAGTCGATATTTGGAACGATTTGAGGCCTATGAGGGAAAAGGAACTATCTTCACA

>C19F203

TACAAACTAGACAGAAGCATTCTCAGACACTGCGTTGTGATGTGTGCATTCAACTCACAGAGTTGAACCTTCCTTTTGAGAGCAGTTTTGAAACAGTCTTTTTGAAGTATCTGCAAGTGGATGTTTGGAGAGATTTGAGGCCTAAGATGGAAAAGGATATATCTTCACC

>C19F206

TACAAACTAGACAGAAGCATCTCAGAAACTGCTTTGTGATGTGTGCATTCAACTCACAGAGTTGAACCTTCCTTTTGAGAGAGAGGTTTTGAAACAGTCTTTTTGTAGTATTACAAGTGGATATTTTTAGTGATTTGAGGTCTAAGATGGAAAAGGAAATACCTTCACC

>C19F214

TCTGAAGTTATTTCCTTTTCCATCATAGGCCTCAAATCGCTCCAAATATCCACTTGCAAATACTACAAAAAGACTGTTTCAAAAGTTCTCTCTCAAAAGGAAGGTTCAACTCTGTGAGTTGAATGCACACATCACAAGGCAGTTTCTGAAAATGCTTCCGTCTAGtTTTTTA

>C19F215

TTTGAAGGTATTTCCTTTTCCTTCTTCGGCCTCAAATCACTGCAAATATCCACTTGCGGATACTACAAGAAGACTGTTTCAAAACCGCTCTCTCAAAAGGAAGGTTCGACTCCGTGAGTTGAATGCACACGTTACGAAGCAGTTTCTGAGAATGCTTCTGTCTATTTTTCA

>C19F216

TACAAACTAGACAGAAGCAATCTCATTAACTGCTTTGCGATGTGTGCATTCAGCTCACAGAGTTGAACCTTCCTTTTGAGAGAGCAGTTTTGAAACAGTTTTTTGTAGTATCCTCAAGTGGATATATGGAGCGATGTGAGGCTTAAGATGGAAACGGGAATATCTTCACA

>C19F218

TTCAAACTAGTCAGAAGCATTCTCAGAAACTGGTTTGTGATGTGTGCATTCTACTCACAGAGTTGAACCTTCCTTTTGAGAGAGCAGTTTTGAAACAATCTTTTTGTATTCTCTACAAGTGGATACTTGGAGCAATGGGAGGACTAAGATTGAAAAGGAAATATCTTCACG

>C19F220

TACAAACTAGACAGAAGCATTCTCAGAAACTGCTTTGTGGTGTGTGCATTCAACTCACAGAGTTGAACCTTCCTTCTGAGAGAGCAGTTTTTAAACAGTCTCTTTGAAATATCTGCAAGTGGATATTTGGAGCGATGGGAAGTCTAAGTTTGAAAAGGAAATATCCTCACA

>C19F227

TAAAAAGTAGACCCAAGCATTCCAGAAGTTCTTTGTGAATGTACATTGGACTCCCAGACTTGAACCTTTCTTTTGATAGAGCAGTGTTGGAACACACTTTTTGTAGAATCTTCATGTGTTCGTTTGGAGTGCTCTGTTGCCTATGGTGGAAAAAGGAATATCTTCACC

>C19F228

TAAAAAGTAGACCCAAGCATTCCAGAAGTTCTTTGTGAATGTACATTGGACTCCCAGACTTGAACTTTCTTTTGATAGAGCAGTGTTGGAACACACTTTTTGTAGAATCTTCATGTGTTCGTTTGGAGTGCTCTGTTGCCTATGGTGGAAAAAGGAATATCTTCACC

>C19F232

TACAAACTAGACAGAAGCGTTCTCAGAAACTGCTTTGTGATGTGTGCATTCACCTCACAGAGTGGAACCTTCTTTGGATAGAGCAGTTTTGAAACAGTCTTTCTCTAGTATCTGCAAGTGTTCATTTTGAGCGCTTTGAGGCCCATGATGGAAAAGGAAATATTTTCACA

>C19F238

TGTGAAGATATTCCCTTTTCCACCATAGGCTTCACAGCGCTCCAAATGAAAACTTGCAGGTCCTACAAAAAGACTGATTCAAAACTGCTCTCCCAAAAGGAAGGTTCAACTGTGTGAGTTGAATGCACACATCACAAAGCAGTTCCTGAGAATGCTTCTGTCTAGTTTGTA

>C19F239

TACAAACTAGACAGAAGCATTCTCAGAAACTGCTTTGTGATGTGTGCATTCAACTCACAGACTTGAAACTTTTTTTGATAGAGCAGTGTTGAAACACACTTTTTGTAGAATCTGCAAGTGTTCATTTGGAGGCTTTTTGCCTATGGTGGAAAAAGAAATATCTTCACA

>C19F24

AATCTGAAAGTGGATATTTGGAGCTCTTTGAGGGCTATGGCGGAAAAGAAAATATATTCACATTAAAGTAGACAGCAGCATTCTCAGAAACTTCTTTAGGATGTTTGCAGTAAACTCACAGAGTTGAACATACCTTTCCGTAGAGCAGTTTTGAAACACTCTGTTTGTGGG

>C19F257

TGCAAACTAGAAAGAAGCATTCTCAGAAACTGCTTTGTGATGGGTGCATTCAACTCAGAGACTTGAACATTTCTTTAGACGGAGCAGTGTTGAAACACACATATGCAGAATCTGCAAGAGTTCATTTGGAGCGCTTTGATGCCTATGGTGGAAAAAGAAATATCTTCACA

>C19F260

TACACACTAGTCAGAAGCATTCTCAGAAACTTCTTTGTGATGTGTGAATTGAACTCACAGAGTTGAACCTTCCTTTTGAGAGAGCCGTTTTGAAACAATCTTTTTGAAGTATCTTCAATTGGATGTTTGTAGTGATTTGAGGCCTAAGATGGAAAGGAAATATCTTCACA

>C19F264

GCCAAACTTGACAGAAGCTTTCTCAGAATCTGCTTTGTGATGTGTGCATTTACCTCACAGAGTGGAACCGTCCTTTTGATAGAGCAGTTCTGAAACAGTCTTTTTGTAGGATCTGCGAGTGTTCATTTTGGAGCGCTTTTAAGCCTTTGGCGGAAAAGGAAATATCTTCACA

>C19F267

AAAAAAGGAGACCCAAGGATTCTCAAAAAGTTCCTTGAGATGTGTGCCTTAAACTCACAGACTTCAAACTTTCTTTTGAGAGATCAGTGTTGGAACACGCTTTTTGTAGAATCTGCAAGTGTTCATTTAGTGCGCTTTGTTGCCTATGGTGGAAAAAGAAATATCTTCAAA

>C19F271

TGTGAAGTCATTTCCTTTTCCATCTAAATCGCCTCAAATCGCTAAAAGATCCACTCGCAGATACTACAAAAAGACTTTCAAAACTGCTCTCTCAAAAGGAAGGTTCAACTGTGTGAGTTGAATGCGCACATCACATAGCAGTTTTTGAGAACGCTTCTGTCTAGTTTGCA

>C19F272

TTTGAAGATATTgCTTTTTCCACCGTAGGCAACAAAGCGCTCCAAAGGAACTCTTGCGGATTCTACAAAAAGCGTGTTTCCACACTGCTCTATCAAAAGAAAGTTTCAAGTCTGTGAGTTGAATTGCACACATCCCAGTGAACTTTCGGAAAATGCATGGGTCTACTTTTCA

>C19F273

TATGAAGATAATTCTTTTTCTACCATAGGCAACAATACGCTGCAAATGAACACTTGCAGATTTCACAGAAACTGTGTTTTAACACTGCTCTATCAAAACAAAGGTCCAAGTCTGTGAGTTGACTGCACACATCACAAAGCAGTTTCGGAGAATGATTCTGTCTAGCTTTTTA

>C19F277

TAAAAACTAGACAGAAGCATTCTCAGAAACTCCTTTGTGATGTGTGTGTTCAATTCACAGAGTTGAACCTTTCTTTTGATAGAGCAGTTTTGAAACACTGCTTTTGTAGAATCTGCTTGTGGATATTTGGAGCTCTTTGAGGAATTCGTTGTAAACGGGATATCTTCACA

>C19F281

TAAATACTAGACAGAAGCATTCTCAGAAACGCCTTAGTGATGTGTTTGTTCTATTCAGAGAGTTGAACCTTTCTTTTGATAGAGCAGTTTTGATACACTGCTTCTGTAGAATCTGCTTGTGGATATTTGGAGCTCTTTGAGGAATTCGTTGTAAACGGGATATCTTCACA

>C19F286

TAAAAACCAGACAGAAGCATTCTCAGAGACTGCTTTGTGATGTGTGTGTTCAATTCGCAGAGTTGAAAGTTGCTTTTGATAGAGCAGTTTTGAAACACTGCTTTTGTAGAATCTGCTTGTTGCTATTGGGGGCTCTTTGAGGAATTTGTTGTAAACGGGATATCTTCACA

>C19F288

TAAAAACTAGACAGAAACATTCTCAGAAAATACTTTGTGATGTGGTTGTTCAATTCACAGGGTTGAACCTTTCTTTAGATAAAGCAGTTTTGAAACACTGCTTTTGTAGAATCTTCTTGTGGATATTTGGAGCTGTTTGAGGAATTCGTTTTAAACGGGATATCTTCACA

>C19F292

TAAAGACTAGAAAGAAGCGTTCTCCGAAACTCCTTTGTGATATATGTGTTCAGTTCACAGAGTTGAACCTTTCTTTTGATTGAGCAGTTTTGAAACACTGCTTTTCTAGAATCTGCTTTTGGATATTTGAAGCTCTTTGACGAATTCGCTGTCAATGTTATATCTTCACA

>C19F295

TACAAACTAGACAGAAGCATTCTCAGAAACTGCTTTGTGATGTGTGTGTTCAATTCACAGGGTTGACTCTTTCTTTTGATTGAGCAGTTTTGAACCACCTGTTTTGTAGAATCTGCTTGTGGATATTTGTAGCTCTTGGAGGAATTCTTTGTAAAAGGGATATCTTCACA

>C19F297

TGTGAAGAGATTCCATTTACAACGAATTCCTCAAAGAGCTCCAATTATCCACAAGCAGAGTCTACAAAAGCAGTGTTTTGAAACTGCTCTGTCAATAGAAAGGTTCAACTCTGTTAATTGAGCACACACATCCAAAGGAGTTTCTGAGAATCCTTCTGTCTAGTTTTTA

>C19F298

TGAAAACTAGACAGAAACATTCTCAGAAACTCCTTTGTGAAGTGTGTGTCAAATTCACAGAATTGAAATATTCCTTTGATAGCGCAGCTTTGAAACACCGCTTTTATAGGATCTGCTTGTGGATATCTGGAGCTCTTTGAGGAATTTGTTGTAAACGGGATATCTTCACA

>C19F302

TACAAACTAGAAAGAAGCCTTCTCAGAAACTCCTTTGAGATGTTTGTGTCCAATTCACAAAGTTGAACCTTTCTTTTGATAGAGCAGATTTGAAACACTGCTTTTGTAGAATCTGCTTGCGTTATTTGGAGGTCTTTGAGGAATTGGGCGTATACGGGATATCTTCACA

>C19F309

TACAAACTAGAAAGAAGCCTTCTCAGAAACTCCTTTGAGATGTTTGTGTCCAATTCACAAAGTTGAACCTTTCTATTGATACAGCAGATTTGAAACTCTGCTTTTGTAGAATCTGCTTGTGAATATTTGGAGGTATTTGAGGAATTGGACGTATACGGGATATCTTCACA

>C19F310

AAAAACTAGACAGAAGCTTCTCAGGAACTTCATTGAGATGTGTGCATTCAAGTAACTGAGTTGAATATGTCTTTTGATAGAGCAGTATTGAAACACTCTTTTGTAGAATCTGCTGTGGATATTGGAACTCTTTAGAATTCTTGGAACGGTATCTTCACA

>C19F314

AAAAACTAGACAGAGCTCTCAGGAACTTCATTGAGATGTGTGCATTCAAGTAACTGAGTTGAATATGTCTTTTGATAGAGCAGTATTGAAACACTCTTTTGTAGAATCTGCTGTGGATATTGGAACTCTTTAGGAATTCTTGGAAACGGTATCTTCACA

>C19F315

AAAAACTAGACAGAAGCTTCTCAGGAACTTCATTGAGATGTGTGCATTCAAGTAACTGAGTTGAATATGTCTTTTGATAGAGCAGTATTGAAACACTCTTTTGTAGAATCTGCTTGTGGATATTGGAACTCTTTGAGGAATTCTTGGAAACgGTATCTTCACA

>C19F31

AATCTGCCAGTGGATATTTGGAGCGCTTGGAGGGCTATTGTGCCAATGGAAATATCTGCCCCTGAAAACTAGACAGAAGCATTCTCAGAAACTACTTCGTGATGTTTGCATTCAACTCACAGAGTTGAACATACCTCTTCACAGAGCAGTTTTGAAAACCTCTTTCTGTAG

>C19F322

TGTGAAGATACCCGTTTCCAACGAATTCTTCAAAGAGTTCCAAGTATCCACAAGCAGATTCTACAAAAGGAGTGCTTAAATGCTGGTCTATCTAAAGACAGATTCAACTCTGTTAGTTAAATGCACACATCTCAGTGAAGTTCCTGAGAATGCTTCTTTCTAGTTCTTA

>C19F323

TGTGAAGATGTCCCGTTTACAACGAATTCCTCCGAGAGCTCTAAATATCTGGAAGCAGATTCCACAAAAGCAGTGCTTCAAACCTGCTCTATCAAAGGAAAGGTTCAGCTCGGGGAATTGAACACAAACATCACAAAGGAGTTTCTGAGAATGCTTCTGTCTAGTGTTTC

>C19F324

TGTAACGATATTTTCTTTGCACCAGAGGCAACAAAGCACACGGAATGAAATCTTGCGGATTCTACAAAAAGTGTGTTTCAACACtGCTCTATCCAAAGAAAGGTTCGAGTCCGTGAGTTGAACGCACACAACAGAAAGCAGTTTCTGAGGATGCTTCTGTCTGGTTTGTA

>C19F325

TGTGAATGTATCTCCTTTTCCACCATAGGCTTCAAAGCGCTCCAAATGAGAACTGGCAGATCCTCATAAAAGACGGCCTCAAAACCGCTCTATCAAAAGAAGGGTTCCACTCCAAGAGGTGAATTCACATATCACGAAGAAGTTTCTGAGAATGCTTCTGTCCAATTTgTA

>C19F326

GGTGAAGATATCACTTTTTCCAACATACGTAAAAAAGTACTCGAAATGAACACTTGCAGATTCTACAAAAAGGATGTTTCAACACTGCTCTATCAAAAGAAAGGTTCAACGATGTGAATTGAACACACACATCACAAAGGAGTTTCAGAGAATGCTTCTGTCTAGTGTTTA

>C19F38

AATCTGCAGGTAGACATTTGGGGTGCTTTGAGGGCTGTGGTGCAAAAGGAAATGTCTTCCCATAGAAACTAGACTGAAGCATTCTCAGCAACTTCTTGGTGACGTTTGCATTCATCTCACAGTGTTGAACATACCTTTCCATAGAGTAGTTTTGAAACACTGTTTTTGTAG

>C19F44

AATCTGCAAGTGGATATTTGGACTGCTTTGAGGCCTTCATGGAAACGGGAATATCTTCACATAAAAACTAGAAGAAGCATTCTCAGAAACTTCTTTGTGATGTGTGCATTCAACTCACAGAGTTGAACCTTCCTTTTATAGAGCAGTTTTGAAACACTCTTTTTGTAG

>C19F46

AATCGGCAAGTGGATATTTGGACTGCTTTGAGGCCTTCATCGGAAACGGGAATATCTTCACATAAACACTAGAGAGAAGCATTCTCAGAAACTTCTTTGTGATCTGTCCATTCAACTCACAGAGTTGAACCTTCCTTTTTATGGAGCAGTTTTGAAACACTCTTTTTGGAG

>C19F54

AATCTGCAAGTGGATATTCGGACCACTTTGAGGCCTTCATAGGAAACAGTAATATCTTCACATAAAAACTAGATAGAAGCATTGTCAGAAAGTTCTTTGTGATGTGTGAATTCAACTCACAGAGTTGAACCTTCCTTTAATAGAGCAGTTTTGAAACACTCTTTTTCTAG

>C19F60

CTACAAAAAGACTGATTCAAAACTGCTCTCCCAAAAGGAAGGTTCAACTGTGTGAGTTGAATGCACACATCACAAAGCAGTTCCTGAGAATGCTTCTGTCTAGTTTGTATGTGAAGAcATTTCCTTTTCCATCATAGGCCTCAAATCGCTCCAAATATCCACTTGCAGATA

>C19F61

CTCCAAAAAGACTGTTTCAAAACTGCTCTCTCAAAGGGAAGGTTCAAGTCCGTGAGTTGAATGCACACATCACAAAGCAGTTCCTGAGAATGCTTCTGTCTAGTTTGTTTCTGAAGTTATTTCCTTTTCCATCATAGGCCTCAAATCGCTCCAAATATCCACTTGCAAATA

>C19F62

GAATCTGCCAGCGGACACTTGGAGCGCTTTGAGGGCTATGGTGGAGAAGGAAATATCTTCCCATAAAAACTAGAAAGAAGCATTCTCAGAAACATTTATGTGAAGCGTGCATTCAACTCACAGAGTTGAACCTTCCTTTTGATACAACAGTTTTGAAACACTCTTTTGAAC

>C19F68

TACAAAAAGACTTTCAAAACTGCTCTCTCAAAAGGAAGGTTCAACTGTGTGAGTTGAATGCGCACATCACATAGCAGTTTTTGAGAACGCTTCTGTCTAGTTTGCATGTGAAGAGATTCCATTTACAACGAATTCCTCAAAGAGCTCCAATTATCCACAAGCAGAGTC

>C19F69

AATCAGCAAGTGGATATTGGGAGCGCTTTGAGGCCTCTGGTGGAAAGGGAATGTCTTCACATAAAAACTGGACAGAAGCATTCTCAGAAACATCTTTGTGATGTTTGCATTCAACTCACAGAGTTGATCCTTCCTTTTAATAGGGCAGTTTTGCAACACTCTTTTTGTAG

>C19F70

CTACAAAAAGACTGTTTCAAAAGTTCTCTCTCAAAAGGAAGGTTCAACTCTGTGAGTTGAATGCACACATCACAAGGCAGTTTCTGAAAATGCTTCCGTCTAGCTTTTTATTTGAAGGTATTTCCTTTTCCTTCTTCGGCCTCAAATCACTGCAAATATCCACTTGCGGATAC

>C19F71

AATCTGCAAGTGGATACTGGGACTGCTTTGAGGCCTTCGTTGGAAACGGGATTATCTTCACATAAAAACTAGACTGAAGGATTCTTAGAAACTTCTTTGTGATGTGTGCATTCAACTCACCGAGTGGAACCTCACTTTTGATAGAGCAGTGTTGAAAGACACTTGTTGTAG

>C19F72

AATCTGCAAGTAGATATTTGGAGCGCTTTGAGGCCTTCGTTGGAAACCGGAATATCTTCACAGGAAAAGTAGATAGAGGCATTCTCAGAAACTTTTTTGTGATATGTAGATTCAACTCACAGCGTTGAACCTTTCTTTGGATGGAGCAGTTTTGAAAAACTCTTTTATCG

>C19F78

GAATTTGCAAGTGGAGATTTCAGCCGCTTTGAGGTCAATGGTAGAATAGGAAATATCTTCCTATAGAAACTAGACAGAATGATTCTCAGAAACTCCTTTGTGATGTGTGCGTTCAACTCACAGAGTTTAACCTTTCTTTTCATAGAGCAGTTAGGAAACACTCTGTTTGTA

>C19F90

GAATTTGCAAGTGGAGATTTCAAGCGCTTTGGGGCCAAAGGCAGAAAAGGAAATATCTTCGTATAAAAACTAGACAGAATCATTCTCAGAAACTGCTCTGTGATGTGTGCGTTCAACTCTCAGAGTTTAACTTTTCTTTTCATTCAGCAGTTTGGAAACACTCTGTTTGTA

>C19F94

ATCTGCAAGCGGATAATTGGCTTCGCTTTGTGTCCTTTGGTGGAAACGGGAATATCTTCTAATAAAAACTAGACAGAAATATTCTCAGAATCTCCTTTGTGATGTGGGCATTCAACTAACACAGTTGAACATTTCTTTTCACAGAGCAGTTTTGAAACACTCTTTTGGTAG

>C19F99

ATCCGCAAGTGGATATTTGGACCGCTTTGAGACCTTTGCTGGAAATGGGAATATCTTCACATATAAACTAGACAGAAGCATTCTCAGAAACTTCTTCGTGATGTGTGCATTCTACTCCCGAATTTGAATCTTCCTTTTCATGAAGCAGTTTTGAAACACTCTGTTTGTGC

>C1F111

AAGTCTGCACGTGGAATTTTGACCACTTAGAGGCCTTCGTTGGAAACGGGTTTTTTTCATGTAAGGCTAGACAGAAGAATTCCCAGTAACTTCCTTGTGTTGTGTGCATTCAACTCACAGAGTTGAACGTTCCCTTAGACAGAGCAGATTTGAAACACTCTATTTGTG

>C1F56

GAATTTGCAAGTGGAGATTTCAAGCGCTTTGGGGCCAAAGGCAGAAAAGGAAATATCTTCGTATAAAAACTAGACAGAATCATTCTCAGAAACTGCTGCGTGATGTGTGCGTTCAACTCTCAGAGTTTAACTTTTCTTTTCATTCAGCGGTTTGGAAACACTCTGTTTGTA

>C1F5

AATTTGCAAGTGGAGATTTCAACCGCTTTGGCAAGTAGAAAAGGAAATATCTTCGTAAACTAGACAGAATCATTCCAAAAATCCTTTGTGATGTGTTCGTTCAACTCACAGAGTTTAACTTTCTTTTCATACAGCGGTTGAAACACTCTGTTTGTA

>C1F63

GAATTTGCAAGTGGAGATTTCAGCCGCTTTGAGGTCAATGTAGAAAAGGAAATATCTTCGTATAGAAACTAGACAGAATGATTCTCAGAAACTCCTTTGTGATGTGTGGTTCAACTCACAGAGTTTAACCTTTCTTTTCATAGAGCAGTTAGTAAACACTCTGTTTGTA

>C1F83

CAATTTGCAAGTGTAGATTTCAAGCGCTTTAAGGTCAAGGCAGAAAAGGAAATATCTTCGTTTCAAAACTAGACAGAATCATTCCCACAAACTGCGTTGTGATGTGTTCGTTCAACTCACAGAGTTTAACCTTTCTGTTCATAGAGCAGTTAGGAAACACTCTGTTTGTA

>C1F88

AAGTCTGCAAGTGGATATTCAGACCTCTTGAGGCCTTCGTTGGAAACGGGATTTCTTCATATTATGCTAGACAGAAGAATTCTCAGTAACTTCCTTGTGTTGTGTGTATTCAACTCACAGAGTTGAACGATCCTTTAACAGAGCAGACTTGAAACACTCTTTTTGTG

>C1F90

AAGTCTGCAAGTGGATATTCAGACCTCTTGAGGCCTTCGTTGGAAACGGGATTTCTTCATATTTGCTAGACAGAAGAATTCTCAGTAACTTCCTTGTGTTGTGTGTATTCAACTCACAGAGTTGAAATCTTTAACAGAGCAGACTTGAAACACTCTTTTTGTG

>C1F119

CTAGAAAGAGAGGGTTTCAAAGCTGCTCTATCAAAAGGAAAGTACAACTCTGTGAGTTGAATGCAAACATCACAAAGAAGTTCCTGAGCATGCTTCCGTTTAGCTTTTATGGGAAGATTATCCCTTTTCCATCGAAATGTTCAAAGAGGTCCACATATCCGCTTGCAGATT

>C1F126

CTACGAAAAGAATGTTTCAAAACTGCTCTATGAAAAGCAATGTTATACTCTGGGAGTTGAACACAAGCCTCACAAAGGAGTTTCTGAGAATGCTTCTGTTTACTTTTTACGTGAAGATATTCCCGTTTCCAAAGAAATCTTCACAGAGTTCCACCTATCCATTTGCAGATG

>C1F132

CTCAAAAAGAGTGTTTCAAACCTGAACTATCAAAGGAAGGTTCAACTCTGTAGTTGAATGCAAACATCACAAAGAAGTTTCTGGAATGCTTCTGTTTAGTTAGGTGAGGTTTATCCCGTTTCCAACGAAATCCTCAGAGAGGTCCAAATATCCACTTGCAGATTC

>C1F133

CTACGAAAAGAGCGTTTCAACCTGAACTCACAAGGGAAGGTTCAACTCTGTCAGTTGAATGCCAACATCACAAAGAAGTTCTGGGAATGTTTCTCTTCAGTTATGTGAGTTTTATCCCGTTTCCAACGAAATTCTCAGAGAAGTACAAATATCCACTTGCATATTC

>C1F14

TACAAAAAGTGTGTTTTGAAAGTGCTCCATCAAAAGATATGCTCAGCTCTGTGAGTTAAACTCAATCATCACAAAGAATTTTCTGAGAATGCTTCTGTCTTGTTTTAGGATGAAGTTATTTCCTTTACGACGATAGGCCTCAAAGAGGTCCAAATCTCCACTTGCAGATT

>C1F150

CTCCAACAAGAGTGTTTCAAACGTGAACTATCAAAGGAAGGTTCAACTCTGGACTTTGAATGCAAACGTCAGAAAGATGTTTCTGCGAAAGCTTCTGTTTAGTTAGGTGACGTTATCCCGTTTCCAACGAAATCCTCAGAGAGGTCCAAATATCCACCTGCAGATTC

>C1F21

TACAGAAAGTGTGTTTTGAAACTGCTCCATCCAAAGGAATGTTCAGCTCTGTGAGTTGAACTCAATCGTCACAAAGTGTTTCCTGGGAATGCTACTGTCTAGTTTTTATGGGCAGTTATATCCTCTGCTGCCATAGGCCTCAAAGCGGTCCAAATCTCCCCTTTCAGATT

>C1F22

TACAGAAAGTGTGTTTTGAAACTGCTCCATCCAAAGGAATGTTCAGCTCTGTGAGTTGAACTCAATCGTCACAAAGTGTTTCCTGGGAATGCTACTGTCTAGTTTTTATGGGCAGTTATATCCTCTGCTGCCATAGGCCTCAAAGCGGTCCAAATCTCCCCTTTCAGATT

>C1F30

CTACCAAAAGTGTGTTTCCAAACGGCTCTATCAAAGGGAATGTTCAACTCTGTGACTTGAATGCAATCATCACAAAGCAGTTTCTGAGAATGCTTCCATGTAGCTTTTATGAGAGATATTTCCTTTTCCACCCCAGGCCTCGAAGCCCTCCAAATGTCCCCTTGCAGATG

>C1F36

CCACCGAAAGAGTGTTTCCAAACTGCTGTATCAAAAGGAATCTTCAACTCCGTGAGTTGAATGCAATCATCACAAAGAAGTTTCTGACAATGCTTCTCTCTAGTTTTTATGTGAAGATATTTCCTTTTCCACCACAGGCCTGAAAGCGCTCCAAATGTCCACTTGGAGACT

>C1F37

CCACCGAAAGAGTGTTTCCAAACTGCTGTATCAAAAGGAATCTTCAACTCCGTGAGTTGAATGCAATCATCACAAAGAAGTTTCTGACAATGCTTCTCTCTAGTTTTTATGTGAAGATATTTCCTTTTCCACCACAGGCCTGAAAGCGCTCCAAATGTCCACTTGGAGACT

>C1F46

CTAAAAAAGAGAGTTTCAAAACTGCTCTATCAAAAGGAATGTTCAACTCTGTGAGTTGAATGCAATCATCACAGAGAAGTTTCTGAGAAGGCTTCTGTCTAGATTTTATGTGAAGATATACCGTTTCGAACGAAGGCCACAAAGTGCTCCAAATATCCACTTGCAGTC

>C1F47

CTAGAAAAAGAGAGTTTCAAAACTGCTCTATCAAAAGGAATGTTCAACTCTGTGAGTTGAATGCAATCATCACAGAGAAGTTTCTGAGAAGGCTTCTGTCTAGATTTTATGTGAAGATATACCCGTTTCGAACGAAGGCCACAAAGTGCTCCAAATATCCACTTGCAGTC

>C1F8

TGCAAAAAGTGTGTTTCCAAACTGCTCCACCCAAAGGAATGTTCAGCTCTGTGAGTTAAACTCAATCATCACAAAGTATTTTCTGAGAATGCTTCTGTCCAGTTTTTACATGAAGCTGTTTCCTTTACTACCGTAGGCCTCAAAGCGTTCCAAATCTCCACTTGCAGATA

>C1F117

AAGTCTGTAAGTGGATATTCTGACATCTTGTGGCCTTCGTTGGAAATGGGATTTCTTCATATTCTGCTAGACAGAAGAATTCTCAGTAACTTCCTTGTGTTGTGTGTATTCAACTGACAGAGTTGAACTTTCATTTAGAGAGAGCAGATTTGAAACACTGTTTTTGTG

>C1F81

GAATTTGCAAGTGGAGATTTCAGCCGCTTTGAGGTCAATGGTAGAATAGGAAATATCTTCCTATAGAAAATAGACAGAATGATTTTCAGAAACTCCTTTGTGATGTGTGCGTTCAATTCACAGAGTTTAACTTTTCATAGAGCAGTTAGGAAACACTCTGTTTGTA

>C20F101

CTACAAAAAGAGTGTGTCAAAACTGCTCAATCAAAAGAAAGGTTCTACTCTGTGAGATGGATGCAAACATCAGAAAGTAGTTTCGCAGAATTCTTCTGCATAGTTTTTATGTGAAGATATTTCCTTCTCCACTATAGGTCTCAAAAGGCTCCAAATATCCACTTGCAGATT

>C20F103

CTACAAAAAGAGTGTTTCAAAACTGCTCAATCCAAAGAAAGGTTGTACCCTGTGAGATGAATGCACGCATCACAAAGTAGTTTCTCAGAATGCTTCTGCATAGTTTTTATGTGAAGATATTTCCTTCTACACTGTAGGCCTGAAAAGGCTCCAAATATCCATTTACAGATT

>C20F105

CTACAGAGAGAATGTTTCAAAACTGCTCAGTCAAAAGAAAGGTTGTACTCTGTGAGGTGAATGCACACATCACAAAGTAGTTTCTCAGAATGCTTCTGCATAGTTTTTATGTGAAGATATTTCCTTCCCCACTATAGGCCTCAAAGGCTCCAAATATCCACTTGTAGATT

>C20F106

CTGCAAAAAGACTTTTTCAAAACTGCTCAATCAAAGGAAAGGTCAACTCTGTGAGTTGAATGCACACAACACAAACAAGTTTCTGAGAATGCTGCTGTCTAGATTTTATGTGCGGATATTTCTTTTCCACCATAGGCATCAAAGCGCTCCAAATATCCAACtGCAGATT

>C20F108

CACAAAAAGAGTGTTTCAAAACTGTACTGTCAAAAGAAATGTTCAACTCTGTTAGTTGAGGACACACATCAGAGACTAGCTTCTGAGAATGCTTCTGTCCAGTTGTTACGGGAAGATATTTCCTTTTTCAACATAGGCCTGAAACCGCTTCAAATGTCCACTTCCAGATAC

>C20F112

ATATCTGGAAGTGTCCATTCGGAGCGCATTCAGGCTTGTGTTGAAAAAGGAAATATCCTCCCATAAAAACTAGACAGAAGCATTCTCAGAAACTTATCTGTGATGTATGTACTCAACTAACAGAACTAAACCATCGTTTTGAAGGAGCAGTTTTGAAACACTCTTTTTGCG

>C20F119

AATCTGCAAGTGGACATTTGGAGTGATTTGAGACATAGGGTGAAAAAGGAATATCTTCCCATAAAAAGTAGACAGAAGCATTCTCAGAAACGACTTTCTGATTTGTGTACTCAACTCACAGAGTTAAACCTTTCCTTTGATACAGCAGTTTGGAAACACTCTTCTTGTAG

>C20F122

TACAAAATGAGTGTTTCAAAACTGCTCAATCATTAGATAGGTTCAACCCTGTGAGATGAATGCACACATAACAAAGAAGTTTTTCAGAATGCTTCTATGTAGTTTTTATTTGAAGGTATTTCCTTTACCACCATAGGTTGCAAGGGGCTCCAAATATCCACTTGCAGATT

>C20F125

GGAAAAAGAGTGTTTCCAACCTGCTCTATGAAAGCGAATGTTCAACTCCGTGACTTGAATGCAACCATCACAAGGAAGTTTCTGAGAATGCTTCTGTCTAGATTTTATATGAAGATATTCCCGTTTCCAACGAAATCCTCAAAGCTATCCAAATATCCACTTGGAGATTC

>C20F130

AATCTGCAAGAGGATATTTGGATAGCTTTGAGGATTTCGTTGGAAACGGGAATGTCTTCATATAAACTCTAGACAGAAGCATTCTCAGAAACTTCTTTGGGATGTTTCATTGAAGTCCAGTGTTGAACATTCCCTTTCATAGAGCAGGTTTGAAACACTCTTTTTGTA

>C20F136

GAATCTGCAAGAGGATATTTGGATAGCTTTGAGGATTTCGTTGGAAACGGGAATGTCTTCAGATAAACTCTAGACAGAAGCATTCTCAGAAACTTCTTTGGGATGTTTCAATTGAAGTCACAGTGTTGAACATTCCCTTTCATAGAGCAGGTTTGAAACACTCTTTTTGTA

>C20F150

AAATCTGCAAGAGGATATTTGGATAGCTTTGAGGATTTCGTTGCAAACGGGAATGGCTTCATATAAACTCTAGACAGAAGCATTCTCAGAAACTTCGTTGGGATGTTTCGATTGAAGTCCCAGTGTTGAACATTCCCTTTTATAGAGCAGGTTGGAAACACTCTTTCTGCA

>C20F155

TACGACAAAAGGAGTGTTTCAAACCTGTTCTGTGAAAGGGAACGTTCCATTCTGTGACTTGAATGCAAACATCACCAAGTAGTTTCTCAGAACGCTGCTGTCTGCTTTTTATATGTATTCCCGTTTCCAACGAAATCGTCAAAGCCAGCCAAATATCCACTTGCAGGTTC

>C20F159

GAATTGCAAGTGGATATTAGGCAGCTTTGAGGATTTCGTTGGAAACGGGAATACATATAAAAAGCAGACAGCAGCATTCTCAGAAACTTCTTTGTGATGTTTGCATTCAAGTCACAGAGTTGAACATTCCCTTTCAAGAGCAGGTTTGAAACACTCTTTTTGTA

>C20F15

TATCTGAAGTGACATTTGGATGCTTTAGGCCTAGTGAAAAAGGAAATATCTTCCCATAAAAACTAGACAGAAGCATTCTCAGAAACTTTTTGTGATGTGTGCCTCAACTACAGAGTTGAACCTTTCTTTTAAGAGCAGTTTTGAAACACTCTTTTTGTAG

>C20F166

GAATCTGCAAGTGGATATTTGGCTAGCTTGGAGGATTTCGTTGGAAACGGGATTACATATAAAAAGGAGACAGCAGCATTCTCAGAAACTTCTTTGTGATGTTTGCATTCAAGTCACAGAGTTGAACATTCCCTTTCATAGAGCAGGTTTGAAACACTCTTTTTGTA

>C20F16

TATCTGAAGTGGATTTGGAGCGCTTTGAGGCCTAGGTGAAAAAGGAAATATCTTCACCTAAAAACTAGACAGAAGCATTCTCAGAAACTTCTTTGTGATGTGTGTATCAACTCACAGAGTTGAACTTTCTTTTGATAAGCAGTTTTGAAACACTCTTTTTGTAG

>C20F175

GAATATGCAAGTGGATATTAGGGCAGCTTTGAGGATTTCGTTGGAAACGGGAATACATGTAAAAAGCAGACAGCAGCATTCTCAGAAACTTCTTTGTGATGTTTGCATTGAAGTCACAGAGTTGAACATTCCCTTTGAGAGAGCAGGTTTGAAACACGCCTTTTGTC

>C20F180

GAATATGCAAGTGGGTATTAGGCCAGCTTGGAGGATTTCGTTGGAAACGGGAATACGTATAAAAAGCAGACAGCAGCATTGTCAGAAACTACTTTGTGATGTTTGCATTCAAGTCACAGAATTGAACACTCCCTTTCACAGAGCAGGTTTGAAACACTCTTTTTGTA

>C20F187

GTATCTGGAAGTGGACATTTGGAGCGCTTTCAGGCCTATGTTGAAAAAGGAAATATCTTCCCATAAAAACTAGACGGAAGCATTCTCAGAAACTTATTTGTGATGTGTTTGCTCAACTAACAGGATTGAACCATCGTTTTGAAGGAGCAGTTTTGAAACACTGTTTTCGTG

>C20F18

GTACAAAAAGAGTGTTTCAAAACTGCTCTATCAAAAAgAAAGGTTCAACTCTGTGAGTTGAATGCACACATCACAAAGAAGTTTCTGAGAATGCTTCTGCTCTAGTTTTTTTTGTGAAGGTGTTTCCTTTTCCACCATAGGCCTCAAAGCGCTCCAAATATCCACTTGCAGATT

>C20F194

AATTTGCAAGTGGATATCTGGACAGCTTTGAGGCCTTCCCTGGAAACGAGTGTATCTTCACATAAACACTAGACAGAAGCATCCTCAGAAACTTGTTGGTGATGCTTGCATTCAACTCACAGAATTGAACATTTCTTTTCCTAGAGCAGTTTTGAAACACTCTTTTTGTAG

>C20F196

TACAAAAAGAGAGATTCAAAACTGCTCAATGAAAAGATAGGTTCAACTCTGTGAGATGAGTGCACATCTCACAAAGAAGTTCCTCAGAATGCTTCTGTGTAGTTTTTATGTGAAGATATTTCCTTTTCCAAAATAGACCTCAAAGTCCTCCAAATATCCACTTCCAGACT

>C20F197

CTACAAAAAGAGAGATTCTAAACTGCTCAATCAACAGATACGTTCAACAATGTGAGTTGAATGCACACATCACAAATAAGTTTCACAGAATGCTTCTGGGTAGTTTTTATTTGAAGAAATTTCCCTTTCCACAATAGGCCTCAAATCGCTCTAAATATCCACTTGCAGATT

>C20F202

CTGCAAAAAGGGAGATTCTAAACTGCTCAATCAACAGATACCTTCAACAATGTGATTTGAATGCACACATCGCAAATAAGTTTCACAGAATGCTTCTGGGTAGTTTTTAGTTGAAGAAATTTCCCTTTCCACAATAGGCCTCAAATCACTCTAAATATCCACTTGCAGATT

>C20F205

CTACAAAAAGAGAGATTCTAAACTGCTCAATCAACAGATACGTTCAACAATGTGAGTTGAATGCACACATCACAAATAAGTTTCACAGAATGCTTCTGGGTAGTTTTTATTTGAAGAAATTTCCCTACCCAGCTCAGGCCTCAAGTCGCTCTAAATATCCACTTGCAGATC

>C20F207

CTGTAAAAAGAAAGATTCAAAACTGCTGAATCAAAAGATACGTTCAACAATGTGAGCTGAATGCACACATCACAAATAAGGTTGACAGAATGCTTCTGGGTAGTTTTTATTTGAAGAAATTTCCCTTTCCACAATAGGCCTCAAATTGCTCTAAATATCCACTTGCAGATT

>C20F208

CTACAAAAAGAGTGTTTTGAAACTGCTCAATCATATGATAGGTTCAACCTTGTGAGATGAATGCCCACATCACAAAGTAGCTTCTCAGAATGTTTTTGTGTAGTTTTTATTTGAAGATATTTCCTTTTCCACCATAGGCCACAAAGGGCTCCAAACTTCCACTTGCAGATTC

>C20F209

CCATGACAATAGTGCTTCAAAACTGCTCAATCCAAAGAAATGTTCAACACTGTGTGATGAATGCACTCATCACAAAGAAGTTTCTCTGAATGCTTCTGTGTATTTTGTATTTGAAGATATTTCCTCTTCCACCATAGGGCTCATAGGGCTCCAAATATCCACTTGCAGATT

>C20F213

CTATCTGGAAGTGGACATTTGGAGCGCTTTCAGGTCTACGGTGAAAAAGGAGATATCTTCCAATAAAAACTAGATAGAAGCAATGTCAGAACTTTTTTCATGATGTATCTACTCAGCAAACAGAGTTGAACCTTTCTTTTGAGAGAGCAGTTTTGAAACACTCTTTTTGTG

>C20F220

TTCCCTGGAAGTGGACATTTGGAGCGCTTTCAGGACGACGGTGAAAATGGAAATATCTTCCAAGAAAATCTAGATAGAAGCAATGTCAGAAACTTTTATGTGATGGATCTACTCAGCTAACAGAGTTGAACCTTTCTTTTGAGAGAGCAGTTTTGCAACACTCTTTTTGTG

>C20F227

AATTTACAAGCGGATATAAGGACAGCAGTGAGGATTTCATCGGAAACGGAATATCTTCACATAAAAGTAGACAGAAGCATTCTCAGAAAGTTCTTTGTAATGTGTGCATTCAACTCACAGAGTTGAAACTTTCTTTTGATAGAGCAGATTGGAAGCCCTCTTTTTGTAG

>C20F230

CTGAAATAAGAGAGATTCAAAACTGCTCAATCAAACGATAGGTTCAAATCTGTGAGTTGAATGCACACCTCACAAAGAAGTTTCTCATAAAGTTTCTGTGTAGTTTTTATGTGAAGATATCTCCTTCTCCAAAATAGGCCtCAATGACCTCCAAATATGCACTTCCAGATT

>C20F232

TACAAAAAGAGAGATTAAAAAATGTTCTATCAAAATATAGGTTCAACTCTGTGAGTTGAATGCGCACATCACAAAAAGTTTCTCAGAATGCTTCTGTGTAGTTTTTACGTGAAGATATCTCCTTTTCCACAATAGGCCTCAAAGCTCTCCAAATATCCGCAAGCAGAGTC

>C20F233

CTGTAAAAAGAGAGATTCAAAACTGCTGAATCAAAAGATAGGTTCAACACTGTGACTTCAGTGCACAACTCACAAAGATGTTTCTCAGAAATCTTCTGTGTAGTTTTTATATGAAGATATCTCCTTCTCCAAAACAGAACTCAAAGCCCTCCAAATATTCACTTCCAGATT

>C20F238

CTGTAAAAAGAGAGATTCAAAACTGGTGAATCAAAAGATAGGTTCAACACTGTGACTTCAGTGCACAACTCACAAAGGTGTTTATCAGAAATCTTCTGTACAGTTTTTATATGAAGATATCTCCTTCTCCAAAACACAACTCAAATCCCTACAAATATTCACTTCCAGATT

>C20F239

CTGTAAAAAGAGAGATTCCAAACTGCTGAATCAAAAGATAGGTTCAACACTGTGACTTAGGTGCACAATTCACAAAGATGTTCCTCAGAAATCTTCTGTGTAGTTTTTATATGAAGATATCTCCTCCTCCAAAACAGATCTCAAAGCCCTCCAAATATTCACTTCCAGATT

>C20F242

TACGGAAAGATTGTGTCAAAACTGCTAAATCAAAACAAAGGTTCAACTCTGTGATGAATGCACTCATCAGAAAGAAGGTTCTCTGAATGCTTCTGTGTAGTTTCTATTTGAAGATATTTCCTTTTCCACTCTAGGGCGAAATAGGGCTCCAAATATTCACTTGCAGATT

>C20F246

GTACGGAAAGATTGTGTCAAAACTGCTAAATCAAAACAAAGGTTCAACTCTGTGATGAATGCACTCATCAGAAAGAAGGTTCTCTGAATGCTTCTGTGTAGTTTCTATTTGAAGATATTTCCTTTTCCACTCTAGGGCGAAATAGGGCTCCAAATATTCACTTGCAGATT

>C20F249

CTGTAAAAAGAGAGATTCAAAACTGCTGAATCAAAAGATAGGTTCAACACTGTCACTTCAGTGCACAACTCACAAAGATGTTTCTCAGAATGCTTCTGTGTAGTTTTTGTGTGAAGATAATTCATTTTCCACAGTACGCCTCAAAGCGCTCCAAATATCCACTCACAGATT

>C20F250

CTATAAAAAGGTGTTTCAAAACTACTCAATCAAAAGAAAGGTTCAACTATATGAGATAAATGCACACATCACAAAATGGTTTCTCAGAATGCTTCTGTGTAGTTTTTATTTGAAGATATTTTCTTTTCCACCATGGGCCACAAAGGGCTACTAAAACCCACTTGCAGATTC

>C20F251

TACAAAAAGAGAGATTCAAAACGGCTCAATCAAAAGATAGGCTCTCCCCTGTGAATTGAATGCACACATCACAAAGAAGTTTCTCAGAATGTTTCTGTGTAGTTTTTTTTTTGAAGATATTTCCTTTTCCACAATAGGCCTCAAAGCTCTCCAAACATCCATTGCTAATTC

>C20F252

CTACAAAAAGAGTATTCCAAAACTGCTCAATCAAAAGAAACGTCTAAAACTGTGAGATGAATGCACACATCACAAAGTAGTTTCTCAGAATGCTTCAGTGTAGTTTTTATGTGAAGACATTAGCTTGTCCACGGAAGGTCTCAAAGCGCTCCAAATATCCACTTGCAGATTC

>C20F253

TACAAAGAGAGAGATTCTAAACTGCTCAATCAAAAGGTAGTTTCAACCCTGTGATATGAATGCACACATCCCAGAGAAGTTTCTCAAAATGCTTCTGTTTAGTTTTTATTTGAAGATATTTCCTTTTCCACCATACGCCTCAAAGGTCTCCAAATATCCACATGCTGCTTC

>C20F254

CTGCAAAAAGGGAGATTCAAAACTGTACAATCAAAAGATAAGTTCAAATATGTGAGTTGAATGCACACAAAACAAAGAAGTTTCTCAGAATACCTCTGTGTAGTTTTTAGGTGAATATATTTGATTTTCCACAGTAGGCCTCAAAGGGCTCCAAATATCCACTTTCAGATT

>C20F255

TTCAAAAAGATCGCTTCAACACTGCTCTATAAAAAGGAAGTTTCAACTCCATTAGATGACTGCACACATCACAAAGAAGTTTCTCAGAATGCCTCTGTGTAGTTTGTATGTGAAGATATTTCCTTTTCCACAGTAGGCCACAAAGGGCTCCAAATATCTACTTGCAGAAT

>C20F256

TACAAAAAGAGAGATTCAAAGCTGTTCAATCAAAAGATAGTTTCAACTCTGCGTGTTGAATGCACAAATCACAAAGTAGTTTCTCTGAATGCTTCTGTGTTGTGTTTGTTTATGTGAAGATATTTGCTTTTCCACTATAGGGCAAAATAGGGCTCCAAATATCCACTTGCAGATT

>C20F257

CTATGGAAAGAGTCTCAAAACTGCTCAGTCAAAATAAAGGTAGAACTCTGTGAGAAGAATGCACACATCACAAAGAAGTTTCTCAGAATATATTTGTGTAGTTTTTATTTGAGGATAGTTCCTTTTCTACCATAGGCTGCAAAGGGCTCCAAATATCCACTTGCAGATGG

>C20F258

CTTCAAAAAGAGTGTTTCAAAAGTTCTCAATCAAAAGAAAAGTTAAACTCCATGAGATGAATGCATACATCACAAAGAAAGTTCTGAGAATGCTTCTGTCTTGTTTTTATGTGAAGATATTTGATTCTCTCCTGTAGGCAACAAGGCGCTCCAAATATCCACTTTCAGATA

>C20F25

CTAGAAAAAGAGTGATTTAAAACTGCTCTATCAACAGAAAGGTTCGACTCTGTGAGTTGAATGCACTTATCACAAAGAAGTTTCTGAGAATGCTTCTGTCTAGTTTTTATGTGAAGATATTTCCTTTTCCACCATAAGCCTCAAAGCGCTCCAAATATCTACTTACAGATT

>C20F260

TGTGAAGaTATTTCCTTTTCCACCATAAGCCTCAAAGCACTCCAAATATCtACTTTCAGATTCTAGAAAAAGAGTGATTTAAAACTGCTCTATCAACAGAAAGGTTCACTCTGTGAGTTGAATtCACTTATCACAAAGAAGTTTCaGAGAATGCTTCTGTCTAGTTTTTA

>C20F266

TGTGAAGATATTTCCTTTTACACCATAGGCCTCAAACCGCTCCAAATATCCACTTGCAGATTCTCAAAAAGACTTTTTCAAAACTGCTCAATCAAAGGAAAGGTTCAACTCTGTGAGTTGAATGCACACAACACAAACAAGTTTCTGAGAATGCTGCTGTCTAGTTTTTA

>C20F274

TGTGCGGATATTTCCTTTTCCACCATAGGCATCAAAGCGCTCCAAATATCCAACTGCAGATTCTACAAAAAGAGTGTTTCAAAACTGCTCTATCAAAGAAAGGTTCAACTCTGTGAGTTGAATGCACACATCACAAAGACGTTTCTGAGAATGCTTCTGCTCTAGTTTTTT

>C20F27

CTACAAAAAGAGTTTCAAAACTGCTCTACAAAAGAAAGGTTCAACTCTGTGAGTTGAATTCACACATCACAAAGAAGTTTCAGAGAATGCTTCTGTCTAGTTTTTATGTGAAGATATTTCCTTTTACACCATAGGCCTCAAACCGCTCCAAATATCCACTTGCAGATT

>C20F280

TTGTGAAGGTGTTTCCTTTTCCACCATAGGCCTCAAAGCGCTCCAAATATCCACTTTCAGATTCCTGAAAAAGAGTGTTTTAAAACTCCTCTATCAACATAAGTGTTCAACTCTGTGAGTTGAATGCACTCATCACAAAGATATTTCTGAGAATGCTTCCGCCTAGTTTTTA

>C20F30

CCTGAAAAAGAGTGTTTTAAAACTCCTCTATCAACATAAGTGTTCAACTCTGTGAGTTGAATGCACTCATCACAAAGATATTTCTGAGAATGCTTCCGCCTAGTTTTTATGTGTAGGTATTTCCTTTTCCACCATAGGCCTCAAAGCACTCCAAATATCCACTTTCAGATT

>C20F39

AGTCTGCAAGTGCATATTTGGATAGCTTTGAGGATTTCATTGGAAACGGGAATATCTTCCCATAAAAACTAGAAAGAAGCATTCTCAGAAACTTCTTTGTGATGCTTGCATTCAACTCACAGAGTTGAACATTCCCTTTCATACAGCAGTTTGGAAACACTCTTTTTGTAG

>C20F41

GTTCTGAGTGACATTTGGAGTGCTTTCAGGCCTAGGTGAAAAAGGAAATATCTTCCCATAAAAACTAGACAGAAGCATTCTCAGAAACTTTTTGTGATGTGTGCCCTCAACTGACAGAGTTGAACCTTTCTTTTAAGAGCAGTTTTGAAACACTCTTTTTGTA

>C20F46

GTGTCTATAAGTGAACATTTGGCGTGCTTTCAGGCCTAACGTGAAAAAGGAAATATCTTCCCATAAAAACTAGACAGAAGCATTCTCAGAAACTTGTTTGTGATGTGTGCCCTCTACTGACAGAGTTGAACCTTTCTTTGCAAAGAGCAGTTTGAAACACTCTTTTTGTA

>C20F56

GTATCTGGATGAGGACATTTGGAGCGCTTTCAGGCGTATGGTGAAAAAGGAAATATCTTCCCGTAAAAACTAGACAGAAGCATTCTCAGAAGTTTATTTGTGATGTGTGCCCTCAACTAACAGAGTTGAACCTTTCTTTTGATAGAGCAGTTTTGAAACACTCTTTTTGTA

>C20F61

GTATCTGGATGTGGACATTTGGATCGCTTTCAGGCCTATGGTGAAAAAGGAAATATCTTCCCATGAAAACTAGACAGAAGCATTCTCAGAAACTTATTTGTGATGTGTGCCCTCAACTGACAGTGTTGAACCTTTGTTTTGATAGAGCAGTTCTGAAACACACTTTTTGTA

>C20F72

AATCTGCAAGTGGTTATTTTGATAGCTTTGAGGCTTTCATTGGAAACGGGAATATCTTCACATAAAAACTAGACAGAAGCATTCTCAGAAACTTCTTTATGAAGTTTGCATTCAACTCACAGAGTTGAACTTTCCGTTCCATACAGCAGTTTTGAAACTCTCTTTTTCTAG

>C20F74

AATCTTCAAGTGGATATTTGGATAGCTTTGAGGCTTTCATTGGAAACGGGAATATCTACACATAAGAACTAGACAGAAGCATTCTCAGAAAGTTCTTTATGAAGTTCGCATTCAACTCACAGAGTTGTACCTTCCTTTTCACACAGCAGTTTTGAGACATTCTTTTTGTAG

>C20F77

CTAAAAAAAGAGTGTTTCAAAACTGCTTATCAAAGAAACATTCAACTCTGTGAGATGAATGCACAGATCACAAAGAAGTTTCTCAGAATGCTTCTGTGTAGTTTTTATGTGAAGATATTTGATTTTCCACAGTAGGCCCCAAGAGCTCCAAATATCCACTCGCAGATT

>C20F81

CTAAAAAAAGAGTGCTTCTAAACTTCTGTACCAAAAGAAAGATTCAACACTGTGAGATGAATGCACAGATCACAAAGAAGTTCCTCAGAATGCTTCTGTGTAGTCTTTATGTGAAGATATTTGTTTTTCCACAGTAGTCCCCAATGAGCTCCAAATATCCACTTGCAGATT

>C20F82

CTAAAAAAAAAGTGTTTCAAAACTGCTATATCAATAGAAACATCCAACTCTGTGAGATGAATGCACAGATCACAAAGAAGTTTCTCAGAATGCTTCTGTGTAGTTTTTATGTGAAGATATTTGATTTTCCACAGTAGGCCCCAACGAGCTCCAAATATCCACTTGCAGATT

>C20F85

CTAAAAAAATAGTGTTTCAAAACTGCTGTATCAAAAGAAAGATTCAACTCTGTGAGATGGATGCACAGATCACAAAGAAGTTTCTCATAAAGCTTCTGTGTAGTTTTTATGTGAAGATATTTGTTTTTCCACAGCAGGCCCCAATGAACTCCAAATATCCACTTGCAGATT

>C20F90

CTACAAAAAGAGTGTTTCAAAACTGCTCAATCAACAGAGACATTCAACTCTGTGAGATGAATGCACCCATCACAAAGAAGTTTCTCAGAATGCTTCTGTGTAGTTTTTGTGTGAAGATATTTCATTTTCCACAGTACGCCTCAAAGCGCTCCAAATATCCACTCTCAGATT

>C20F95

CTACAAAAAGAGTGTTTCAAAACTGCTCAATCCAAAGAAAGGTTCTACTCTGTGAGATGAATGCACACATCACAAAGTAGTTTCTCAGAATGCTTCTGCATAGTTTTTATGTGAAGATATTTCCTTCTCCACTATAGGCCTCAAAAGGCTCCAAATATCCACTTGCAGATT

>C20F98

CTAAAAAAAGAGTGTTTCAAAACTGCTCAATCCAAAGAAAGGTTCTACTCTGTGAGATGAATGCACACATCACAAAGTAGTTTCTCAGAATGCTTCTGCATAGTTTTTATGTGAAGATATTTCCTTCTCCACTATAGGCCTCAAAAGGCTCCAAATATCCACTTGCGGATT

>C21F11

CTACAAAAAGAGTTTCAAACTGCTCAATCAAAAGAAATTCAACTCTTGAGATGAATGCACACATCACAAAGAAGTTTCTCAGAATCTTCTTTAGTTTTTATGTGAAGATATTTCCTTTTACATAGGCCCAAAGTAAATATCCTTGCAGAT

>C21F12

CTACAAAAAGAGTTTCAAACTGCTCAATCAAAAGAAATTCAACTCTGTGAGATGAAGCACACATCACAAAGAAGTTTCTCAGAATTTCTGTGTAGTTTTATGTGAAGATATTTCTTTTACATAGGCCTCAAACCTCAAATTCCATTTGCAGAT

>C21F36

CGCTAAAAGAGTGTTTCAAAACTGCTCAATCAAAAGAAAGGTTCTAGTCGGTGAGATGAATGCACACATCACAAAGAAGTTTCTATGAATGCTTCTGTCTGATTTATATTGAAGATATTTCCTTTTTCACCGTAGGCCTCAGAGTGCTTAAAATATCCATTTGCAGATAC

>C21F37

CTGCAGAAAGAGTTTTTCAAAGCTGCTCAATCAAAAGAAAAGTTCAACTCTTTGAGATGAATGCACACATCATGAAGTTCCTCAGAATGCTTCTATTTTTATGTGAAGATATATCCTTTTCTACCATAGACCACAAAACGCTCCAAATATCCCCTTGCAGTTTC

>C21F38

TAGAAAAGACTGTTTCCAAACTGCTCAATCAAAATAAAGTTCAACTCAGTGAGATGAATGCACACATCACCAAGACGTTTCTGAGAAAGATTCTGTCTCGTTTTTATGTGAAGATATTTCCTGTTTCCCCAGAGGCATCAATGGGCTCACAAATATTCCTTTGCATATTC

>C21F46

TACAAAAAGAGTGTTTCAAACCTGCTCTATGAAAGGCCATGTTCATCTCTATGAGTTGAATGGAAATATCCGAAAGAAATTTCTGGGAATGCTGCTGTCTAGTGTTTATACGAATTCCCGCTTCCAACGAAATCCTCAAAGCAATCCAAATATCCACTTGCAGAATC

>C21F51

CACAAAAAGAGTGTTTCAAAACTGCTCTATCAATAGAAAGGTTCAACTCTTTTAGTTGAGTACACACATCACGAACAAGTTTCTGAGAATGCTTCTGTCTGGCTTTTATTGGAAGACGTTTCCTTTTCACCAAAGGCATCAAAGCGCTCCAAATGTCCACTTCCAGATTC

>C21F53

CACAAAAAGAGTGTTTCAAAACTGCTCTATCAATAGAAATGTTCAACTCCTTTGGCTGGGTACACACATCACAAACAAGTTTCTGAGAATGCTTCTGTCTAGTTTTTATGGGAAGACGTTCCCTTTTTCACCAAAGGCATCAAAGCGCTCCAAATGTCCACTTCCAGACAC

>C21F56

CCACAAAAATAGAGTTTCAAAGCTGCTCTGTAAAAAGAAAGGTTCCACTCTGTTAGCTGAGTACACACATCACAAACTTGTTTCTGAGAATCCTTCTGTCTCGTTTTTATGGGAAGATATTTACTTTTCCACCGTAGGCATCAAAGCGCTCCAAATGTCCACATCCAGATAC

>C21F59

CTACAAAAAGAGTGTTTCCAAACTGCTGCATCAAAAGAGAGGTTCCACTCTGTTAGCTGAGTACACACATCACAAACTTGTTTCTCAGAATCCTTCTGTCTCGTTTTTATGGGAAGATATTTACTTTTTCACCGTAGGCATCAAAGCGCTCCAAATGTCCACATCCAGATAC

>C21F64

TACAAAATGACTGTTTAGAAGGTGCTCAATCAAAAAAAAAGTTCAACAGTGTGAGATGAATGCGCCCATTCAAAGGAAGTTTCTCAGAATTCTTCTATCTAGTTTTTATGTGAAGATATTTCCTTTTTCACTATAGGCCACAAAGTGCTCCAAATATCCACTTGCAGACTC

>C21F65

CTACAAAAAGACTGTTTCCAAACTGCTCAATCAACAGAAAGTTTCAACCCGGTGAGTAGAAGTCACACATGACAAAATAGTTTCTCAGAAAGTATCTGTCTAGTTTTTATGTGAAGATATTTCCTATCACCCCAGAAGCCTCAATGGGCTCACAAATATTCCTTTGCAGATT

>C21F69

TTCCAAAAGAGTGTTTCAAACGTGCTCAAAGTAAGGGAATGTTCAACTCTGTGACTTGAATGCAGATATCACCAAGTAGTTTCTAATAGTGCTTCTGTCTAGATTTTAGATGATGATATTCCCGTTTCCAACGAAATCGTTAGAGCTATCCAAATATCCAGTTACAGTTT

>C21F72

TACAAAAAGAGTGTTTCAAACGTGCTCTAAGAAAGCGAATGTTCAACTCTGTGACTTGAATGCAGATATCACAAAGTAGTTTCTGAGAGGGCTTCTGTCTAGATTTTAGATGATGATATTCCCGTTTCCAACGAAATCATTAGAGCTATCCAAATATCCACTTACAGTTT

>C21F74

TACAAAAAGACTGTTCCCAAACTGCTCAATAAAAAGAAAGTTTCAACTCTATGAGATAAAAGCAAATATCACAAAGAAGTTTCTCAGAAACTTTCTATCTAGTTTTTATGTGAACATATTTCTTATCACCCCATAGACCTCAATCGGCTCACAAGTATCCTTCTGCAGATTG

>C21F75

CTCCAAAAAGAGTGTTTTAAAACTGCTGTACCAAAGAAAGTTTCATGTCTGAGATATGACTGCATACAACACAGAGAAgTTTCTCAAAGTGCTTCTGTTTATTTTTtTTATGAAGATATTTCCTTTTCCACTATGGGCCACAGAGCGCTCCAAATATCCACTGGCAGATTC

>C21F76

TACTAAATGACTGTATCGAAGCTGCTCAATCAAAAGACGGGTTTAACAGTGTGAGACGAAAATACACCTTCCTAGGAAGTTTCTCAGAATTCTTCTTTCTAGTTTTTTATGTGAAGATATTTCCTTTTCCACTATAGGCCTCAAAGCGTTCCAAATATCCACTTGCAGATAC

>C21F77

CTACCAAAAGGGTGTTTCCAAATTGCTGCATCAAAAGAAAGGTTCAACTCTGTTAGTTGAGGACACACATCACAAAGAAGTTTGTGAGAATGCTTCTGTCTAGATTTTGTATGACGATATTCCCTTTTCCAACGATATCGTTAAAGCAATCTAAATATCAATTTGCAGAAT

>C14F65C22F103

TACAAAAAGAGTGTATCAAAACTGCTCTGTCAAAAGGAAGGTTCTTCTCTGTTAGGTGAGTGCATACGTCATAAAGGAGTTTCTGAGAATGTTTCTGTCTAGTGGTTATGGGAAGATATTTGCTTTTTCACCGTAGGCCTCAGAGCGCTCCAAATATCCACTTGCACATAC

>C22F106

TACAAAAAGAGTGTATCAAAAATGCTCTGTCAAAAGGAAAGTTCTTCTCTGCTAGTTGAGTACATACGTCATAAAGAAGTTTCTGAGAATGTTCCTGTCTAGTGGTTATGGGAAGATATTTGCTTTTTCCCCGTAGGCCTCAAAGCGCTCCAAATGTCCACTTGCACATAC

>C22F107

CTACAGAAAGAGTGTTTCAAAACAGTTCTATCAAAAGAAAGGTTCAACTCTGTTAGTTGAGTTCACACATCcGAAACTAGTTTCTGAGAATTCTTCTGTCTAGTTAATTTGGGAAGATATTTCCATTTTCACCGAAGGTCTCAAAGCGCTCCAAATGTCCACTTCCAGA

>C22F108

TTATAAAAAGAGAGATTCAAAACTGCTCAATGAGAAAATAAGTTAAACTCTGTGGGTTGAGTGCACACCTCACAGAGAAGTTTCTCAGAATGCTTCTGTGTAGTTTTTATGTGAAGATATTTGCTTTTCCATAATAGGTTTCAAAGCTCTCCAAACATCCACTTGCAGATT

>C22F109

TGCAAAAAGAGAGATTCAAAACTGCTCATTCTTAAGATAGGTTCAAGTCTGTGAGTTGAATGCATACATCACAAAGAAGTTTATCAGAATGCTCCTGAATAGTTTTTATGTGAAGATATTTACTTTTCCACAATAGCCCTCAAAGGGCTCCAAATATCCAGTTGCAGATTC

>C22F111

CTACAAAAAGACTGTTTCCAAACTGGCCCATATAGCATGTTTCAACTATGTGAAATGAATGCACTCATCAAAAAGAAGTTTCTCAGGATTCTCCTGTCTAGTTTTTATGTGAAGATATTTCCTTTTTCACCGTAGGCCACAAATTGCTCCAAATATCCATTTGCAGATT

>C22F117

CTACAAAAAGACTGATTCCAAACTGCTCAATCAGAAGAAGGGTTCAATTCCGTGTGACAAACGTGCACATCACCAAGAAATTTGTCAGAAAGCTTCTGTCTACTTTTTATGTGAAGATATTTCATATTTCAACAAAGGCCATAAAGGGCTCACAAATATCCCTTCGCAGATT

>C22F121

TACAGAAAGACTTTCCAAACTGCTCAATCAAAAGAAAGGTTCAACACTGTGAGATGAAGGCACACATCACCAAGAAGTTTCTCAGAAACCTTCTGTCTAGTTTTTAGGTGAAGATACTTCGTATTTCACCACAGGCCATAAAGGGCTCACAAATATCCCTTTGCAGGTT

>C22F127

TACCAAAGACTGTTTCCAAACTGCTCAACTGAAAGAAAGGTTGAATTCTGTGACATGAATTCACACATCACAAAGAGGTTTTTCAGAAATCTTCTGTCTGGTTTTTAGGTGAAGATACTTCCTTTTTCACCACGGGCCTCAAATATCTCCAAATATCCATTTGCAGATTC

>C22F130

TGCCAAAAGAGAAATTCAAAACTGCTAAATCAAAAGATATGTTCAGCCCTGTGAGTTGAATGCACACATCACAAATAAGTTTCTGAGAATGTTTCTTTGTAGTTCTTATTTGAAGATATTTCCTTTTCTACCATAGCCCTCAAAGGGCTCCAATTATTCACTTGCAGATT

>C22F131

GTACAAAAAGAGTGTTTCAAAACTGCTCCATCAAAAGAAAGTTTCAACTCTATGACATGAATGCACGCACCACAAAGAACTTTCCCAGAATATTTATGTGTAGTTTTTATGTGAAGACATTTCCTTTTCCACAATAGGCCACAAAGCTTTGCAAACATACACTTGCAGATT

>C22F132

CTACAAAAAGAGTGTTTCAAAACTGCTCAATCAAAGGAAAGTTTCAACTCTGTGAGACGAATGCACACATCACTAAGAAGTTTCTCAGAATGCTTCTGTTTAGTTTTTGTTTGTAGGTATATGCTTTTCCACGGTAGGCCTCAATTCCCTCTAAATATCCACTTGCAGAcT

>C22F133

CTACAAAAACAGTGTTTCAAAACTGCTCAATAAAACGGTAGGTTCAAACCTGTGAGATAAATGCACATATCACAAAGAAGTTTCTCAGAATTCTTCTGTGTAATTTTTATCTGAAGATATTTCCTTTTCCACCATAGGACACAATGGGCTCCAAATATCCACTTGTACATT

>C22F134

TACCACAGAAAGAGTGTTTCAAACCTGCTCTATCAAATGGAATGTTCAACTCTGTAACTTGAAAGCAATCATTACAAGGAAGTTACTGAGAATCTTTCTGTGTAGATTTTATATGAAGATATTCCCGTTTCCAAGGAAATCGTGAAAGCTATCCAAATATCCACTTGCAGGTTC

>C22F135

TACAAAAAGtGTGTTTCAAACCTTCTGTATGAAATGTAATGTTCAACTCTGTGACTTGAATGCAAACAGCAGTAGGAAGTTTCTGAGAATCCTTCTGTGTAGATTTTATGTGAAGATATTCCCGTTTCTAACGAAATCGTTAAAGCTATCCAAATATGCACTTGCAGGTT

>C22F137

CTACAAAAAGACTGTATCCAAACTGCTCAATAAAAAGAAAGTTTTAACTCTGTTAGATTAATGGACACATCAAAAAGTAGTTTCTCAGAAAACTTCTGTGTAGTTTTTATGTGAAGATATTTCCTTTGTCACCATTGGCCTCAAAGCACTCCTAATATCCATTTACAGATGT

>C22F13

CTAGAAAAAGAGTGTTTCCAAACTCCTCAATCAAAGGATAGTTTCAATTCTGTGAGAGAAAGCACACATCACAACGAAGTTTCTTAGAAAGCGTCTGTGTAGTTTTTATGTGAAGATACTTCACATTGCATCACAGTACTCAATGGGCTCAGAAATATCCCCTTGCAGATC

>C22F140

GCAGAGAAAGACTGTCTCTAAACTGCTCAAATAAAATAAAGTTTCAACACGGTGAGATGAATGCACACATCACAAAGAAGTTCCTCAGAAAGCTTCTGTCTGGTTTTTATGTGAAGATATTTCCTTTTTCACCATAGGCCTTACACCGCTCACAAATATCCTTCTGCAGATAC

>C14F71C22F148

CACAGAAAGACTGTTTCAAAACTGCTCTGTCAATAGAAAGGTTCAACTCTGTTAGCTGCGTGCATATATCCCAAAGAAGATTCTGAGATTGCTTCTGTCTAGTTTTTATGGGAAGATATTTCCCTTTTCACCGTAGGCGTCAAGGCGCTCCAAATGTCCACTTCCAGATAC

>C22F151

TAGAAAAAGACTGTTTCCAAACTGCTCCATCAAAAGAAAATTTCACCTATCTGAGATGAATGCACACATCATAAAGAAGTTCCTCAGAATTCTTCTGTCTAGTTTTTATGTGAAGATGTTTCCATTTTCACCTTAGGCCACAAAGCGCTCCAAACATCCTTTGCAGATGA

>C22F156

CTAAGAAAAGACGTTTTCCAAACTCCTCAATCAAAAGAAAGGTTTAACTCTGTGAGATGAATGGACACATCACGAAGAAGTTTCTCAGAAAGCTTCTGTCTAGTTTTTCTGTGAAGATATTTCTTTTTCACCATAGGCCTCAAGCAGCTAAGAAATTTCCCTCTGCAGCTTC

>C22F15

CTAGAAAAAGAGTGTTTCCAAACTCCTCAATCAAAAGAAAGTTTCAATTCTGTGAGATGAAAGCACACATCACACCGAAGTTTCTTAGAAAGCTTCTGTCTAGTTTTTATGTGAAGATACTTCACAGTGCATCATAGTACTCAATGGGCTCAGAAATATCCCTTGGCAGATT

>C22F160

CTACAAAAAGAGTGTTTCAAAACTGCTCCATCAAGAGAAAGTTTTAACTCTGTGAGATGAATGCAAACATCACAAAGATGTTTCTCTGAATGCTTCTGTGTAGTTTTAATCTGAAGATAATTGCTTTTCCACGGTAGGCCTTAAGGCCCTCAAAATATCCAGTTGCAGATTC

>C22F161

CTACAAAAAGAGAGACGCAAAACTGCTCAAAGAGGACATATGTTCAACTCTGTGAGTTGAATGCACACGACACAAAGAAGTTTCTCAGAATGGTTCTGTGTAGTTTTTATGTGAAAATATTTCCCTTTCCACAATATGCCTGAAAGCTCTCCAAACATCCCCTTGCAGATTC

>C22F163

CTACAAAAGGACTGTTTCAAAACTGCTCAATCCAAAGAAAGTTTCAACTATGTGAGATGAATGCACACGTCACGAAGAAGTTCCTCAGAATGCTTCTGTCTAGTTTATATGTGAAGAAGATTCCTATTTCACCATAGGCAATAAAGGGCTCACAAATATTTTTTGCAGATT

>C22F167

CTACAAAAATATGCTTTCCAAAGTGCTGAATTAAAAGAAACCTTCAACTCTGTCAGATGAATGGAGACATCACAAAGAAGTTCCTCAGAATGCTTCTGTCTAGTTTAAATGTGAAGATATTTCTTTTTCACCATAGACCTCAAAGGGCTCAGAATTAGACCTTTGCAGATT

>C22F171

ACTACCAAGAGACTTTCTCCAAATTGCTAAATCAAAAGAAAGGTTCAACTCTGTGAGATGAATACACACATCAAAAAGAAGTTTCTCAAAATGCTTCTGTCTAGTTTTCATGGGAAGATATTTATTTTTCACCGTTGGCCCCAAACCGCTCAGAAATATCCCTTTGCAGTTTG

>C22F179

AACAAAAAGTGTTTTTCAGAACTGCTCTATCAAAAGAAAGATCCACCTCTGTTAGCTGAGTTCACACATCACAAACAAGTTTATGAGAATGCTTCTGTCTAGTTTTTATTTGAAGATATTTCCTTTCTCACCATAGACCTGAAAGCTGTCCTAATGTTCACTTCCAGATAC

>C22F182

AACAGTGTTTTTCAGAACTGCTCTATCAAAAGAAAGATCCACCTCTGATAGCTGAGTTCACACTTCACAAACAAGTTTATCAGAATGCTTCTGTCTAGTTTTTATTTGAAGATATATCCTTTCTCACTATAGACCTGAAAGCTCTCCTAAAGTTCACTTCCAGATAC

>C22F183

AACAAAGTGTTTTTCAAAACTGCTGTATCAAAAGAAAGATCCACCTCTGTTAGCTGAGTTCACACTTCACAAACAAGTTTATCAGAATTCTTCTGTCTATTTTTTATTTGAAGATATAGCCTTTCTCACTATAGACATGAAAGCTCTCCTAAAGTTCACTTCCAGATACT

>C22F186

CTACAAAAAGAATGTTCCCAAACTGGTCAATCAAAAGAAAGGCGCAACTCTGTGAGACGAAAGCACACATCACAAAGAAGTTTCTCGGAAAGCCTCTGTCTACATTTTATGTGAAGGTATTTCCTTTGGCACCATAGGCCTTAAACCGCTCGCAAATATAACTCCACTTA

>C22F189

CTACAGAAAGAATGTTTCCAAACTGGTCAATCAACAGAAAGGCTCAACTCTGTGAGACGAAAGCACACATCACAAAGAAGTTTCTCAGAAAGCTTCTGTCTACATTTTATGTGAAGGTATTTCCTTTGGCACCGTAGGCCTTAAACCACTCACAAACATAACTCCGCTTATA

>C22F194

CTAAAAGACCACCATTTCCATACTTCTCAATCAAAAGAAAGGTTAAATTCTGTGAGGTTAATGCACACATCAGAATGAAGTTTCTCAGAATTCTCCTGTCTAGTTTTCATGTGAAGATATTTACTATTTCACTATAGGCTTCAAATGTCTCAAAAATATCCCTTTGCAGATT

>C22F199

TAGAAAAAGACTGCTTCCAAACTGCTCAATGAAAGGAAATGGTCAACTATTAGAGATGAATGGAAATGTCACAAAGAGTTTTCTCAAAAAGCTACTGTGTCGTTTTTATGTGAAGACATTGCCTTTTGCACCCTAGGCCTTAAAACTCTCTAAATACACATTCACAGATT

>C22F21

CACAGAAAGAGTGTTTCCAAACTGCTCAATCAAAAGAAAGTGTTTAACTCTGTGAGGTGAAAGCACACATCTCAAAGAAGTTTCTCCGAAAGCTTCGGTCTAGTTTTCATGTGATGATATTTCCAGTCTCACCATAGGCCTCAAAGGGCTAAGAAATATCCCTTTCCAGATT

>C22F26

CTACAAAAAGACTGTTTCCAAACTGCTCAATCAAAGGAGAGGTTCAACTCTGTGACGTGAATGGACACATCACAAAAAATTTCTTGGAATGCTTCCGTCTAGTTTTTATGGGAAGATATTTCTCTTTCACCATAAGCCTCAAACGGATCAGAATTCTCCCTTTGCAGATT

>C22F31

TACGAAAAGACTGTTTCCAAACTGCTCAATCAAAAGAAATTTTCAACTCTGTGAGATGAAAGCACACATCACAAAAAAGTTTCTCAGAAATCTTCTGTCTCGCTTTTATCTCAAGATAATTCCTATTTTGCCATAGGAATCAAGGGGCTCACATATATCCCTTTGCAGATTC

>C22F34

TACAAAAGTTCTCTTTACAAACTTCTCAATCAAAAGAAACGTTCAACATTGTGAGATGAATGAACACATCCCAAAGAAGTTTCTCAGGTTGCTTCTGTCTGGTTGCTATGTGAAGATGTTTCCTTTTTCACCATAGTCTTTAAGCCACTCAAAAATATCTGTCTGCAGACT

>C22F40

GTACAATAAGCCTCTTTCCAATCTGCTCAATCAAAAGAAAGTTTCAACTCTGTGAGGTGAATGCACACATCACAAGGGAGTTTCTCAGAAAGCTTCTGTTTAGTTTTTACGTGAAGATGTTTCGTTTTTCAACATGGGCCTCAAAAGCTCTCCAAATATCCATTTGCAGATT

>C22F44

GTACGATAAGCCTCTTTCCAATCTGCTCAATCAAAAGAAAGTTTCCACTCGGTGAGGTGAATGCACACATCGCAAGGGAGTTTCTCAGAAAGCTTCTGTTTAGTTTTTACGTGAAGATATTTCGTTTTTCACCACGGGCCTCAAAAGCTCTCCAAATATCCATTTGCAGATT

>C22F46

TACAAAAAGAGTGTTTCAAACTGCTCTATGAAAGGGAATGTTCAACTCTGTGACTTGAATGCAACATCACAAAGAAGTTTCTGAGAATGCTTCTGTCTAGATTTTATATGAAGATATTCCCGTTTCCAACGAAATCCTCAAACTATCCAAATATCCACTTGCAGATTC

>C22F47

CTACAAAAAGAGTGTTTCAAAACTGCTCATCAAAAGAAAGGTTCAACTCTGTGAGTGAATGCACACATCACAAAGAAGTTTCTAGAATGCTTCTGTTAGTTTTTATGTGAAGATATTTCCTTTTCCACCATAGGCCTCAAAGCTCCAAATTCCACTTGCAGATT

>C22F59

TACAAAAAGAGTGTTTCAAACCTGCTCTATGAAAGGGAATGTTCAACTCTGTGACTTGAATGCAACATCACAAAGAAGTTTCTGAGAATGCTTCTGTCTAGATTTTATATGAAGATATTCCCGTTTCCAACGAAATCTCAAATCTATCCAAATATCCACTTGCAGATTC

>C14F25C22F65

TACAAAAAGAGCGTTTCAAACCTGCTCTATGAAAGGCAATGTTCAACTCTGTGACTTGAATGCAGACATCACAGAGCAGTTTCTGAGAATGCTTCTGTCTAGATTTTATAGGAAGATATTCCCGTTTCCAACGAAATCTTCACAGCTATCCAAATATCCACTTGCAGATTC

>C22F68

TACAAAAAGAGCGTTTCAAACCTGCTCTATGAAAGGCAATGTTCAACTCTGTGACTTGAATGCAGACATCACAGAGCAGTTTCTGAGAATGCTTCTGTCTAGGTTTTATAGGAAGATATTCCCTTTTCCAACGAAATCTTCCAAGCTATCCAAATATCCACTTGCAGATTC

>C22F71

TACAAAAAGAGTGTTTCAAACCTACTCTGTGAAAGGGAATATTCAACTCTGTGACTTGAATGCACATATCACAAAGAAGTTTCTGAGAATGCTTCTGTCGAGATTTTATATGAAGATATTCCCGTTTCCAACGAAATCCTGAAATCTATCCAAATATCCCCTCGCAGATT

>C22F77

TACAAAAAGAGTGCCTCAAAGCTGCTCTCTGAAACGGAATGTTCAACTCTATGAGTTGAATGCAAACATCACAAAGACGTTTCTGAGAATGCTTCTGTCTAGATTTGATATGAAGATATTCCCGTTTCCAACGAAATCTTCAAATCTATCCAAATGTCCACTTGCAGATTC

>C22F82

TACAAAAAGAGTGCCTCAAAGCTGCTCTCTGAAATGGAATGTTCAACTCTATGAGTTGAATGCAAACATCACAAAGACGTTTCTGAGAATGCTTCTGTCTAGATTTGATATGAAGATATTCCCGTTTCCAACGAAATCTTCAAATCTATCCAAATGTCCACTTCTATAAT

>C14F44C22F87

TACAGAAAGAGTGTTTCAAAACTGCTGTACGAAAGGGAATGTTCAACTCTGTGACTTGAATGCACACATCACAAAGAAGTTTCTGAGGATGCTGCTGTCTACTTTTTATACGTAATCCCGTTTCCAACGAAATCCTCCAAGCTATCCAAATATCCACTTGCAGATTC

>C22F91

CTGCAAAAAGAAAGATTCAAAAATCCTCAATCAAAAGATAGGTTCAACTCTTGTGAGTTGAATGCACACATTGCAAAGAAGTTTCTCAGAATACTTCTATGTAGTTTTCATGGGAAGATATTTCCTTTTCCACAACAGGCCTCAAAGGGCTCCAAATATCCACTTGCAGATT

>C22F92

CTACAAATAGAGAGATTCAAAACTGCTCAATGAGAAGAAAAGTTTAACTCTGTGGGTTGAATGGACTCCTCATAAAGAAGTTTCTCAGAATGCTTCTGTGTAGTTTTTATGTGAAGATATTTCCTTTTCCACAATATTCCTCAAAGGGCTCCAAATATCCAGTTGCAGATT

>C14F56C22F96

CTACAAAAAGAGTGTTTCAAAACTGCTCTGTAAAAAGAAAGGTTCAACTCTGTTAGTTGAGTACACACATCACAAACAAGTTTCACAGAATGCTTCTTTCTAGCTTGTAGGGGAAGATATTCCCTTTATCACCATGGGCCTCAAACCGTCCGAAACGTCCACTTCCATATAC

>C2F11

AATTTGCATGTGGATATTTGGACAGCTTTGAGGATTTCGTTGGAAACGGGAAAATCTTCATATGAAATCGAGACACAAGCATTCTCAGAAACCTCTTTGGGATGTTAGCaTTCGAGTCACAGAGTTGAACATTCCCTTTCATAGAGCAGGTTTGAAGCACTCTTTTTGTAG

>C2F12

GAATCTGCAAGTGGATATTTGGATAGCTTAGAGGATTTCGTTGGAAACGGGAATATGTCCATAAAAACCTAGACAGAAGCATTCTCAGAAAAATCTCTGTGAGGATTGCATTCAAGTCCCAGAGTTGAACATTCCCTTTCATAGAGCAGGTGTGAACACAAGATTTTGTAGTA

>C2F19

GTATCTGGATGTGGACATTTGGAGCGCTTTCAGGCCTATGGTGAAAAAGGAAATATCTTCCCCTGAAAACTAGACAGAAGCATTCTCAGAAACTTATTTGTGATGTGCGCCCTCAACTAACAGTGTTGAACCTTTCTTTTGATAGAGCAGTTTTGAAACACTCTTTTTGT

>C2F20

GTATCTGGATGTGGACATTTGGAGCGCTTTCAGGCCTATGGTGAAAAAGGAAATATCTTCCCCTGAAAACTAGACAGAAGCATTCTCAGAAACTTATTTGTGATGTGCGCCCTCAACTAACAGTGTTGAAGCTTTCTTTTGATAGAGCAGTTTTGAAACACTCTTTTTGTA

>C2F23

TATCTGGAAGTGGACATTTTGATCGCTTTGAGGCGTAAGGTGAAAAAGGAAATATCTTGCCACAAAAACTACACAGAAGCATTCTCAGAAACTACGTTGTGATGTGTTTACTCAACTAACAGAGTTGAACCTTTCTTTTGATAGAGCAGTTTTGAAACACTCTTTTTGTAG

>C2F27

AATCTGGAAGTGGACATTTGGATCGCTTTGAGGCCTGGTGAAAAAGGGAATATCTTCGAATAAAAACTAGACAGAAGCATTCTCATAAACTAGTTTGTGATGTGTGTGCTTAACTAACAGAGCTGAACCTTTCTTTTCATAGAGCAGTTTTGAAACACTCTTTTTGTAG

>C2F28

AATCTGGAAGTGGACATTTGGATCGCTTTGAGGCCTGCGGTGAAAAAGGTATATCTTCGCATAAAAACTAGACAGAAGTATTCTCATGAACTAGTTTGTGATGTGTGTGCTCAACTACCAGAGTTGAACCTTTCTTTTGATAGAGCAGTTTTgAAACACTCTTTTTGTgG

>C2F30

TATGGAACTGGACATTTGGAGTGCTTTGTGACCTATTGTGAAAAAGGAAATATCTTCCCATATAAACTAGGCAGAAGCATTCTCAGAAACCAGTTTGTGATGTGTGTACTCAACTAACAGGGTTGAACCTTTCTTTTGAGAGAGCACTCTTGAAACACTCTTTTTGTA

>C2F33

GTATCTGGAAGTGGACATTTGGAGCGCTCTCAGGACTACGGTGAAAAAGGAAATATCTTCCAATAAAAGCTAGATAGAAGCAATGTCAGAAACTTTTTCATGATGTATCTACTCAGCTAACAGAGTTGAACCTTTCTTTTGAGAGAGCAGTTTTGAAACACTCTTTTTGTG

>C2F38

GTATCTGGAAATGGACATTTAGATCTCTTTGAGGTCTATGGTGAAAAAGGAAATATCTTCGCATAAAAACTAGACGGAAGCAGTCTCCAAAACTTGTTTGGAATGTGTGTACTCAACTAACAGAGTTGAATCTTTCTTTTGATAGAGCAGTTTTGAAACACTCTTTTTGTA

>C2F3

AATCTGCAAGTGGATATTTGATAGCTTTGAGGATTTCGTTGGAAACGGGATTATATAAAAAGCAGACAGCAGCATTCCCAGAACTTCTTTGTGATGTTTGCATTCAAGTCACAGAGTTGAACATTCCCTTTCATAGAGCAGGTTTGAAACACTCTTTTTGTA

>C2F8

ATCTGCAAGGGATATTTGGAAAGCTTTGAGGATTTCGTTGGAAACGGGAATATCTTCATATAAAATCTAGACAGAAGCATTCTCAGAAACATCTTTGGGATGTTTGCATTCAAGTCACAGAGTTGAACATTCCCTTTCATGGAGCAGGTTTGAAACACTCTTTTTGTGG

>C2F9

AATCTGCAGGTGGATATTTGGATAGCTTTGAGGATTTCCTTGGAAAAGGGAATATCTTCATATAAAATCTAGACAGAAGCATTCGCAGAATCACCTTTGTGATGTTTGCATTGAAGTCAGAGAGTTGtACATTCCCTTTCATAGAGCAGCTTTGAAACACTCTTTTTGTA

>C3F117

CACAGACAAGTGTTTCAAATCTGCACTGTCTAAAGGAAGGTTCAACCCTGTGAGTTGAATACACACACACAGAAAAAAATTCACTGAGAATTCTATTGTCTATCATTACACGAAGAAATCCCGTTTACTACGAAGGCCTCAAAGAGGTCCAAATATCCAGCTGCAGACAT

>C3F120

TGCAAAAAGAGTGTTTCGAAACAACTGTATGAAAAGAAAGGTTAAACACTGTGAGTTGAACGCACACATTGCAAAGCAGTTTCTGAGAATGATTCCGTCTAATTATTATACGAAGGTATTTCCTTTTCTATCATTGGCCTCAAAGCGCTTGATACCTCCACCTGAAAATTC

>C3F125

CACAAAAAGAGTGTTTCCAAACTGCTCTATCAAAGGAAGTTTAAACTCTGTAGCTAATGCAAGCATCACAAAACAGCTTCGGAGAATGAATCTGCCTAGTTTTTCTGTGAAGATATTTCTTTTTCTGCCATAGACCTCAAACCGCTGTAAAAATCCACTTGGAAATTC

>C3F132

CCACAAAAAGAGTGTTTAAAACCGCTCTATCCAAAGAAAGGTTAAACTCTGTCAGCTGAATGCGCACATCACAAAGTAGCTTCAGAGAACAATTATGTCTAGTTTTTCTGTGAAGATATTTCTCTTCTACATAGGCCTGAAACCGCTCTAAATATTCACTTGGAAATT

>C3F142

CAACAAAAAGAGTTTTTCAAAACTGCTCCATCAAGAGGAATATTCAACTCTGAGAGTTGAAGGCAGGTATCACAAAGTAGTTCCCGACAATGCTTCTGTCTAGATTTTATGTGAAGACATTCCCTTTTGTACCACAGGCCTGAAAGCACTCTAAATATAGAATTGCAAATT

>C3F149

TACAAAGAGACTGTTTCATAACTGCTCTATAGGAAGAAAGGTTCAACTCTGTGAGTTGAATGCAGAGATCACAACGTGGTTTCTGCGAATGATTCTTTGTAGTTTTTACATGAAGATATTTCGTTGTCAACCGTAGGCTTCAAAGCACTCAAAGTATTCACTTGGAACTTT

>C3F58

TGCAAAAAGAGTGTTTCAAAACCGCTCCATTAAAAGGAATGTTGAACTCTGTGAGTTGAATGCAAACATCACAACTCAGTTTCTGAGAATGCTTCTGACTAGATTTTATGGTAAGATATTTCCTTTTCTACCGTAGGCTTCAATGCCCTCTAAATACACCCTTGCAAATTC

>C3F61

CACAAAAAGAGTGTTTCCAATCTGCTCTGTCTAAAGGAAGGTTCAACTCTGTGAGTTGAATACACACACACAAAAAGAAGTTACTGAGAATTCTTCTGTCTAGCATTATATGAAGAAATCCCGTTTCCAACGAAGGCCTCAAAGAGGTCCAAATATCCACTTGCAGATTC

>C3F62

CACAAAAAGAGTGTTTCCAATCTGCTCTGTCTAAAGGAAGGTTCAACTCTGTGAGTTGAATACACACACACAAAGAAGCTACTGAGAATTCTTTTGTCAAGAATTATAAGAAGAAATCCCGTTTCCAACGAAGGCCTCAAAGAGTTCCAAATATCCACTTGCACACT

>C3F66

CACAAAAAGAGGTTTCCAATCTGCTCTGTCTAAAGGAAGGTTCAACTCTGTGAGTTGAATACACACACACAAAAAGAAGTTACTGAGAATTCTTCTGTCTAGCATTATATGAAGAAATCCCGTTTCCAACGAAGGCCTCAAAGAGGTCCAAATATCCACTTGCAGATTC

>C3F83

CACAAAAAGTGTTTCCAATCTGCTCCGCCTAAAGGAAGCTTCAACTCTGTGAGTTGAATACCCACAACCCAAAGAAGTTACTGAGAATTCTTCTGTCTAGCATTATATGAAGAAATCCCGTTTCCAACGAAGGCCTCAAATACATCCAAATATCCAGTTGCTGACTT

>C3F86

TACAAAAAGAGTGTTAGAAAACTGCTCTTTCCAAAGTAAGGTTCAACTCTGTGAGTTGAATGCACACATAACAATCAAGAAGTTTCTGAGAATTCTTCTGTCCTGGTTTATATGAAAAATCCCGTTTCCAACGAAGGCCTCAAAGACGTTTAAATATCCACTTGCAGACT

>C3F88

TACAAAAAGAGTATTCAAAACTCTTCTATCGAAAGGAAGTTTCAACTCCATGAGTTAAATGCACATATCACAAATAATTTTCTGAGGATTCTTCTTTcAAGTTTTATATGAAGAAATCCCGTTTCCAAAGATGGCCTCAGAAAAGTCCCAATATACACTTGCAGATTC

>C3F95

TACAAACTGAGTGTTTCCAAACTGCTCTATGAAAAGAAAGTTAAACTCTGTGAGTTGAATGCACACATCACAAAGTAGTTTCTGAGAATGATTCTGTCTAGTTTTTATACGAAGATATTTCCTTTTCTACCATTGGCCTCAAAGCGCTTGAAATCTCCACTTGCAAATTC

>C3F96

TACAAACTAAGTCTTTCCAAACTGCTCTATGCAAAGAAATGTTCAACTCTGTGAGTTTAATGCACACATCACAAAGCAGTTTCTGAGAATGATTCCTCTAGTTTTTATACGAAGATAGCCTTTTCTACCATTGGCCTCAAGGCTCTTGAAATCTCCACCTGAAAATTC

>C3F98

TACAAACTGAGTGTTTCCAAACTGCTCTATGAAAAGAAAGGTTAAACTCTGTGAGTTGAACGCACACATCACAAAGTAGTTTCTGAGAATGATTCTGTCTAGTTTTTATACGAAGATATTTCCTTTTCTACCATTGGCCTCAAAGCGCTTGAAATCTCCACTTGCAAATTC

>C3F9

TACAAAAAGAGTGTTTCAAACTGCTCTTCAAAGGAAGGTTCAACTCTGTGAGTTGAATCACACATCACAAAGAAGTTTCTGAGAATTCTTCTGTCTAGTTATATGAAGAAATCCCGTTTCCAACGAAGGCCTCAAAGAGGTCCAAATATCCACTTGCAGATT

>C4F116

GAATCTGCAAGTGGATATTTGGATAGCTTTGAGGATTTCGTTGGAAACGGGATTACATATAAAATCTAGAGAGAAGCATTCTCAGGAACTTCTTTGTGATGTTTGCATTCAAGTCACAGAATTGAACATTCCCTTTATAGAGCAGGTTTGAAACACTCTTTCTGTA

>C4F157

GAATCTGCAGGTGGATATTTGGATAGCTTTGAAGATTTCGTTGGAAACGGGAATTTCTTCATATAAAATCAAGACAGAAGCATTCTCAGAAACATCTCTGTGATGTTTGCATTCAACTCATAGAGTTGAACACTTCCTTTCATAGAGCAGGTTTGAAACACTCTTTCTGCA

>C4F187

GAATCTGCAGGTGGATATTTGGATAGCTTAGAGGGATTCGTTGGAAAGGGGATATCTTCATATAAAATGTAGACAGAAGCATTCTGAGGAACTACTTTGTGATATTTGCATTGAAGTCACAGAATTGAACATTCACTTTGATAGAGCAGGTTGGAAACACTCATCCTGTA

>C4F190

GTGCAAAAACACTGCTTCAGAACTGCTCTCCCAAAGGAAGGTTCAACTCTGTGAGTTGAATGCAGACATCACAAAAAAGTTTCTGAGAATGCTTCCGTCCAGTTTGTATGTGCAGATACCCCGTTTACAACGAATTCCTCAAGAGCTCCAAATATCCTCAAGCAGATT

>C4F194

CTACAAAAGCAGTATTTCAAGACTGCTCTATCAAAAGGAAGGTTGAATTCTGTGAGTTGAATCCACACATCACAAAAAAGTTTCTGAGAATGCTTCTGTCTAATTTGTATGTGCAGATATCCCGTTTACAACGAATTCCTCAAAGAGCTCCAAATATCCTCAAGCAGATT

>C4F198

CTTCAAAAGCAGTATTTCAAGACTGCTCTATCAAAAGGAAGGTTGAACTGTGTGAGTTGAATCCACACATCCCAAAAAAAGTTTCTGAGAATGCTTCTGTCAATTTTGTGCAGATATCCCGTTGGCAACGAATTCCTGAAAGAGCTCCAAATATCCTCAAGCAGATT

>C4F199

CTGCAAAAGCAGTGCTGCAAAACTGCTCTATCgAAAGAAAGGTTGAACTCTGTGAGTTGAGTTCACACATCACAAAAATGTCTCTGACCATGTTCCTGTCTACCTTGTATGTGCAGATATCCCGTTTACAACGAATTCCTCAAAGACCTACAAATATCCTCAAGCTGATT

>C4F19

CTACCTGGAAGGGACATTTCGAGCGCTTTGAGGCCTATGGTGAAAAAGGAAATATCTTCTCATAAAAACCAGAAAGAAGCATTCTCAGAAACTTCTTTGTGTTGTGTGTACTCAAGTAACAGTGTTGAACCTTCCTTTTGACACAGCAGTTTTGAAACACTCTTTTTGTAG

>C4F201

CTAGAAAAGGATGGTTTCAAAACTGATCTATCCAAAAAGACGTTCAACTCTGTGACTTGAATGCACACAACACAAAATTTTCTGAGAATGCTTCTGTCTAGATTGTATGTGGAGATATCTCGTTTGCCACGAATTCCTCCAAGAGCTCCAAATATCCTCAAGCAGATT

>C4F205

CTACAAAAAGAATGTTTCAAAACTGCTCTATCAAAGACGGAGTTCAACCCTTTGACTTCAATGAACACAACACAGAGCAGTTTCTGAGAATGCTTCTGTGTAGTTTTTATCCGAAGATATTTCCTTTTCCACCATAGGCCTCAAATCGCTCCAAATATCCACTTGTAGATC

>C4F207

CTgCAAcAAGAATGTTTCacAACTGCTCTATCAAAGAAAGCGTTCAcCTCTGTGAGTTGAATGCACACATCCCAAAGCAGTTTCTGAGAATGCTTCTGTGTAGCTTCCATGTGAAGGTATCTCCTTTTCCACCATAGGCCTCATATCGCTCCAAATACCACTTGGTaTa

>C4F211

ATGCAGAAAGAATGTTTCcAATCTTCTCTATCAAAGGAAAGaTTCAACTCTGTGAGTTGAATGGACATATCACAAAGAACTGTCTGAGAATACTTCTGTCTAGcTTTTATGTGAAGATATTTCCTTTTCCACCgTAGGCTTCAAAGCGCTCCAAATGAACACTTGCAGATT

>C4F214

CTGCAAAAGCAGGATTTCAAGACAGCTCTATCAAAAGGAAAGTTGAACTCTGTGAGTTGAGTTCACACATCCCTAAATGTTTCTGAGAATGCTTCTGTCTAGTTTATATGTGAAGATATTTCACTTTCCACCATGAGCCTCTAAGCGCTCCAAATGAACACTTGCAGAGT

>C4F217

GTAAGAAAACACTGTTTCAAAACTGCTTTCTCAAAAGGAAGGTTCAACTCTTTGAGTTCAATGCGCACATCACAAGGAACTGTCTGAGCAAACTTCCGTCTAGTTTTTATGTGACGATATTACCATTTCCACCGCAGGTTTCGAAGCGCTCCAAAAGAACACTTGCAGATA

>C4F41

TATCTGGAAGTGGACATTTCAAGCGCTTTCAGGCCTATGGTGAGAAAGGAAATATCTTCAATAAAAACTAGACAGAAGCATTCTCAGAAACTTATTTGTGATGTGTGTCCTCAACTAACAGAGTTGAACCTTTGTTTTGATACAGCATTTTGGAAACACTCTTTTTGTA

>C4F86

AATCTGCAAGTGGATATTTGGATAGCTTTGAGGATTTCGTTGGAAACGGGATATCTTCATATAAAATCTAGACAGAAGCATTCTCAGAAACTTCTTTGTGCTGTATGTCCTCAATTAACAGAGTTGAACCTTTGTTTGGATACAGCATTTTGGAAACATTCCTTTAGTAG

>C5F106

GAATTTCCAATTGGAGATTTCAAGCGCTTCGGGGCCAATGGTAGAAAAGGAAAAATCTTCACATAAAAACTAGACAAAATCATTCCCAGAAACTGTGTAGTGATGTGTATGTTTAACTCACAGAGTTTATCCTTTCTTTTCATAGAGCAGTTGGGAAACACTCTGTTTGAA

>C5F107

CGATAAAAGAGTTTTACAAAACTGCTCTATCAAAAGAAAGGTTCAACGCTGTGAGTTGAATCCACATATCACGAAAAAGTTTCTGAGAATGCCTCTATCTACTTTTTATGTGAAGATATTCCGGTTTCCAACGAAGGCCTCAAAGCGCTCCAAATATCTACTTGCAGATT

>C5F116

CTACCAAAAGAGTGTTTCAAAACTGCTCTGTGAAAAGAAACGTTCAACTGTGTTAGTTGAATGCCCACATCACAAAGAAGATTCTGAGAATATTTCTGTCTAGTTTTTATTAGAAGATATTCCCGTTTCCACCAAAGGACACAAAGCGAAGCCAACTATCCGCTTGCAGAT

>C5F123

CTTACAAAAACACGTTTCAAAACTGCTCTATCAAAGGAAAGGTTCATCTCTCTGGGTTCAACGCACACATCACAAAGAAGTTTCTGAGAATGCTTCTGGCTAGTTTGTGTGTGAAGATATTCCCATTTCCAACAAAGGCTTCAAAGCCCTCCAAATATTCACCTGCAATTG

>C5F12

CTACAAAAAGAGTGTTTCAAAACTGCTCTATAAAAGGAAGGTTCAACTCTGTGAGTTGAATGCACACATCACAAAGAAGTTTCTGAGAATGCTTCTGTCTAGTTTTTATGTGAAGATATTTCCTTTTCCACCATAGGCCTCAAAGCGCTCCAAATATCCACTTGCAGATT

>C5F130

AAGTCTGCAAGTGGATATTCGACCTCTTTGAGGCCTTCGTTGGAAACGGGTTTTTTTCATATAAGGCTAGACAGAAGAATTCTCAGTAACTTCCTTGTGTTGTGTGTATTCAACTCACAGAGTTGAACGTTCCTTTACACAGAGCAGATTTGAAACACTCTTTTTGTG

>C5F137

AAGTCTGCAAGTGGATATTCGACCTCTTGAGGCCTTCGTTGGAAACGGGTTTTTTTCATATAAGGCTAGACAGAAGAATTCTCAGTAACTTCCTTGTGTTGTGTGTATTCAACTCACAGAGTTGAACGATCCTTTACACAGAGCAGATTTGAAACACTCTTTTTGTG

>C5F163

AAGTCTGCATGTGGATATTTGGACCGCCATGAGGCGTTCTTTGGAAATGGTATTTCTTCATTTAAGGCTACACAGAAGAATTCTCAGTAACTTCCTTGTGTTGTGTGTATTCAGCTCACAGAGTTGAACCTTCTTTTAGATAGAGCAGATTTGAAAGACACTTTTTGGG

>C5F164

CACAGAAACAGTGTTTCAAAACTGCTCTAACAAAAGAAAGATTCAACTCCGTGATTTGAATGCACACATCACAAAGCATTTTCTGTGAATCCTTCTGTCTAGTTTTTATATGAGGATATTTCCTTTTCTACCACGGGCATCCAAGCGTTCCAATTCTCCAATTGTAGATTG

>C5F171

CTACAGAGTGTTTCAAAACTGCTCTATCCAAAAAAAGTTTCAACTCGGTGAGTCGAATGCACATATCACAAAGCAGTTTCTGAGAATGCTTTCGTCTATTTTTCCCAGGAAGATATTTCCTTTTGGACCGTAGGCCTCAAATCGCTCCAGATATCCACATGCAGATT

>C5F186

TACAAACTAGACAGAAGCATTCTCAGAAACTGCTTTGTGATGTGTGCATTCAACTCACAGAGTTGAACCTTCCTTTTGAGAGAGCAGTTTTGAAACAGTCTTTTTGTAGTATCTGCAAGTGGATATTTGGAGCGATTTGAGGCCTATGATGGAAAAGGAAATATCTTCACA

>C5F198

TAAAAAACTAGACGGAAGCATTTTCAGAAACTGCCTTGTGATGTGTGCATTCAACTCACAGAGTTGAACCTTCCTTTTGAGAGAGAAGTTTTGAAACAGTCTTTTTGTAGTATTTGCAAGTGGATATTTGGAGCGATTTGTGGAGTATGGTGGAAAATGAAATATCTTCACA

>C5F200

TAAAAAATAGACAGAAGCATTCTCAGAAACTGCTTTGTAAATGTGCATTCAACTCACAGAGTTGAACCTTCCTTTTGAGAGAGCGGTTTTGAAACAGTCTTTTTGTAGTATCTGCAAGTGGATATTTGAGGATTTGAGGCAAGAAGGAAAAGGAATACCTTCAAA

>C5F207

TACAAACTAAACAGAAGCATTCTCAGAAACTTCTTGTGATGTGTGCATTCACCTAACAGAGTGGAACCGTTCTTTTGATAGAGCAGTTTTGAATCAGTCTTTTGGTAGGACCTGCAAGTTTTCATTTGGAGCGCTTTGAAGCCCATGGTGGAAAAGGGACTATCTTCACA

>C5F211

TAaAAACTAGACAGAAGCtTTCTCAGAAACTGCTTTGTGATGTGTGCATTTAACTCAcAGtCTTGATCCTTCTTTaGTAGAGCAGTGTTGAAACACACTTTTTGTAGAACCTGCTAGTGTTCATTTGGAGAGATTTGTTGCCTATGGTGGAAAAAGGATTATCTTCTCT

>C5F216

TAAAAAGTAGACCCAAGCATTCTCAGAAGGTTCTTTGTGATGTGTGCGTTCAACTCACAGACTTGAAACTTTCTTTTGATAGAGCAGTGTTGAAACACACTTTTTGTAGAATCCACAAGTATTCTTTGGAGCGCTTTGTTGCCTATGTGGGAAAAAGGAATATCTTCACT

>C5F220

TAAAAACTAGACAGAAGCATTCTCAGAAACTCCTTGTGAGTGTGTGTTCAATTCACATTGTTGAACCTTTCTTTTGATAAGCAGTGTTGAAACAACATTTTGTAGAATCTGCAAGTGTTCATTTCAGTGCTTTGCCTATGTTGGAAAAAGTGATATCTTCACC

>C5F232

TAAAAACTAGACAGAAGCATTCTTAGAAACTGCTTTGTGATGTGTGTGTTCAATTCACAGAGTTGAAACTTTCCTTTGACAGAGCAGGTTTGAAACACTGCTTCTGTAGAATCTGCTTGTGGATATTGGGAGCTCCTTGAGGAATACGTTGTAAAAGGCATATCTTCACA

>C5F239

AAAAACTAGGCAGAAGCCTTCTCAGGAACTTCATTGAGATGTGTGCATTCAACTAACAGAGTTGAAACTGTCTTTTGACAGAGGAGGAATGAAACACTCCTTTTGTAGTATCTGATTGTGTATATTTGGAACTCTTTGAGTTATTCGTTGGAAACGGGTATCTTCACA

>C5F66

CTACAAAAATAGTGCTTCAAAACTACTCTATGGAAAGGTATGTTCAACACTGTGAGATGAATGCAAACGTCACAAAGAAGTTGCTGAGAATGCTTCAGTCTAGTTTCTATGGGAAGACATTTCCTTTTGCACCACAGCCCTCAAAGCACTCCAAATGTCTACTTGCAGATT

>C5F73

GAATTTGCAAGTGGAGATTTCAAGCGCTTTGAGGTCAAAGGTAGAAAAGGAAATATCTTCGTATAAAAACTAGACAGAATCATTCTCAGAAACTGCTTTGTGATGTGTGCGTTCAACTCACAGAGTTTAACCTTTCTTTTCATAGAGCAGTTAGGAAACACTCTGTTTGTA

>C5F80

GAATTTGCAAGTGGAGATTTCAAGCGCTTTGAGGCCAAAGGCAGAAAAGGAAATATCTTCGTATAAAAACTAGACAGAATCATTCTCAGAAACTGCTCTGCGATGTGTGCGTTCAACTCTCAGAGTTTAACTTTTCTTTTCATTCAGCAGTTTGGAAACACTCTGTTTGTA

>C5F87

CAATTGGCAAGTGGAGATTTCAAGCGCTTTAAGGTCAATGGCAGAAAAGGAAATATCTTCGTTTCAAAACTAGACAGAATCATTCCCACAAACTGCGTTGTGATGTGTTCGTTCATCTCACAGAGTTTAACCTTTCTTTTCATAGAGCAGTTAGGAAACAGTCTGTTTGTA

>C5F92

GAATTTGCAAGTGGAGATTTCAGCCGCTTTGAAGTCAAGGTAGAAAAGGAAATATCTTCCTATAAAAACTAGACAGAATGATTCTCAGAAACTCCTTTGTGATGTGTGCGTTCAACTCACAGAGTTTAACCTTTCTTTTCATAGAGCAGTTAGGAAACACTCTGTTTGTA

>C6F13

GTATTTGCATGTGTATATTTAGAGCGCATTGAAGCCCACAGTAGAAAAGGAAATAACTTCACCTAAAACCTAGACAGAAGCAATCTCAGAAACTACTTTGTGATGTGTACATTCAACTCACAGAGTGGAACTTTCCTCTTTATAGAGCAGTGTTGAAACACTCTTTTTGTAG

>C6F16

AATCTGCAAGTGGATATTTGGACCTCTTTGAGGCCTTCGTTGGAAACGGGATTTCTTCTATAAACTAGACAGAAGAATTCTCAGAAACTTCTTTGTGATGTGTGCATTCAACTCACAGAGTGGAACCTTCCTTTTGATAGAGCAGTTTTGAAACCTGTTTTTGTAG

>C6F32

TATTTCCAAGCGGATATTTAGAGCGCCTTGAAGCCTATGGTAGAAAAGGAAATATCTTCCCATAAAACCTAGACAGAAGCAATCTCAGAAACTACTTGTGATGTCTGCATTCAACTCACAGAGTGGAACATTTCTCTTGATAGAGCAGTTTTGAAACACTCTTTCTGTAG

>C6F44

AATTTGCAAGGGTACATTGAGAGCGCTTTCAGGCCTATGGTAGAAAAGGGAATATCTTTCCATAAAAGGTAGACAGAAGCAATCTCAGAAACTACTTTGTGATGTGTGCATTCAACTCACCGAGTGCAACGTTCCTCTTGACCGAGCAGTTTGGAAACATTGTTTCTGTAG

>C6F46

AATCTGCAAGTGGATAATTGGACCTCCTAGAGGCCTTCGTTGGAAACGGGATTTCTTCATCTAAACCTACAGAGAAGAATTCTCAGTAACTTCTTCGGATGTGTGCATTCGACGCACAGAATGGAACATTCCCTTTGATAGAGCAGTTTTGAGACACCGTTTTTGTAG

>C6F48

GATTTCCAAGTGGATATTTAGACCACTTTGAAGCCTATGATAGAAAAGGAAACATCTTCATGGAAAACATAGATAGAATCATCCTCAGAAACAACTTTGTGATGTGTGCGTTGAACTCACCGTCTTTAACCTTTCTTTTGGTAGAGAAGTTTTGAAACACTCTCTTTGTA

>C6F49

AATTCCCAAGTGGATATTTAGAGCACTTTGAAGTCTCTGCTAGAAAAGGAAACATCTTCATGTAAAAAGTAGATAGAATCGTTCTCAGAAAGTGCTTAGTGACGTGTGCGTTCAACTCACAGAGTTTAACGTTTCTTTTGATAGAGCGTTTCTGAAACACCCTTCTTGTA

>C6F50

GTAGCTGCAAGTGGATATTTGGACCTATTGGAGGCCTTCTTTGGAAATGGGATTTCTTCATGTAACTCTAGATTGAAGAATTCTCAGAAACTCCTTTGTGATGTGTGCATTCAATTCAAAGAGTGAAACCTCCCTTTTCACAGAGCAGTTTTGAAACACTGTTTTTGTAG

>C6F51

AATCAGCTTGTTTGTATTTGGACCTCCTTGAGGCCTTCGTTGGAAACGGGTTTTCATCTTATAAACCCAGACAGAAGAATTCTCAGAGTCTTCTTTGTGATGTGTGCTTTCAACTCACCGAGATAAAGATTTCTCTTGATAGAGCAATTTGGAAACACTCTTTTTGTAG

>C6F54

AAGTCTACAAGTGGATATTTTGAGCCCTTGGAGGCATTCTTTGGAAAAGGGAATGTCTTCACATAAAAGGCAGACGGAAGTGTTCTCAGAAACTGCTTTGTGATGTCTGTGTTCAACTCACAGAGTTTAACATTTCCTTTGATAGAGCGGTTTAGTAACCCTCTCTTTGTAG

>C6F57

AATCTGCAAGTGGATAATTGGACCGCCTTGAGGCCTTCGTTGGAAACGGGATTTCTTCATGTTACTCTAGATAGAAGAATTCTCAAACACTGCTATGTGATGTTTGCATTCAAGTCACAGAGTGCAACATTCCTCTTGGTAGAGCAGTTGGGAAACACTCCTTTTGTAG

>C6F9

GATTTCCAAGGGGATATTTATAGCGCATTGAGCCTACGGCGGAAAAAGAAACATCTTCCTATAAAAACTAGACAGAATACTTCTCAGAATCTGCTTTGCGATGTGTGCGTTCAACTCACAGAGTAAAACTTTTCTTTTGATAGAGCAGTTTTGAAACACTCTTTTTGTA

>C7F16

CTACAAGAAGATTGTTTCCAGGCGGCTCTATCAAAAGAAATGTTCAACTATGGGAGTAGAATACACACATCACAAAGTCGTTTCTGAGAATGCTTCTGTCTAGTTTTTATGTGAAGATATTTCCTTTTCTACCAGAGGCCTGAAAGCGCTCCAAATATCCAATTGCAGATT

>C7F18

TTACAAACAGAGTGTTTCCAGACTGCTCTATCAAAAGAAATTTTCAACTATGGGAGTAGAATGCACACATCACAAATTCGTTTCTGAGAATGCTTCTGTCTAGTTTTTATGTGAAGATATTTCCTTTTCTACCATAGGCATAAATGTGCTCCAAATATCCACTTGCAGATT

>C7F23

CTACAGAAAGAGTGTTTCAAAACTGCTCTATCAAAAGGAAGGTTCAACTCTGTGAGTTGAATGCAGACATCATAAAGGAGTTTCTGAGAATGCTTCTGTCTTGTTTTAATGTGAAGATATTTCCTTTTCAACCGATAGCCTAAAAGAGCTCCAAATGTCCAACTGCAGATT

>C7F24

CTACAAAAGGAGTGTTTCAAAACTGCTCTAACAAAAGAAAGGTTCAACCTGGGAGTTGAATGCACACATCACAAAGAAGTTTCTGAGAATGCTTCTGTCTAGTTTTAATGCGAAGATATTCCCTTTTCCACCATAACCTTCAAAGCGCTCCAAATGTCCATTTGCAGATT

>C7F26

CTAAAAAAAGAGTGTTTCAAAACTGCTCTATCAAAAGGAAGGTTCAATTCTGTGAGTTGAATGCACACATCACAAAGTAGTTTCTGAGAATGCTTCTGCCTAATTTTTAATTTTAAGATATTCCCATTTCCAAAGAAGGCTTCAATTTGCTCCAAATATCCACTTGCAGATT

>C7F29

ATAAAGGAAGAGTTTTTCAAAACTGCTCAATCCAAAGGAAGGTTCAGCTCCGTGAGTTCAATGCCCACATCACAAAGAAGTTTCTGAGAATGCTTCTGTCTAGTTTCAAGGGGAAGATATTCCCGTTTCCAACGAAGGCTTCAAAGCGCTCCAAATATCCACTTGCAGAGT

>C7F32

CTTCAAAAAGAGGGTTTCAAAACTGCTCTAGCAAAAGGAAGTTTCAACGCTGTGAGTTGAATGCACACATCACAAAGAAGTTTCTGAGAATGCTTCTGTCTAATTTTTATGTGAAGATAATCCTGTTTCCCAAGAAGGCTTCAAAGCACCCTAATATCCGCCTCCAGATT

>C7F33

GTGCAAAAAGGGTGTTTCAAAACTGCTCTATCAAAAAGAAGCTTCAACTCTGTGAGTTGAATGCACAAATCACAAAGAAGTTTCTGGGAATACTTCTGTCTATTTTTATGTGACGATACTCCCGTTTCCAAAGAAGGCTTCAAAGCACTCCAAATATCCACCTGCAGATT

>C7F34

GTACAAAAAGAGGGTTTCAAAACTGCTCTATCAAAAGGAAGGTTCAACTCTGTGAGATGAATGCACACATCACAAAGTGGTTTCAGAGAATGCTTCTTTCTAGTTTTTAGGTGACGATATTCCCGTTTCCAATGTAGCACTCAAAGAGCTCCAAATATCCTCCTGCAGATT

>C7F36

CTTCGGAAAGACTGTTTCAAAACTGCTCTATCAAAAGAAAGTTTCAACTTTGTGTGTAGAATGCACACATCACAAAGTCGTTTCTGAGAATGCTTCTGTCTAGTTTTTATATGAAGATATTTCCTTTTCTACCTAAGCCTCAAAGCACTGCAAATATGAACTGGCAGATT

>C7F40

AAGTCTGCAAGTGGATATATGGACCTCTTTGAGGCCTTCGTTGGAAACGGGATTTCTTCATTTAATGCTAGACAGAAGAATTCTCAGTAAATTCTTTGTGTTGTGTGCATTCAACTCACAGAGTGGAACGTCCCTTTAGACAGAGCAGATTTGAAACACTCTTTTTGTG

>C7F51

ATGTCTGCAAGTGGATATTTGGACCTCTTTGAGGCCTTCGTTGCAAACGGGGTTTCTTCCTTTCATGCTAGACTAAGAAGAGTTCTCAGTAACTTTTTTGTGTTGTGTGTATTCAACTCACAGAGTTGAACCTTGCTTTAGAGAGAGCAGATTTGAAACACTCTTGCTGTG

>C7F58

GAATTTGCAAGTGGAGATTTCAAGCGATTTGATGCCAACAGTAGAAAAGGAAATATCTTCAAATAAAAACCAGACAGAATCATTCTCAGAAATCTTTGTGATGTGTGCGTTCAACTCACAGAGTTTAACCTTTCTTTTCATAGAGCAGTTTGGAAACACTCTGTTTGTA

>C7F75

TTTTAAAAAGAATGTTTCAAAACTGCACTATCAAAAGAAAGGTTCAGCTCTGTGAGTTGAATGCACACATCTCAAAGAAGTTTCTGAGAATGGTTCTGTCTAGTTTTTATGTGAAGGTATTCGCGATTCCAATGAAGTCTTCAAAGCGCTCCAAATATCTAAATGCGGATTC

>C7F77

TACAAAAAGAGTGATTCAAAACTGGTCTATGAAAAGGAAGGTTCAGCTCTGTGAGTTGAACGCACACATCACAAAAAGTTTCGGACAATGCTTCCATTTAGTTTTTAGGTGAAGATATTACCTTTTCAACCACAGCCTTCAAAACGCTCCAAATGTCCACTTGCAGATT

>C7F82

CTACAAAAAGACTGTTTCAAAACTGCTCTATAAAAGTAAGGTTGTACTTTGTTAGTTGAATGCACCCATCAAAATGAAGTTTCTGAGAACACTTCTGTCTACTTTTCATGTGAAGATATTTCCTTATCCACAATAGTCCCCAAAGCCCTCAAAATGCCCACTGAAGATT

>C7F85

CTACAAAAAGAGTGTTTCAAAACTTCTCTATCAAAAGTAAGGGTCTGCTTTGTGAGTTGAATGTACACATCAAAATGAAGTTTCTGATTATACTTCTGTCTACTTCTTATGGGAAGATATTTCCTTATCCGCAATGGTCCTCAAAGCCCTCGAAATGCCCACTTGAAGATT

>C7F89

CTTCAAAAAGAGTGTTTCAAAACTGCTCTATCAAAACTGTTCTATCAACTCTGTGAGTTGAATGGACACATCACAAAGACGTTTCTGAGAATGCTTCTGTCTAGGTTTTAGGTGAAGATATTCTCGTTTCCAAAGAATGCTTCAAAGAGTACTTAAATATCCGCCTGCAGATT

>C8F12

GAATCTGCAAGTGGATATTTGGATAGCTTTGAGGATTTCGTTGGAAACGGGATTCATATAAAATAGACAGCAGCATTCTCAGAAACTTCTTTGTGATGTTTGCATTCAAGTCACAGAGTTGAACATTCCCTTTCATAGAGCAGGTTTGAAACACTCTTTTTGTA

>C8F15

GAATCTGCAAGTGGATATTTGGATAGCTTTGAGGATTTCGTTGGAAACGGGAATGTCTTCAAAGAAAATCTAGACAGAAGCATTCTCAGAAACACCTTCGTGATGTTTGCAATCAAGTCACAGAGTTGAACCTTCCGTTTCATAGAGCAGGTTGGAAACACTCTTTTGTA

>C8F23

GAATCTGCAAGTGGATATTTGGATAGCTTTGAGGATTTCGTTGGAAACGGGATTAATATAAAAAGTAGACAGCAGCATTCTCAGAAACTTCTTTGTGATGTTTGCATTCAAGTCACAGAGTTGAACATTCCCTTTCATAGAGCAGGTTTGAAACACTCTTTTTGTA

>C8F33

AATCTGCAAGTGGATATTTGGATAGCTGTGAGGATTTCGTTGGAAACGGGAATGTCTTCATAGAAAATTTAGACAGAAGCATTCTCAGAACCTTGATTGTGATGTGTGTTCTCCACTAACAGAGTTGAACCTTTCTTTTGACAGAACTGTTCTGAAACATTCTTTTTATAG

>C8F35

GTGTCTGGAAGCGGGCATTTGGAGCGCTTTCAGGCCTATGCTGAAAAAGGAAATATCTACCTACAGAAACTAGACAGAAGCATTCTGAGAATCACGTTTGTGATGTGGGTACTCAACTAACAGTGTTGATCCATTCTTTTGATACAGCAGTTTTGAACCACACTTTTTGTA

>C8F43

GTATCTGGAAGTGGACATTTGGAGCGCTTTCTGAACTATGGTGAAAAAGGAAATATCTTCCAATGAAAACAAGACAGAAGCATTCTGAGAAACTTATTTGTGATGTGTGTCCTCAACAAACGGACTTGAACCTTTCGTTTCATGCAGTACTTCTGGAACACTCTTTTTGAA

>C8F45

GTATCTGGAAGTGGACATTTGGAGGGCTTTGTAGCCTATCTGGAAAAAGGAAATATCTTCCCATGAATGCGAGATAGAAGTAATCTCAGAAACATGTTTATGCTGTATCTACTCAACTAACTGTGCTGAACATTTCTATTGATAGAGCAGTTTTGAGACACTCTTCTTTTG

>C8F8

GAATCTGCAAGTGGATATTTGGAGCTTTGAGGATTTCGTTGGAAACGGAATATCTCAATAAAAACTAGACAGAAGCATTCTCAGAAACTCTTTGTGATGTTGCATTCAACTCACAGAGTTGAACCTTCTTTCATAGAGCAGTTTGAAACACTCTTTTTGTA

>C9F33

GAATCTGCAAGTGGATATTTGGATAGCTTTGAAGATTTCGTTGGAAACGGGAATATCTTCATATAAAATCTAGACAGAAGCATTCTCAGAAACTTCTTTGTGATGTTTGCATTCAAGTCACAGAGTTGAACATTCCCTTTCATAGAGCAGGTTTGAAACACTCTTTCTGTA

>C9F58

AATCTGCAAGTGGATATTTTGATACCTTTGAGGATTTCGTTGGACACGGGATATCTTCATATAAAATCTAGACAGAAGCATTCTCAGAAACTTCTTTGTGCTGTATGTCCTCAATTAACAGAGTTGAACCTTTGTGTGGATACAGCATTTTGGAAACATTCCTTTAGTAG

>CXF14

CTACAAAAAGAGTGATTCCAATCTGCTCTATCAATAGGATTGTTCAACTCCATGAGTTGAATGCCATCCTCACAAAGTCGTTTCTGAGAATGCTTCTATCTAGTTTTTATGTGAAGATATTTCCTTTTCCACCACAGGCCTCAAAGCCCTCCAAACGTCCACTTGCAGATT

>CXF21

GTAGAAAAAGTGTGTCAAAGCTGCGCTATCAAAGGGAAAGTTCAACTCTGTGAGGTGAATGCAAACATCCCAAAGAAGTTTCTGAGAATGCTTCCGTTTAGCTTTTAGGTGAAGATTATCCCGTTTCCAACGAAACCTTCAAAGAGGTCCAAATATCCCCTTGCGGATC

>CXF26

CTCGAAAAAGAGTGTTTCATAGCTGCTCTTTCAAAAGGAAAGTTCAACTCTGGGAGTTGAATACAAACATCACAAAGTAGTTTCCGAGAATGCTTCTGTTTAGTTTTTATGTGAAGATGATCCCGTTTCCAGTGAAATCTTCAAAGAGGTCCACATATCCCCTTGCAGATT

>CXF31

CTACTACAAGGGTGTTGCAAACCTGAACTATCAAAGGAAGGTTCAACTCTGTGAGTTGAATACAAACATCACAAAGAATGTTCTGAGTTTGCTTCCGTTCAGTTATGGGAAGTTGATCCCGTTTCCAACGAAATCCTCAGAGAGGTCCAAATATCCCCTTGCAGATTC

>CXF36

CTCGAAAAAGAGTGTTTCATAGCTGCTCTTTCAAAAGGAAAGTTCAACTCTGGGAGTTGAATACAAACATCACAAAGAATGTTCTGAGTTTGCTTCCGTTCAGTTATGGGAAGTTGATCCCGTTTCCAACGAAATCCTCAGAGAGGTCCAAATATCCCCTTGCAGATTC

>CXF38

CTGCCAAAAGAATATTTCAAAACTGCTCTATGAAAAGCAATGTTAAACTCTGTGGCTCGAACACAAACATCACAAAGCAGTTTCTGAGAATGCTTCAGTTTAGTTTTTCTGTGGAAATATTCCCGTTTCCAAAGAAATCTTCAAAGAGGTCCACGTATCCACTTACAGATT

>CXF41

CCACAGAAAGAGTGTTTCGAAACTGCTGTTTCAAAAGGAATCTTCAACTCTGTGAGTTGAATGCAATCATCACAAAGAAGTTTCTGACAATGCTTCTCTCTCGTCTTTCTGTGAAGATAAAGGAAAAGGCTTTCAGGCCTTTTCCACCACAGGCCTGAAAGCGCTCCAAATGTCCACTTGCAGATT

>CXF45

CTACAAAAAGAGTGTTTGCAAACTGCTCTATCAAAAGGAATGTTCAACTCTGGGAGTTGAATGCAATCATCACAGAGCAGTTTCTGAGAATGCTTCTATGTCGTTTTTAGGAGAAGATATTTCCTTTTCCAACACAGTCCTCCAAGCCCGCTAAATAGCCACTTGCACATT

>CXF50

CTACAAAAAGACAGTTTCAAAACTGCTCCATCAAAAGGAGGGTTCAACTGTGTGACTTGAATGCAATCATCACTCAGAAGTTTCTGAGAATGCTTCTCTTTAGTTTTTACGTGAACATATACCCGTTTCGAACGAAGGCCACCCAGTGGTCCAAATATCCACTTGCAGATT

>CXF56

CTACAGAAAGAGTGTTTCGAACCTGAACTCTCAAAGGCAGGTTCATCTCTGCGAGTTAAATGCATTCATCATGAAGAACTTTCTCAGAGTGTTTGTGTTTAGTTATGGGAAATTATTCCCGTTTCCAACGAAATCCTCAGAGAGCTCCAAATATCCACCTGCAGATTC

>CXF60

TACCAAAAGTGTATTTGGAAACTGCTCCATCAAAAGGCATGTTCAGCTCTGTGAGTGAAACTCCATCATCACAAAGAATATTCTGAGAATGCTTCCGTTTGCCTTTTATATGAAGTTCCTTCCTATACTACCGTAGGCCTCAAAGCAGTCCAAATCTCCATTTGCAGATT

>CXF63

TACAAAACGTGTGTTTGGAAACTGCTCCATCATAACGAATGTTCAGCTCCCTGAGTTAAACTCCATCGTCACAAAGAATTTTCTGAGAGTGCTACCGTCTGGTTTTTATATGAAGTTCTTTCCTTCACTACCACAGGCCTCAAAGCGGTCCAAATCTCCACTTGCAGATT

>CXF66

CCAAAGAAAGAGGGTTTCAAAACTGCTCCATCAGAAGGATTGTTCAACTCTGTGAGTTGAATGCAGTCATCGCAGAAAACTTTCTGAGAATGCTTCTGTCTAGGTTTGATGTGAAGATATAGACGTTTCAAACGAAGGCTACAAAGTGGTCAAAATATACACTTGCAGATT

>CYF15

TACAAAAAGAGAGTTTCAAAAcTGCTCTATCAAAAGATAGGTTCAAcTcTGTGATATGAATGCACACATCACAAAGaAGTTTCTCAGAATGCTTCTGTGTAGTTTTtATGTGAAGATATTTCCTTTTCCACCATAGGCCTCAAAGCACTCCAAATATCCACTTGCAGATTC

>CYF16

TACAAAAAGAGAGTTTCAAAAGTGCTCTATCAAAAGATAGGTTCAACTATGTGATATGAATGCACACATCACAAAGTAGTTTCTCAGAATGCTTCTGTGTAGTTTTTATGTAAAGATATTTCCTTTTCCACCATAGGCCTCAAAGCACTCCAAATATCCACTTGCAGATTC

>CYF19

TACAGAAAGACACTTTAAAAACTGCTCTATCAAAAGATCAGTTCAAGTCTGTGGTTTGAATGCACACATCACAAAGAATTTTCTCAGAATGCTTCTGTGTAGTTTTCATATGAAGATATTTCCTTTTCCACCATAGGCCTCAAAGCACTCCAAATATCCACTTGCAGATTC

>CYF20

TACAAAAAGAAAGTTTCGAAATGCTCTCTCAAACGATAGTTTCGACTCTGTGGTATGAATACACACATCACAAAGAAGTTTCTCAGAATGCTTCTGTGTAGTTTTTAAATGAAGATATTTCTTTTTCCACCATAGGCCTCAAAGCACTCCAAATATGCACTTCCAGATTC

>CYF21

CACAAAAAGAGTGTTTGCAAACTGCTCAATCAAAAGAAAGATTTAACTCTGTGAGATGAATCCACACATGACAAAGAAGTTTCTCAGAATGCTTCTGTGTAGTTTTTATGTGAAGATATTTCCTTTTCCACAATAAGACCCAAAAGGCTCCAAATATTCACTTGCAGATT

>CYF26

TACAAAAAGAGTGTTTCAAAACTGCACAATCAAAAGATAGTTCAACTCTGTGAGTTGAATGCGCACAACAAAAAGATGTTTCTCAGAATTATTTCTGTGTAGTTTTTATGTGAAGATATTTCCTTTTCCACAATGGGCCTCAAAGTGCTCCAAATATCCACTTGCAGATTC

>CYF27

TACAAAAAGAGATTTTCAAAACTAtTCAATCAAAAGAAAGGTTCAACTCTGTCAGTTGAATGCACATATCACAAACAAGTTTCTtGGAATGCtTCTGTGTAGTTTTTATGTGAAGATATTTCCTTtTCCACAACAGGCCTCAAAGTGCTCCgAATATCCACTTGCAGATTT

>CYF31

TACAAAAAGAGTGTTTCAAAACTGCTCAATCAAAAGAAAGGTTCGACTCTGGGAAATTAATGCACACATCACAAAGAAGTTTCTCAGCTTCTGTGTAGTTTTCATGTGAAGTTATTTCCTTTTCCACAATAGGCCGCAAAGGGCTCCAAATATCAACTTACAGATTC

>CYF32

tACTAAAgatGTGTTTCCAAACTGCTCAATCAAGAGGAAGTTTCAAGTCTGTGAGcTGAAcGCACACATcACAAAGTAGTTtCTGAGAATGCTTCTGTGTAGTTTTTATGTGAAGATgTTTCCTTTTCCACCAtAGGCtgCAAAGgGCTcCAAATATCCACTTGCAGATTC

>CYF35

TACAAGAAGATTGTTTCAAAACTGCACAAAAAAAGAAATGTTCAATTCTGTTTGATGAATGCACACATCACAAAGAAGTTTCTCAGAATGCTTCTCTGTAGTTTTTATGTGAAGATATTTCCTTTTCCACAATAGGCCTCAAAGGGCTCCAAATATCCACTTCCAGATT

>CYF36

CTAAAAAAAACAGTGTTTCAAAACTGCTCAATCAAAAGATAGTTCAACTCTGTGAGAAGAATGCTCACATCACTGAGAAGTTTCTCAGAATGCTTCTGTGTAGTTTTTATATGAAGATATTTCCTTTCCCACCGTAGGCCACAAAAGGCTCCAAATATCCACTTGCAGA

>CYF39

TACAAAAAGAGAGTTTCAAAACTGCTGTATCAAAAGATAGGGTCAACTCTGCGAGTTGAATAAACACATCACAAATAAGTTTCTGGGAACGCTTCTGTATAGTTTTATGTGAATATATTTCCTTTTCCACCATATGCCTCAAAGCACTCCAAATATCCACTTGCACATT

>CYF40

CTACAAAAGGAGTATTTCAAAACTGCTCAATCAAAAGAAAGGTTCAACTCTGTGAGATGAATGGACACATCACAAAGAAGTTTCTCAGAATGCTTCTGTGTAGTATTTTTGTGAAGATATTTCTTTTCCACCATAGACCGCCAGGGGACACAAATATCCACTTTCAGATTC

>CYF41

TAGGAAAAGAGAGTTTCAAAACTGCTCTACGAAAAGATAGGTTGAACTCTGTGAGATGAATGCACACATCACAAAGAAGTTTCTCAGAATGCATCTGTGTAGTTTTTACGGGAAGATATTTCCTTTTCCACCATCTTCCACAAAGGTCTCCAAGTAACCACTTGCAGATTC

>CYF42

ATAGAAACATAGTCTTTCAAAACTTGTCAATCAAAGAAAGGTTCAACTCCGTGAGATGAGTGCACACATCACAGAGAAGTTTCTCGGAATGTTTCTGTGTAGTTTTTATGTGAAGATATTGCCTTTTCCACAATAGGCCTCAAAGCGTTCCAAATATCCAATTGCAGATT

>CYF43

TACAAAAAGAGAGTTTCAAAACTACTCAAACAAAAGGTTCAATTCTGTGAGTTGAAAGCAAACATCACAAAGAAGTTTCTCAGAATGCGTCTGTGTAGTTTTGATGTGAAGATATTTCCTTTTCACAGTAGAATGCAAAGGGCTCCAAATATCCACTTGGAGATTC

>CYF44

CCACAAAAAAAGTTTTTTAAAACTGCTCAATCAAATGATAGATTAAACTCTGTGAGATTAGTGCACACATGTCAAAAAAGTTTCTCAGAATGCTTCTGTGTACTTTTTAGGGGAAGATATTTCCTTTTCCACCATCGGCCACAAAGGACTCCAAATAACCACATGCAGATT

>CYF45

CTATGAAAAGAATATTTCCAAACTGCTCAATCATAGGAAATGTTCAACTCTGTGAGATGAATGCACACATCACAAGAAATTTCTCAGAATCCTTCAGTGTAGGTTTTATGAGAAGATAATTCCTTTTCCACAATAGTTCTCAAAGCACTCAAAATATCCACTTGCAGATT

>CYF46

TACAACAAGAGAGGTTCAAAACTACTCGATCAAGAGATGGTTTCAACTATGTGAGTTGAATGCACACATCACAAAGAACTATGTCGGAATTCTTCTGTGTAGTTTTTATGTGAAGATATTTCCTTTTCCACAATAGACGTCAAAGTGATCCAGATATCCACTTGCAGATTC

>CYF47

CACAAAAAGAGTGTTTCAAAAGTGCACAACCAAAAGAAAGGTTCAACTAGGTGAGATGAATGCACACATCAGAAGGAAGTTTCTCAGAATGCTTCTGCATAGCTTTTAAGGGAAGATACTTCCTTTTCCAACATAGGCCTCAAAGCACTCCAAATATCCTCCTGGAGATAC

>CYF48

CACAAGAAGAGTGTTTCAAAACTGCTGTATCAAATAAAGTTGAACTCTGTGAGGTGAATGCACACAGCACAAAATGGTTTCTCAGAATGCTTCCTTGTTGTTTTTATATGAAGATGTTTCCTTTTCAACAATAGGCCTCAAAGTGCTTCAAATGTCCACTTGCAGATTC

>CYF49

TACAAAAACCGTGTTTCAAAACTGCCGAATCAAAAGAAAGGTTCAACTCTGTGAGATGAATGCACACATAACAAAGGAGTTTCTCAGAATGCTTCTGTGTAGCTTTTATATGAAGACATTTAGTTTTCCACAACAGGCCTCAAAGCTCTCTCCATATCCACTTGCAGATTC

>CYF50

TACCAAACGAGTATTTCAAAACTGCTCAATCAAATGGAAGGTTCAAAACTGTGACATGAATGCCCACATCACAAAGTAGTTTCTCAGAATGCTTCTGTGTAGTTTTTATGTGAAGATATTTCCTTTTCCACAACAGCGTGCAAAACGCTTCAAATATGCCCTTAGAGATTC

>CYF52

CACAAAAAGAGTGTTTCCAAACTACTCAAATCAAAAAATGATTTCAACTCTGTGAGATGAATGCACACATCACAAACTAGTTTCTCAGAATGTTTCTGCCTGGTTCTCATGCGAAGATAGTTCCTTTTTCACCATAGGCCGCAATGTACTCCAAATATCCACCTGCAGATTC

>CYF53

TACTATGAAAAGAGAGTTTCAAAACTGCTCATTCAAAAGATAGGTTCAACTCTGTGGTTTGAATGCACACAGCACAAAGAAGTTTCACAGAATGTGTCTGTGTAGTTTTTATGTGCGGATGTTTCCTTTTCCACCATATGCCTAAATATTTCCCAATTTCCACTTGCAGATTC

>CYF54

TACAAAAAGAGTTTCAAAACCGCTCTGTCAAATGATAGGTTGAACTCCCGGAGGTGAATACACACATCACAAAGAGGTTTCTCAGCATGCTTCTGTGTAGTTTTTATGTAAACATATTTCCGTTTCTATCATAGGCCTCAAAGTGCTCCAAATATTCACTTGTACATTC

>CYF55

CTATAAAAAGGAATGTTCAAAATTGCTCAATAAAAATAAAGTTTCAACACCGTGAGATGAGTGCACAAATCACAAAGAAGTTTCTCAAAATGCTTCTGGGTAGTTTTTCTGTGAAGATAGTTCCTTTTCTACCATGGGCCACAAAGGGCTCCAAATACCCACTTGCAGATTC

>CYF56

TAGAAAAAGATTGCTTGGAAACTGCACAATGAAAAGAAAGGTTCAAATATATGAGATGAATGCACACATCACAAAGAAGTTTCTCAGAATCTCTCTGTGTAATTTTTATGTGAAGATATTTCCTTTCCCACCTTAGGTCTTAAAACGCTCCAAATATCCACTTGCAGATAC

>CYF57

CTAGTAACACAGAGTTTCAAAACTGCTCTATCAAAAGATAAGTTCAACTCTGAGAGTTTAGTGCAACCATCGTGAAGAAGTTTCTCAGAATGCTTCTGAGTAGTGTTTATGTGAAGATATTTCCTTTTCCACCATAGGCCTGAAAGCCCTCCAAATATCCACTTGCAGATCC

>CYF58

TACAAAAAGAGTGTTTCAGAACTGCTCAATCAAAAGGAAGGTTCCAGTCTGAGACAAATACACACATCAAAAGGTAGTTTCTCAGAATGCTTCTGTGTAGTTTTTATGTGAAGATATTTTCCTTTCCACCATAGGCCACAAATGGCTCTAAATACCCACTTACATTTTC

>CYF59

CACAAAAAGAGAGTTTCAAAACTGCTCTACCAAAGGTAAGTTTAACGCTGTGAGTTAAGAACATCACAAAGAAGTTTCTCAGAATGCTTCTGTGTAGTTCTTACGTAAAGATATTTCCTTTTACACAATAGGCAGAAAAGTGCTCCAAATATCCACTTGAAGATTC

>CYF60

TACAAAAGTGAGTTTCAAAACTGCTCTATCAAAAGATCAGTTCGTCTCTGTGAGTTGAATGCATACATCAAAAAGAAGCTTCTCAAAATGCTTCTGTGTGGTTTTTCGGTGAAGATAGTTCTTTTTCTACCATAGGTCTCAAACCACTCCAAATATCCACTTGTAGATT

>CYF61

TACAAAAAGAGAGTTTCACAACTGCTCTATCAAACAATATGTTCAACTTTGTGGGTTGAACACAAATATCACAAGAATTTTCTCCCAATGCTTCTGTGTAGTTTTTATGTGAAGACATTTCTTTTCCCTCCATAGTCCACAAAGTGCTCCAAATATCCACTTACATATTC

>CYF62

TACCGAAAGAGTGCTTCCAAACTGCTCAATCAAAAGAGACATTCAAATCTGTGAGGTGAATGCAGACATCGTAAAGAAGTTTCTCAGAATGCTTCTGTGTATTTTTTGTGTGAAGTTATTCGTTTTTGCACCATAGGCCTCCAAGCGTTCTAAATATCCACTTCTAGATTC
