## Additional File 2 for "CENdetectHOR: a comprehensive tool for CENtromere profiling and HOR detection"

>C1F17

AGTATAAGAACTTAAACCGCAACCGCATCTTATAAGCCTAAGTAGTGTTTCCTTGTTAGAAGACACAAAGCCAAAGACTCATATGGACTTTGGCTACACCATGAAAGCTTTGAGAAGCAAGAAGAAGGTTGGTTAGTGTTTTGGAGTCGAATATGACTTGATCTCATGTGTATGATTG

>C1F26

AGTATAAGAAATTAAACCGCAACCGGATCTTAAAGGCGTAAGAAATTTATCCTTGTTAAAAGACACAAAGCCAAAGACTCATATGGACTTTGGCTACACCATGAAAGCTTTGAGAAGCAAGAAGAATGTTGGTTAGTGTTTTGGAGTCGAATATGACTTGATCTCATGTGTATGATTG

>C1F27

AGTATAAGAATTTAAACCGCAACCGGATCTTAAAGGCGTAAGAATTGTACCCTTGTTAAAAGACACAAAGCTAATGACTCATATGGAATGTCTACACCATGAAAGCTTTGAGAAGCAAGAAGAAGGTTGGTTAGTGTTTTGGAGTCGAATGTGACTTGATCTCATGTGTATGATTG

>C1F28

AGTATAAGAACATAAACCGCAACCGGATCTTAAAGGCGTAAGAATTGTATCCTTGTTAGAAGACACAAAGCCAAAGACTCATATGGACTTTGGCTACACCATGAAAGCTTTGAGAAGCAAGAAAAAGTTTGGTTAGTGTTAGTCGAATATGACTTGATGTCATGTGTATGATTG

>C1F29

AGTATAAGAATGTAAACCGCAACCTGATGTTATAAGCCTAAGTAGTGTTTCCTTGTTAGAAGACACAAAGCCAAAGACTCATATGGACTTTGGCTACATCATGAAAGCTTTGAGAAGCAAGAAGAAGGTTGGTTATTGTTTTAGGGTCGAATATGACTTGATGTCATGTGTATGATTG

>C1F2

AGTATAAGAACTTAAACCGCAACCGGATCTTAAAGGCGTAAGAATTGTATCCTTGTTAAAAGACACAAAGCCAAAGACTCATATGGACTTTGGCTACACCATGAAAGCTTTGAGAAGCAAGAAGAAGGTTGGTTAGTGTTTTGGAGTCGAATATGACTTGATCTCATGTGTATGATTG

>C1F30

AGTATAAGAACTTAAACCGCAACCGGATTTTAAAGGCGTAAGAATTGTATCCTTGCTAGAAGACACAAAGCCAAAGACTCATGTGGACTTTGGCTACACCATCAAAGCTTTTAGAAGCAAGAGGAATGTTGGTTAGTGGTTTGGAGTCGAATATGACTTGACCTCATGGGTATGATTG

>C1F32

GAGTATAAGAACTAGAACCGCAACCGGTTCCCGAAAGCTAAAGGAGTATTTCCTTGTTAGAAGATACAAAGCCAAAGACTCATACAGACTTAGCTACACCATCAAAGCTTTGAGAAGCCTTAAGAAGCTTGGTTAGTTTTTTGGAGTCAAATATGACTAGATGTCATGTGTATGATT

>C1F33

GAGTATAAAAACAAGAACCGCAACGGTTCCTAAAAGCTAAAGTAGTGTTTCCTTGTTAGAAGATACAAGCCAAAGACTCATACGGACTTTGGCTACACCATAAAAGCTTTGAGAAGCAATAAGAAGCTTGGTTAGTGTTTTGGAGTCAAATATGACTATGAGTCATTTGTATGATT

>C1F34

AAGTATAAGAACTAGTACCGCAATCGGTTCCCAAAAGTTAAAGTAGAGTTTCATTGGTACAAGATACATAGCCAAAGACTCATACGGACTTCGGCTACACCATCAAAGCtTTGAGAAGCAATAAGAACCTTGGTTAGTGTTTTGGAGTCAAATATGACAAGATGTCATGTGTATGATT

>C1F37

AATGATACACAAAGCATCTAGTCATATTTGACTACAAATCCGCTAACCAAGCTTCTTCTTGATTCTCAAAGCTTTGATGGTGAAGCCGAAGTTCTTATGAGTTTTTGGTTTTGAATCGTATAACAAGGAAGCACTACTTTACTTTTCGGGATCTGTTGAGGTTCTAGTTTTATATTC

>C1F43

GAGTATAAGAACTAGAACCACATCCGGTTGCCAAAAGCTAAAGTAGAGTTTCCTGGTTAAAAGATACAAAGCCAAAGACTCATATGGACTGCGACTACACCATCAAAGCTTTGAGAAGCAATAAGAAGCTTGGTTAGATGTCATGTGTATGAAT

>C1F44

AATCATACACTAACATCTAGATATTTATCCAAACGCTAACAATTTTTCTTTAATATTTTGATCCTAAACACTAAACCTAAACCCTAAACCCTAAATCCCAAACCCTAAAACCTAAACCCCAACTACTAC

>C1F45

CAACCTAAACCTAAACCCAAACCCTAAACCCTAAACCCTAAACTCTTAACCCTAAACCCTTAAACCTAAACCCTAAACCATAGACCCAAACTTTAAAACCTAAATCCTACTTTAGCTTCCGGAATCCGGTTGCGGTTCTAGTTCTTATGCTC

>C1F51

AATCATACACATGACATCTAGTCATATTTCACTCGAAACGCTAACCAAGATTCTTCTTGCTTCTTAAAGTATTATATATATTTGCTCCTAAACACTAAACCTAAACCCTACACCCTAAATCCCAAACCTAAAATCTAACCCTAAACaC

>C1F53

AATCATACACATACATCTAGTCATATTTACTCCGAAACGCTAACCAAGATTCTTCTTGCTTCTTAAAGTATTATATATATTTGCTCCTAAACACTAAACCTAAACCCTACACCCTAAATCCCAAACCTAAAATCTAACCCTAAAcC

>C1F58

AATCATACACATGACATCTAGTCATATTTCAGTCTGAAACGCTAACCCATATTCTTCTTGCTTCTCTAAGTATTATAGTATATTTGATCCTAAACCCTAAACCCTAAACCATATACCCCAAACTTTAAAACCTAAACCCCACTTTAGCTTCCGGAATCCAGCTGCGGTTCTAGTTCTTATACTC

>C2F18

AATCATACACATGACAACAAGTCATATTCGACTCCAAAACACTAACCAACCTTCTTCTTGCTTCTCAAAGCTTTCATGGTGTAGCCAAAGTCCATACGAATGTTTGGCTTTGTGTCTTCTAACAAGGATACAATTCTTACGCCTTTAAGATCCGTTTGCGGTTTAAGTTCTTATACT

>C2F23

AATCATACACATTCCATCAAGTCATATTCGACTCCAAAACACTACCCAAGCTTCTTCTAGCTTCTCAAAGCTTTCATGGTGTGGCCAAAGTCCGTATGACTCTTTGGCTTTGTGTCTTCTAACAAGGAAACACTACTTAGGCTTTTAAGATTGGGTTGCGGTTTATGTTCTTATACTC

>C2F24

CAATCATAAACAAAAAATCAATTCATATTCGACTCCAAAACACTAACCAACCTTCTTTTTGCCTCTCAAAGCTTTCATGGTGTAGCCAAAGTCCATATGAGTCTTTGGCTTTTGTCTTTTAACAAGGAAACACTACTTAGGCTTTTAAGTTCTGGTTGCGGTTTAAATTCTTATACT

>C2F25

CAATCATACAAATGCCATCAAGTCATATTCGACTCAAAAACACTAACAAGGCTTCTTCAAGCTTCTCAAAGCTTTCATAGTGTGGCCAAAGTCGTATGAGTCTTTGGCTTTGTATCTTCTAACAAGGAAACACTACTTAGGCTTTTATGATCGGGTTGCGGTTTAAGTTCTTATACT

>C2F28

CAATCATACACATGACATCAAGTCATATTCGACTCCGAAACACCAACCAACCTTCTTCTTGCTTCTCAAAGCTTTCATGGTGTAGCCAAAGTCCATATGATTCTTTGGCTTTGTGTCTTCTAACAAGGAAACACTACTTAGGCTTTTATGATCGGGTTGCGGTTTAAGTTCTTATACT

>C2F30

CAATCATACACATAACATCAAGTCATATTCGCCTCCAAAACACTAGCCAACCTTTTTCTTGCTTCTCAATGCTTTCATGGTGTAGCCAAAGTCCATATAAGTATTTGGCTTTGTGTCTTCTAACAAGGAAACACTACCTAGGCTTTTAAGATCGGGTTGCGGTTTAAGTTCTTATACT

>C2F32

CAATCATACACATGACATCAAGTCATATTCGATTCCAAAACACTAACCAACCTTCTTCTTGGTTCTAAAAGCTTTCATGGTGTAGCCAAAGTCCATATGAGTCTTTCGCTTTGTGTCTTCTAACATGGATACAATTCTTACGCCTTTAAGATCGGGTTGCGGTTTAAGTTGTTATACTC

>C2F34

CAATCATACACATGACATCAAGTCATATTCAACTCCGAAACACTAACCAACCTTCTTCTTGCTTCTCAAATCTTTCATGGTGTAACCAAAGTCCATATGAGTATTTGGCTTTGTGTCTTCTAACAAGGAAAATCTACTTAGGCTTTTAAAAATTGGGTTGCGGTTTAAGTTGGTAGACT

>C2F35

AATATATTATTTGGTGAAATTAAGTGTTGAGTGCTCTTCACAAGACTAGATTTCATCCCTCAAGGATTTTTCCAAGTTCACCTCTTGGAAAAATATATTCATTTCGGTGAAATCTAAGTGTTGAGTTCGCCTTCACAAGATAGATTTATCTAAATTTTCATtA

>C2F3

CAATCATACACATGACATCAAGTCATATTCGACTCCAAAACACTAACCAACCTTCTTCTTGCTTCTCAAAGCTTTCATGGTGTAGCCAAAGTCCATATGAGTCTTTGGCTTTGTGTCTTCTAACAAGGAAACACTACTTAGGCTTTTAAGATCGGGTTGCGGTTTAAGTTCTTATACT

>C3F10

CAATCATACACATGACATCAAGTCATATTCGACTCCAAAACACTAACCCACCTTCTTCTTGCTTCTCAAAGCTTTCATGGTGTAGCCAATGTCCATATGAGTCTTCGGCTTTGTGTCTTCTAACAAGGATACAATTCTTACTCCTTTAATATCGGGTTGCGGTTTAAATTCTTATACT

>C3F11

CAATCATACACATGACATCAAGTCATATTCGACTCCAAAACACTAACCAACCTTCTTCTTGCTTCTTAAAGCTTTCATGGTGTAGCCAAAGTCCATATGAGTCTTTGGCTTTGTGTTTTCTAGCAAGGATACAATTCTTACGCCTTTAAGATCCGATTGTGGTTTAAGTTCTTATACT

>C3F12

AATATATACATACATCAAGTCATATTCGACTTTAAAACCTAACCAACCTTCTTCTTGCTTCTCAAAGCTTTCATGGTGTACCAAGTCCATATGAGTCTTTGGCTTTATGTCTTCTAACAAGGAAAACTCTTAGCTTACAAGATCGGTTTGGTTTAAGTTTTATACT

>C3F13

TATAAACATAAACTATTCGATTAAACCTAATAGTTTCTTGTTACCAAAGCAAATTCATATGATTGTAGCTTTGAGAACAAGAAAaTTCTTAGTTTTAATCGAATGATGTTTCATGTTTACT

>C3F14

AGTATAAGAACTTAAACCGCAACCCGATCTTGTAAGCCTAAGTAGTGTTCCTTGTTAGAAGACACAAAGCCAAAGACTCATATGGACTTTGGCTACACCATGAAAGCTTTGAGAAGCAAGAAGAAGGTTGGTTAGTGTTTTAAGTCGAATATGACTTGATGTCATGTGTATATTG

>C3F19

CAATCATACACATGACATCAATTCATATTCGACTTTAAAACACTAACCAACCTTCTTCTTGCTTCTCAAAGCTTTCATGGTGTACCCAAAGTCCATATGAGTCTTTGGCTTTATGTCTTCTAACAAGGAAACACTACTTAGGCTTACAAGATCGGGTTGCGGTTTAAGTTCTTATACT

>C3F20

CAATGATATACATGACATCAAGTCATATTCGACTCCAAAAAACCTAACCAACCTTCTTATTGCTTCTCAAAGCTTTCATGGTGTAGCCAATGTCCATATGAGTCTTTGGCTTTGTGTCTTCTAACAAGGATACACTTCTTAGGCTTACAAGATCGGGTTGCGGTTTAAGTTGTTATACT

>C3F22

CAATGATATACATGACATCAAGTCATATTCGACTCCAAAACACTAACCAACCTTCTTCTTGCTTCTCAAAGCTTTCATGGTGTAGCCAATGTCCATATGAGTCTTTGGCTTTGTGTCTTCTAACAAGGATACACTTCTTAGGCTTACAAGAGCGGGTTGCGGTTTAAGTTGTTATACT

>C3F25

CAAACATACACATTACATCAAGTCATATCCGACTCCAAAACATTAACCAACCTTCTTCTTACTTCTCAAAGCTTTCATGGTGTAGCCATTGTCCATATGAGTCTTTGGCTTTGTGTCTTCTAACAAGGATACACTACTTAGGCTTACAAGATCGGGTGTTGTTTAAGTTCTTATACT

>C3F26

CAATCATACACATGACATGAAGTCATATTCGACTCCAAAACTCCAACCAACCTTCTTCTTGCTTCTCAACGCTTTCATGGTGTAGCCAATGTCCATATGAGTCTTTAGCTTTGTGTCTTCTAACAAGGATACAATTCTTACGCCATTAAGATCCGGTTGCGGTTTAAGTTCTTATACT

>C3F27

CAATCATACACATGACATCAAGTCATATTCGACTCAAAAACACTAACCAACCTTCTGCTTTCTTCTCATATCTTTCATGGTGTAGCCAAAGTCCATATGAGTCTTTGGCTTTGTGTCTTCTAACAAGGATACAATTCTTACGCCTTTAAGATCCGGTTGCGGTTTAAGTACATATACTC

>C3F2

CAATCATACACATGACATCAAGTCATATTCGACTCCAAAACACTAACCAACCTTCTTCTTGCTTCTCAAAGCTTTCATGGTGTAGCCAAAGTCCATATGAGTCTTTGGCTTTGTGTCTTCTAACAAGGATACAATTCTTACGCCTTTAAGATCCGGTTGCGGTTTAAGTTCTTATACT

>C3F8

CAATCATACACATGACATCAAGTCATATTCGACTCCAAAACACTAACCAAACTTCTTCTTGCTTCTCAAAGCCTTTCCTGGTCTAGCCAAAGTCCATATGAGTCTTTGGCTTTGTGTCTTCTAACAAGGATACAATTCTTACGCCTTTAAGATCCCGTTGCGGTTTAAGTTGTTATACT

>C3F9

AGTATAAGAACTTAAACTGCAACCAGATCTTAAAGGCGTAAGATTTGTATCCTTGTTAGAAGACACAAATCCAAATACTCATATGGAATTTGGCTACACCATGAAAGCTTTGAGAAGCAAGAAGAAGGTTGGTTAGTGTTTTGGAGTCGAATATGACTTGATGTCATGTGTATGATTG

>C4F10

GAGTATAAAAACTAGAACCGCAACTGGTTCCCAAAAGCAAAAGTAGTGTTTCCTTGTTAGAAGATACAAAGCCAAAGACTCATACGGACTTCGGCTACACCATGAAAATTTTGAGAAGCAATAAGAAGCTTGGTTAGTGTTTTGGAGTCAAATATGACTAGATGTCATGTGTATGATT

>C4F13

GAGTATAAGAACTAGAATGGCAACCGATTCCCAATAGCTAAAGTTATGTTTCCTTGTTAGAAGATACAAAGCCAAAGACTCATACGGACTTCGTGTACACCATCAAAATTTTGAGAAGCAATAAGAAGCTTGGTTAGTATTTTAGAGTCAAATATTACAGGATATCATGTGTATCATT

>C4F16

GAGTATAAGAACTAGAACTGCAACTGGTTCCCAAAAGCTAAAGTAATGTTTCCTAGTTAGAAGATACAAAGCCAAAGACTCATACGGACTTTGGCTACACCATCAAAGTTTTGAGAAGCAATAAGTAGCTTGGTTAGTATTTTGGTGTCAAATAAGACAAGATGTTATGTGTATGATT

>C4F17

GAGTATAAGAACTAGAACCGCAACCGTTTCCCAAAAGCTAAAGTAGTGTTTCCTTGTTAGAAGCTACAAAGCCAAAGACTCATACGGACTTCGGCTACACCATCAAAGTTTTGAGAAGCAATAAGAAGCTTGGTTAGTATTTTGGAGTCAAATAAGACAAGATGTTATGTGTTTGATT

>C4F19

GAGTATAAGAACTAGAACCGCAACCGATTCCCAAAAGCTAAAGTAATGTTTCTTGTTTGTAGATACAAAGCCAAAGACTCATACCGACTTCGGCTACACCATCAAAGCTTTGTGAGGCAATAAGAAGCTTGGTTAGTTTTTCGGAATCAAATATGGCTAGATGTCATGTGTATGATT

>C4F21

GAGTATAAGAACTAGAACCGCAACCGATTCCCAAAAGCTAAAGTAATGTTTCCTTGTTTGAAGAtACAAAGTCAAAGACTCATACGGACTTCGGCTACACCATCAAAACTTTGTGAGGCAATAAGAAGCTTCGTTAGTTTTTTGGAGTCAAATATGACTAGATGTCATGTGTATGATT

>C4F23

GAGTATAAGAACTAGAACCGCAACTAATTCCCAAAAGCTAAATTAATGTTTCCTTGTTAGAAGATACAAAGCTAAAAACTCATACGGACTTCGGCTAAGCCATTAATGTTTTGAGAAGCAATAAGAAGCTTGGTTATTTTTTTGGAGTCAAATATGACTAGATGTCATGTGTATGATT

>C4F26

GAGTATAAGAACTAGAACCGCAACCGTTTCCCAAAAGCTAAAGTAGTGTTTCCTTGTTAGAACATACAAAGCAAAAGAATCATATGGACTTCAGCTACACCATCAAAGCTTTGAGGAACAATAAGAAAAGCTTTGTTATGGTTTTGGAGTCACATATGACTAGAGGTCATGTGGATGATT

>C4F27

GAGTATAAGAACTAGAACCGCAACATCTTCCCAAAAGCTAAAGTTGTGTTTCCCTTCTAGAAGATACAAAGCCAAAAACTCATACGGACTTTGGCTACACCATCAAAGCTTTAAGAAGCAACAAGAAGTTTGGTTAGTGTTTTGGAGTCAAAAAAGACTATATGTCATGTGTAGTATT

>C4F31

AATCATACACATGACATCAAGTCATATTCGACTCCAAAACACTAACCAACCTTCTTCTTGCTTCTCAAAGCTTTCATGGTGTAGCCAAAGTCCTATGAGTCTTTGGCTTTGTTCTTCTAACAAGGAAACACTACTTAGGCTTATAAGATCGGGTTGCGGTTTAAGTTCTTATACT

>C4F32

AATCATACACATGACATCAAGTCATATTCGACTCCAAAACACTAACCAACCTTCTTCTTGCTTCTCAAAGCTTTCATGGTGTAGCCAAAGTCCTATGAGTCTTTGGCTTTGTTCTTCTAACAAGGAAACACTACTTAGGCTTATAAGATCGGGTTGCGGTTTAAGTTCTTATACTC

>C4F34

CAATCATACACATGACATCAAGTCATATTTGACTCCAAAACACTAACCAACCTTCTTCTTGCTTCTCAAAGCTTTCATGGTTAGCCAAAGTCCATATGAGTCTTTGGCTTTGTGTCTTCTAACAAGGAAAGACTACTTAGGCTTTTAAGATCGGGTTGCGGTTTAAGTTGTTATACTC

>C4F37

AATCATACACATGACATCAAGTCATATTCGACTCCAAAACACTAACCAACCTTCTTCTAGCTTCTCAAAGCTTTCATGGTGTAGCCAAAGTCCGTATGAGTCTTTGGCTTTGTATGTTCTAACAAGGAAACACTACTTAGGCTTATAAGATCGGGATGCGGTTTAAGTTCTTATACTC

>C4F40

CAATCATACACATGACATCAAGTCATATTCGTCTCCAAAACACTAACCAACCTTCTTCTTGCTTTTCAAAGCTTTCATGGTTTAGCCAAAGTCCCTATGAGTCTTTGGCTTTGTTTCTTCTAACAAGGAAACACTACTTAGGCTTTTATGATCGGGTTGCGGTTTAAGTTGTTATACTC

>C4F42

AATCATACACATGACATCAAGTCATATTTGACTCCAAAACACTAACCAACCTTCTCTTTGCCTTTCAAAGCTTTCATGGTGTAGCCAAAGTCCATATGAGTCTTTGGCTTTGTGTCTTCTAACAAGGAAACACTACTTAGGCTTTTATGATCGGGTTGCGGTTTAAGTTGTTATACTC

>C4F43

cAATCATACACATGACATCAAGTCATATTTGACTCCAAAACACTAACCAACCTTCTCTTTGCCTTTCAACGCTTTCATGGTGTAGCCAAAGTCCATATGAGTCTTTGGCTTTGTGTCTTCTAACAAGGAAAGACTACTTAGGCTTTTAAGATCAGGATGCGGTTTAAGTTCTTATACTc

>C4F44

CAATCATACACATGACATCAAGTCATATTCGACTCCAAAACACTAACCAACCTTCTTTTGCTTCTCAAAGCTTTCATGGTGTAGCCAAAGTCCATATGAGTCTTTGGCTTTGTGTCTTCTAACAAGGATACAGTTCTTACGCCTTTAAGATCCGGTTGCGGTTTAAGTTGTTATACT

>C4F45

CAATCATACACATGACATCAAGTCATATTCGACTCCAAAACACTAACAACCCTTCTTCTTGCTTCTCAAAGCTTTCATGGTTTAGCCTCAGTCTATATGAGTCTTTGGTTTTGTGTCTTCTAACAAGGATACAATTCTTACGCCTCTAAGATCCGGTTGCGGTTTAAGTTGTTATACTC

>C4F48

AATCATACACATGACATCAAGTCATATTCGACTCCAAAACACTAACCAAGCTTCTTCTTGCTTCTCAAAGCTTTGATGGTTTAGCCAAAGTCCATATGAGTCTTTGTCTTTGTATCTTCTAACAAGGAAACACTACTTAGGCTTTTAGGATAAGTTGCGGTTTAAGTTCTTATACTC

>C4F4

AATCATACACATGACATCAAGTCATATTCGACTCCAAAACACTAACCAACCTTCTTCTTGCTTCTCAAAGCTTTCATGGTTTAGCCAAAGTCCATATGAGTCTTTGGCTTTGTATCTTCTAACAAGGAAACACTACTTAGGCTTTTAAGATCGGTTGCGGTTTAAGTTCTTATACTC

>C4F51

AATCATACACATGACATCAAGTCATATTCGACCCCAAAACAATAACCAAGCTTCTTCTTGCTTCTCAAAGCTTTGATGGTTTAGCCGAAGTCCATATGAGTCTTTGTCTTTGTATCTTCTAACAAGGAAACACTACTTAGGCTTTTAGGATAAGATTGCGGATTAAGTTCTCATACTC

>C4F52

AATCATACACATGACATCAAGTTATATTCGACTCCAAAACACTAACTAAGCTTCTTCTTGCTTCTCAAAGCTTTGATGATTTAGCCGAAGTCCATATGAGTCTTTGTCTTTGTATCTTCGAACAAGGAAACACTACTTAGGCTTTTAGGATAAGATTGCGGTTTAAGTTCTTATACTC

>C4F53

AATCATACACATGACATCAAGTCATATTCGACTTTAAAACACTAACCAAGCTTCTTCTTGCTTCTAAAAGTTTAGATGGTTTAGCCGAAGTCCATATGAGTCTTTATCTTTGTATCTTCTAACAAGGAAACACTACTTAGGCTTTTAGGATAAGGTTGCGGTTTAAGTTCTTATACTC

>C4F55

ATACATACACATGACATCAAGTCATATTCGACTCCAAAAAACTAACCAAGCTTCTTCATGTTTCTCAAAGCTTTAATGGTTTAGCCGAAGTCCATATGAGTCTTTGTCTTTGTATCTTCTAACAAGGAAACACTACTTAGGCCTTTAGGATAAGATTGTGATTTAAGTTCTTATACTC

>C4F56

AATCATACACATGACATCAAGTCATATTCGACTCCAAAACACTAATCAAGCTTCTTCTTGCTTCTCAAACCTTTGATGGTTTAGCCGAAGTCCATATGAGTCTTTATCTTTGTATCTTCTAACAAGGACACACTACTTAGGCTTTTAGGATATGGTTGCTGTTTAAGTTCTTATACTC

>C4F57

ATTCATACACATGACATCAAGTCATATTCGACTCCAAAATACTAATGAAGCTTCTTCTTGCTTCTCAAAGCTTTGATGATTTAGCCGAAGTCCATATGAGTCTTTGTCTTTGTATCTTCTAACAAGGAAACACTACTTAGGCTTTTAGGATAAGATTGCGGTTTAAGTTCTTATACTC

>C4F58

AATCATACACATGACATCAAGTTATATTtGACTACAAAATACTAACCAAGCTTCTTCTTCCTTCTCAAAGCTTTGATGGTTTAGCCAAAGTCCATATGACTCTTTGTCTTTGTATCTTCTAACAAGGAAACACTACTTAGGCGTTTAGGATAAGATTGTGATTTAAGTTCTTATACTC

>C4F60

AATCATACACATGACATCAAGTCATATTCGACTCCAAAACACTAACCAAGCTACTTCTTGCTTCTCAAAGTTTTGATGGATTAGCCGAAGTTCATATGAGTCTTTATCTTTGTATCTTCTAACAAGGAAACACTACTTAGGCTCTTAGTATAAGATTGCGGTTTAAGTTCTTATACTC

>C4F61

AATCATACACATGACATCAAGTCATATTCGACTCCAAAACACTAACAAAGCTTCTTCTTGCTTCTCAAAGCTTCTTCTTGCTTCTCAAAGCTTTGATGGTTTAGCCAAAGTCCATATGAGTTTTTATCTTTGTATCTTCTAACAATGAAACACTACTTAGGCCTTTAGGATAAGGTTGCGGTTTAAGTTCTTATACTC

>C4F62

AATCATACAAATGACATCTAtTCATATTTGACTCCAAAACACTAACCAAGCTTCTTATTGCTTCTCAAAGCTTTGATGGTGTAGCCGAACTCTGTATGAGTCTTTTGCTTTGTATCTTCTAACAAGGAGATACTACTTAGGCTTTCAAGATCCAGTTGAGATTCTAGTTCTTATACTC

>C4F63

tGGACAAAATGGGTATAAGTGTTGTCTAAACACTCCTAATCCATCTCTAACTCTTATAATTAGTCAAATGCATTGGATTGTGACACATTTTGACCATAGAAACACTAACAAAGCTATTTACTGCTTCTAAGCAATgTTTTGTTGGTTTTAGCCTCTtTTG

>C4F64

tAgTCAAATGCATTGGATTGTGACACATTTTGACCATAGAAACACTAACAAAGCTATTGACTGCTTCTAAGCAATTTTTTGTTGGTTTTAGCCTCTTTTGGGAGAAAATGGGTATAAGTGTTGTCTAAACACTCCTAATCCATCTCTAACTCTTATAATT

>C4F7

GAGTATAAGAACTAGAACCGCAACCGATTCCCAAAAGCTAAAGTAATGTTTCCTTGTTAGAAGATACAAAGCCAAAGACTCATACGGACTTCGGCTACACCATCAAAGTTTTGAGAAGCAATAAGAAGCTTGGTTAGTTTTTGGAGTCAAATATGACTAGATGTCATGTGTATGATT

>C5F15

AGTATAAAACTTAAACCGCAACCCGATCTTAAAAGCCTAAGTAGTGTATCCTTGTTAGAAGACACAAAGCCAAAGACTCATATGGACTTTGGCTACACCATGAAAGCTTTGAGAAGCAAGAAGAAGGTTGGTTAGTGTTTTGGAGTCGAATATGACTTGATGTCATGTGTATGATTG

>C5F18

AGTATAACAACTTAAACCGCAACCGATCTTAAAAGCCTAAGTAGTGTATCCTTGTTAGAAGACACAAAGCCAAAGACTCATATGGACTTTGGCTACACCATGAAAGCTTTGAGAAGCAAGAAGAAGGTTGGTTAGTGTTTTGGAGTCGAATATGACTTGATGTCATGTGTATGATTG

>C5F20

AGTATAAGAACTTAAACCGCAACCGGATCTTAAAGGCGTAAGAATTGTATCCTTGTTAGAAGACACAAAGCCAAAGACTCATATGGACTTTGGCTACACCATGAAAGCTTTGAGAAGCAAGAAGAAGGTTGGTTAGTGTTTTGGAGTCGAATATGACTTGATGTCATGTGTATGATTG

>C5F21

AGTATAAGAACTTAAACCGCAACCGGATCTTAAAGGCGTAAGAATTGTATCCTTGTTAGAAGACACAAAGCCAAAGACTCATATGGACTTTGGCTACACCATGAAAGCTTTGAGAAGCAAGAAGAAGGTTGGTTAGTGTTTTGGAGTCGAATATGACTTGTGTCATGTGTATGATTG

>C5F27

AGTATAAGAACTTAAACCGCAACCCGAATTTAAAGGCGTAAGAATTGTATCCTTGTTAGAAGACACAAAGCCAAAGACTCATACGGACTTTGCCTACACCATGAAAGCTTTGAGAAGCAAGAAGAAGGTTGGTTACTGTTTTGGAGTCAATATGACTTGATGTCATGTGTATGATTG

>C5F29

AGTATAACAACTTAAACCGCAACTCGATCTTAAAAGCCTAAGTAGTGTTTCCTTGTTAGAAGACACAAAGCCAAAGACTCATATGGACTTTGGCTACACCATGAAAGCTTTGAGAAGCAAGAAGAAGGTTGGTTAGTGTTTTGGAGTCGAATATGACTTGATGTCATGTGTATGATTG

>C5F3

AGTATAAAACTTAAACCGCAACCGATCTTAAAAGCCTAAGTAGTGTATCCTTGTTAGAAGACACAAAGCCAAAGACTCATATGGACTTTGGCTACACCATGAAAGCTTTGAGAAGCAAGAAGAAGGTTGGTTAGTGTTTTGGAGTCGAATATGACTTGATGTCATGTGTATGATTG

>C5F53

AGTATAAGAACTTAAACCGCAACACGATCTTATAAGCCTAAGTAGTGTTTTCTTGTTAGAAGATACAAAGCCAAACACTCATACAGACTTTGGCTACACCATGAAAGCTTATAGAAGCTAGAAGAAGGTTGGTTAGTGTTTCGGAGTCGAATATGACTTGATGTCATGTGTATGATTG

>C5F58

CGTATAAGAACTTAAACCACAACCCGGTCTTAAAAGCCTAAGTAGTGTTTCCTTGTTAGAAGATACAAAGCCAAAGAATCATACGGACTTTGGCTACACCATGAAAGCTTTGAGAAGCTAGAAGAAGCTTGGTTAGTGTTTTAAAGTCTAATATGACTTGATGTCATGTGTATGATTG

>C5F59

AGTATAAGAACTTAAACCGCAACCCGATCTTAAAAGCCTAAGTGTTGTTTCCTTGTTAGAAGATACAAAGCCAAAGACTCATACAGACTTCAGCTACACCATGAAAGCTTTGAGAAGCTAGAAGAAGCTTGGTTAGTGTTTTGGAGTGGAATATGACTTGATATCATGTGTATGATTG

>C5F5

GAGTATAAAAACTAGAACCGCAACGGATCTTAAAAGCCTAAGTATTGTATCCTTGTTAGAAGATACAAAGACAAAGACTCATACGGACTTCGGCTACACCATCAAAGCTTTGAGAAGCAAGAAGAAGCTTGGTTAGTGTTTTGGAGTCGAATATGACTTGATGTCATGTGTATGAGT

>C5F60

AGTATAAGAACTTAAACCGCAACCGGATCTTAAAGGCGTAAGAATTGTATCCTTATTAGAACAACAAAAGCAAAGACTCATACGGACTTTGGCTACACCATGAAAGATATGAGAAGCAAGAAGAAGGTTGGTTAGTGTTTTGGAGTCGAAAATAACTTGATGTCATGTGTATGATTG

>C5F61

AGTATAACAACTTAAACCGAAAATCGATCTTAAAAGCCTAAATAGTGTATCCTTGTTAGAAGACACAAAGCCAAAGACTCATATGGACTTTGGCTACACCATGAAAGCTTTGAGAAGCAAGAAGAAGTTTGGTTAGTGTTTTTGAGTCAAATATGATTTATTGTTATGTGTATAATTG

>C5F63

AGTATAAGACTTAAATCGCAACCGGATCTTAAAGGCGTAAGAATTGTACCGTTGTTAGAAGACACAAAGCCAAAGACTCATATAGACTTTGGCTACACCATGAAAGCTTTGAGAAGCAAGAAGAAGGTTGTTTTGTTTTTTGGAGTTGAATATGACTTGATGTCTTGTGTATTATTG

>C5F64

AGTATAACAACTTAAACCCCAACTCGATCTTAAAAGCCTAAGTAGTGTATCCTTGTAAGAAGATACAAAGCCAAAGACCCATACGGACTTTGGCTACACCATGAAAGCTTTGAGAAGCTAGAAGAAGGTTGGTTAGTGTTTTGGAGTCGAATGTGACTTGATGTCATGTGTATGATTG

>C5F65

AGTATAACAACTTTAACCGCAACCAGATCTTAAAAGCCTAATTAGTGTTTTCTTGTTAGAAGACCAAAGCCAAAGACTCATATGGACTTTGGCTAAACCATGAAAGCTTTGAGAAGCAAGAAGAAGGTTGGTTAGATTTTTCGAGTCGAATATGACTTGATGTCATGTGTATGATTG

>C5F68

AGTATAACAATTTAACTGCAACCCGATCTTAAAAGCCTAAGTAGTGTTTCTTTGTTAGAAGATACAAAGCCAAAGACTCATACGGACTTTGACTACACCATGAAAGCTTTGAGAAGCAAGAAGAAGGTTGGTTAGTGTTTTGGAGTCGAATATGACTTGTTGTCATGTGTATGATTG

>C5F70

AGTATAAGAACTTAAACACAACATAATCTTATAAGCCTAAGTAGTGTTTCCTTGTTAGAAGATACAAAGCCAAAGACTCATACGGACTTTGGCTACACCATGAAAGCTTTGAGAAGCTAGAAGAAGGTTGGTTAGTGTTTCGGAGTCGAATATGACTTGATGTGATGTGTATGATT

>C5F71

AGTATAAGACTTAAACCGCAACTGGATCTTAAAGGCGTAAGAATTGTATCGTTTTTAGAAGACACAAAGCCAAAGACTCATATGGACTTTGGCTACACCATGAAAGCTTTAAGAAGCATTAAGAAGGTTGGTTAGTGTTTTGAAGTTGAATATGACTTGATGTCATGTATATTATTG

>C5F72

AGTATAAGAACTTAAACCGCAACACGATCTTATAGTCTAAGTAGTGTTTCTTTATTAGAAGAAACAAAGCCAAAGACTCATATGGACTTTGGATACACCATGAAAGCTTTGAGAAGTAACAAGAAGGTTGGTTAGTGTTTTGGAGTCGAATATGACTTGATGTCATGTGTATGATTG

>C5F73

AGTATAAGACTTAAACCGCAACAGATCTTAAAGGCGTAAGAATTGTATCTTTGTTAGAAGACACAAAACCAAAGACTCATATGGACTTTGGCTACACCATGAAATCTTAGAGAAGCAAGAAGAAGGTCTATTAGTGTTTTGGAATCCAATATGACTTGATGTCATGTGTATGATTG

>C5F74

CAaTCATACACATGACATCAAgTCATATTCGACTCcAAAACACTAACAACTTCTCaAAcTTTAtGGTGTAGCCAAAGTCCgTATGAGTCTTTGGCTTtGTaTaTTTAACATgGAaACAtTaCTTAgGCTTTAGAtCCGTTGTGATTTAGTTCTATACT

>C5F78

GAGTATGACAACTAGAACCATAACCGGATCTTAAAAACCTAAGTATTGAATCTTTGTTAGAAGATACAAAGACAAAGACTCATACGGACTTCGACTACACTATCAAAGCTTTGAGAAGCAAGAAGAAGCTTGGTTAGTGTTTTGGATTCGAATATGACTTGATGTCATGTGTATGACT

>C5F83

GAGTATAAAGACTATAACCGCTACGGATCTTAAAAGCCTAAGTATTGTATCCTTGTTAGAAGATACAAAGACAAATACTCATATGCACTTCGGCTAAACCATCAAAGCTTTGAGAAGCAAGAAGAAGCTTGGTTAGTGTTTTTGAGTGAAATATGACTAGATGTCATGTGTATGAGT

>C5F84

AGTATAAGACTTAAAACCGCAAACGGATCTTAAAGGCGTAAGAATTGTATCCTTGTTAGAAGACACAAAGCCAAAGACTCATATGAAAGCTTTGAGAAGCAAGAAGAAGGTTGGTTAGTGTTTTGGAGTCGAATATGACTGGATGTCATGTGTATGATTG

>C5F85

aAATTCTAACAAAAGATCCGCGTAGCAATTGTCTAAAGCAAACATTCTAGTTTAATACTGGATCAGAATTCCGGGTTCGAAGCCCGGCAGCGGAAAAATCCTTTTGGTGCTTTTTATATGTATCTTGTATGACAATTGAAGATACTTTTTCATATTGTTCTTTCTTCTCTGCTCATTGTTAT
