## Supplementary figures and images for "CENdetectHOR: a comprehensive tool for CENtromere profiling and HOR detection"

### Supplementary Figure 1

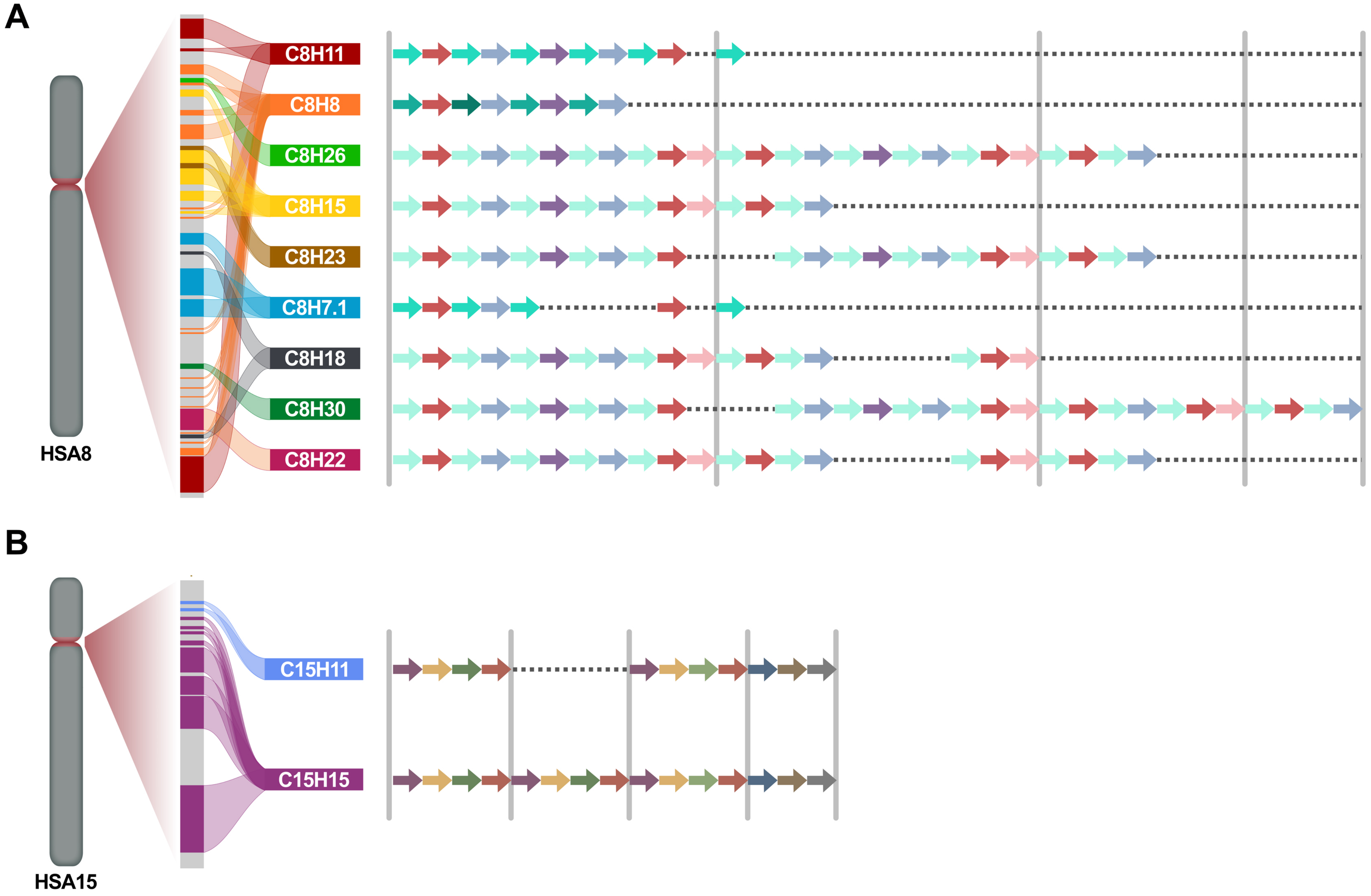

### Supplementary Figure 2

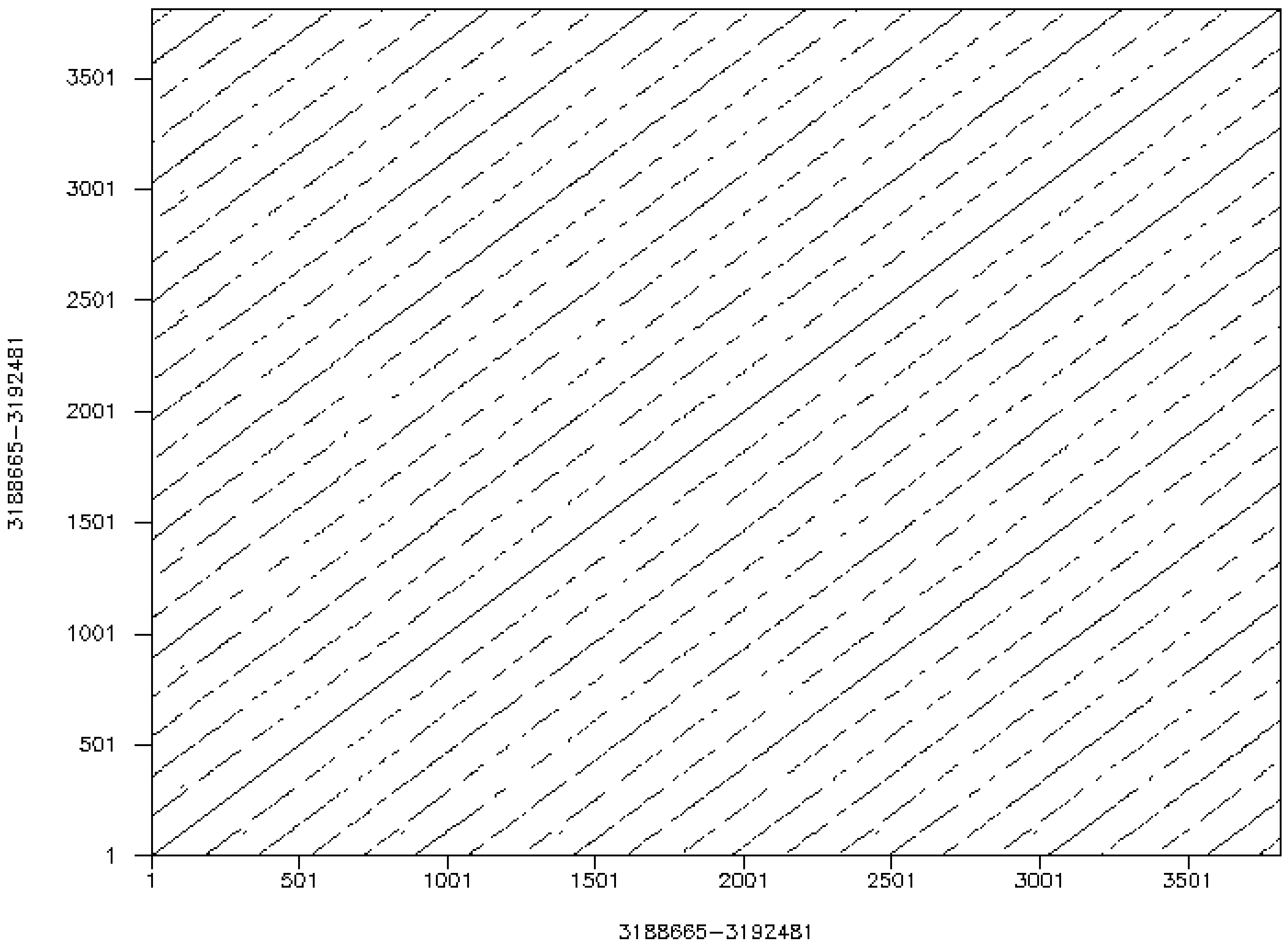

### Supplementary Figure 3

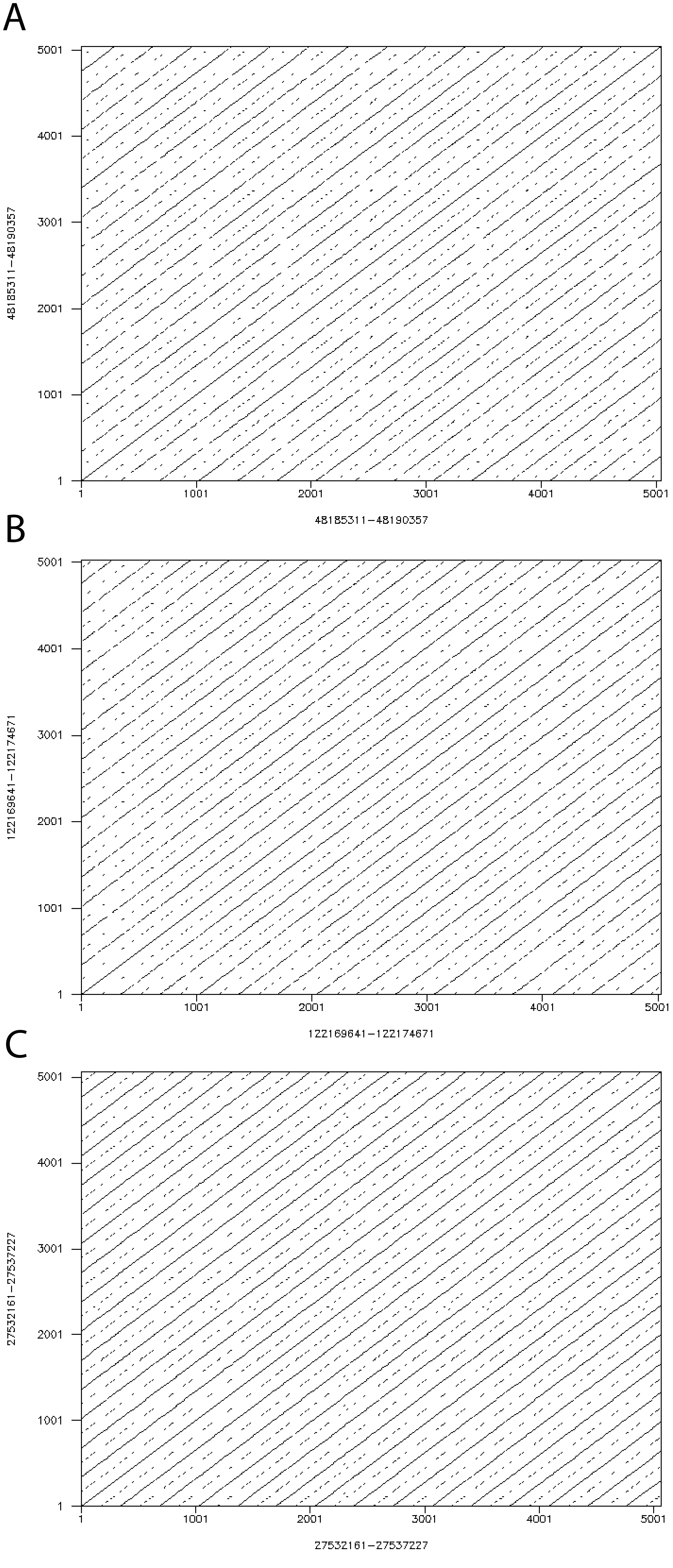
