## Supplementary information for "CENdetectHOR: a comprehensive tool for CENtromere profiling and HOR detection"

Operational definitions

*Independent set of clades* definition.

Given a phylogenetic tree T, a set C of the clades is said to be *independent* if no clade of C is nested in another clade of C.

*Clade labelling* definition.

Given a set of identified monomers, M, a set of sequences S = {s₁, ..., sₙ} defined over the alphabet of monomers M, a phylogenetic tree T with the monomers of M as its leaves, and a maximal independent set of clades C, a clade labelling of S with C is a set of sequences S' = {s'₁, ..., s'ₙ}, where each sequence s'ᵢ is derived from sᵢ by replacing each monomer identifier with the corresponding clade from C.

*HOR* definition.

Given a set of identified monomers M, a set of sequences S = {s₁, ..., sₙ} defined over the alphabet of monomers M, a phylogenetic tree T with the monomers of M as its leaves, and a positive integer k, a higher-order repeat (HOR) on S, T, and k is defined as a sequence h over an independent set of clades C from T, such that the clade labelling of S with C contains at least two instances of a consecutive sequence where h is repeated k times. The set of maximal consecutive sequences where h is repeated at least k times is referred to as the coverage of a HOR.

*HOR* non-redundancy properties.

Given a set of DNA sequences split into monomers and a phylogenetic tree of these monomers, many HORs can be identified. A HOR is non-redundant (hence, considered) if at least one of the following properties is verified:

**broader coverage**, i.e. the region covered by the HOR is not a sub-region (or equal) of a region already covered by another HOR;

**longer repeat period**, i.e. the length of the repeated sequence of clades is greater than that of other HORs covering the same region (or a region encompassing it);

**greater intra-HOR diversity**, i.e. the number of distinct clades within the sequence is higher than the one in other HORs covering the same region (or a region encompassing it);

**greater generality**, i.e. it is more generic (one or more clades higher in the phylogenetic tree) than all other HORs covering the same region or subregions, while maintaining the same period and diversity;

**greater inter-HOR diversity**, i.e. it belongs to a set of two or more non-overlapping HORs that collectively cover a region otherwise covered by only a single HOR.

Evolutionary analysis of the monomer families

The evolutionary analysis of the families composing HOR variants in chromosome 10 was inferred in MEGA11 [1] using the Minimum Evolution method [2]. The tree is drawn to scale, with branch lengths in the same units as those of the evolutionary distances used to infer the phylogenetic tree. The evolutionary distances were computed using the Maximum Composite Likelihood method [3] and are in the units of the number of base substitutions per site. The ME tree was searched using the Close-Neighbor-Interchange (CNI) algorithm [4] at a search level of 1. The Neighbor-joining algorithm [5] was used to generate the initial tree. All ambiguous positions were removed for each sequence pair (pairwise deletion option).
